# Genetic and behavioral analyses suggest that larval and adult stages of *Lucilia cuprina* employ different sensory systems to detect rotten beef

**DOI:** 10.1101/2024.12.20.629795

**Authors:** Juan P. Wulff, Rachel K. Laminack, Maxwell J. Scott

## Abstract

**Background:** The blowfly *Lucilia cuprina* is a destructive parasite of sheep that causes flystrike or myiasis. Larvae consume the animal’s living flesh, producing large wounds that can lead to death. Growing resistance to conventional control methods has prompted the analysis of alternative strategies.

**Methods:** An RNA-Seq analysis was used to identify sensory receptors and other genes relevant to the physiology of *L. cuprina* larvae. Adult females and larvae of the same species carrying a loss-of-function mutation for the *L. cuprina* odorant coreceptor gene (*LcupOrco*) were obtained by gene editing. Their response to fresh and rotten meat at different temperatures was evaluated.

**Results:** The RNA-Seq analysis of whole larvae at different stages and third instar head and gut tissues, suggested that odorant (OR), gustatory, ionotropic and *pickpocke*t receptors may not play a central role in the *L. cuprina* larval sensory signaling and digestive systems. Rather, ATP-binding cassettes (ABCs) were highly enriched in head and gut RNA, and odorant-binding proteins (OBPs) only in the head. To confirm that ORs are not essential for larval detection of rotten beef, diet-choice assays were performed including larvae and adults homozygous for a null mutation in *LcupOrco*. While the attraction of adult females to rotten beef was fully disrupted, *LcupOrco* mutant larvae showed no change in diet preference.

**Conclusions:** The expression pattern of the ABC and OBP gene families suggests a central role in the sensory system of the *L. cuprina* larva for these receptors. Behavioral assays showed that ORs are essential for the adult female response to rotten beef, but not for larval behavior. These findings are consistent with high levels of expression of *LcupOrco* in the adult female antenna but very low expression in larvae.

## Background

Immature stages of blowfly species (Diptera: Calliphoridae) display a wide spectrum of feeding specializations [1]. Based on their feeding habits, species within this family can be classified from obligate parasites of live animals to obligate necrophagous (dead tissue consumers), with many facultative forms in between [1–3]. Some of these species are considered livestock pests, such as *Lucilia cuprina* (Wiedemann, 1830), *Lucilia sericata* (Meigen, 1826) and *Cochliomyia hominivorax* Coquerel, 1858 [2,4]. *Lucilia cuprina* is a facultative ectoparasite and one of the main causative agents of flystrike in sheep in Oceania [5,6]. Conventional insecticides constitute the current control method for this pest [7], however, an increasing resistance to organophosphates and benzoylphenyl ureas [7], have motivated the development of alternative genetic strategies for population control, such as the sterile insect technique (SIT) and female specific release of insects carrying a dominant female lethal gene (fsRIDL) [8–10].

Flystrikes on live animals are associated with the female oviposition behavior in blowfly parasitic species [2]. *Lucilia cuprina* females lay eggs in open wounds or moist tissues of host animals, clustering them in masses of ∼200 eggs per ovarian cycle [11,12]. After egg hatching, the immature larval stages congregate in clusters of maggots called myiasis [12]. Within the myiasis the larvae complete the three larval (L) stages, from L1 to L3 [12]. This maggot arrangement favors the action of the larval secretions and excretions (SE) improving larval feeding and survival [13]. *Lucilia cuprina* larval SE includes several components such as gut microbiota, lipases, peptidases and ammonia [14,15]. Peptidases and lipases together with the gut microbiota facilitate host-tissue ingestion and digestion [15]. Ammonia, along with other SE, has been shown to control deleterious fungi and bacteria that impair larval survival [16]. Furthermore, maggot masses produce heat, reaching up to 18.7°C above ambient temperature depending on the number of larvae [17]. Temperature is a key factor driving *L. cuprina* larval development and survival, and larvae are constantly moving in and out of the maggot mass to avoid overheating [18]. The highest survival rate for this species, considering the whole life cycle, was observed at 24±1°C [19,20]. Temperatures below and above this range can be deleterious to *L. cuprina* decreasing larval survivorship [18].

The behavior of *L. cuprina* and other blowflies in locating hosts for oviposition has been extensively studied [13]. The main attractants to gravid females are sulfur-rich compounds, ammonia and other volatiles emitted from decomposing organic material, such as meat or feces [1,13,21]. Dimethyl disulfide (DMDS), dimethyl trisulfide (DMTS), Indole and p-cresol are among the most relevant blowfly attractants collected from decomposing organic material [22–24]. Adult gravid females and the last larval stage of *L. cuprina* are more attracted to 5-day-old rotten beef compared to fresh beef [25,26]. In addition, larvae preferred cold beef (∼25°C) rather than warm (∼33°C) [26]. These observations could point to sensory systems conserved between the adult and larval stages of *L. cuprina*.

RNA-Seq studies have been addressed different larval biological processes in blowfly species, such as development [27], metabolism [28], aging [29], adaptation to temperature changes [30,31] and immune response [32]. However, none of these studies focused on larval sensory receptors potentially associated with physiological and behavioral processes. In *Drosophila melanogaster* Meigen, 1830, different sensory receptors, including odorant (ORs) and gustatory receptors (GRs), have been linked to many larval biological processes such as chemotaxis, thermotaxis and locomotion [33,34]. For example, *DmelOR42a* and *DmelOR42b* appear to be important for larval olfaction and locomotion [35–39]. Regarding GRs, such as *DmelGr64a*, have been associated with taste [40], whereas others such as *DmelGR28* have been associated with light avoidance and thermal sensing [41,42]. Ionotropic receptors (IRs) were also associated with taste and thermal sensing in the *Drosophila* larvae [43,44].

Among other sensory receptors, the transient receptor potential channels (TRPs) and *Pickpocket* receptors (PPKs) were identified in the *Drosophila* larval chordotonal organs and showed a role in proprioception, thermotaxis and locomotion [45–48]. DmelTRPN1 (*nompC* gene) and other TRPs belonging to the ankyrin and vanilloid subfamilies were associated with proprioception [48] and thermotaxis [46,47,49]. Among odor-molecule carriers, odorant-binding proteins (OBPs) and chemosensory proteins (CSPs) were identified in the larva of *D. melanogaster* [50]. However, the role of these proteins in the larval olfaction and other physiological aspects has not been extensively studied. Further, the role of other chemosensory proteins previously studied in adult flies [51–53], such as the superfamily ML (MD-2 (myeloid differentiation factor-2)-related Lipid-recognition and Niemann-Pick C2 disease proteins (ML/NPC2), CD36-sensory neuron membrane proteins (CD36/SNMPs), and ammonium transporters (AMTs), remains to be addressed.

The main objective of the present work was to identify sensory receptors and other genes relevant to the physiology of *L. cuprina* larvae. We aimed to highlight genes potentially associated with the processes mentioned above, and other physiological functions such as feeding, digestion, excretion, diuresis and immune response. Our RNA-Seq analysis of different larval stages and tissues suggested that the ORs, GRs, IRs and PPKs may not play a central role in the larval sensory signaling and digestive systems of *L. cuprina*, except for *LcupGR94a* which was biased to the larval gut. ABC transporters and OBPs were the most relevant within the sensory-related gene families in terms of number of identified sequences and gene expression, both in the larval head and across all larval stages. Since the odorant coreceptor (*Orco*) is a necessary element for the normal functioning of all ORs [54], we tested both adult females and larvae homozygous for a loss-of-function mutation created by gene editing. Behavioral assays showed that *LcupOrco* was essential for the female response to rotten beef, but not for larval behavior. These findings are consistent with high levels of expression of *LcupOrco* in the adult female antenna [25] but very low expression in larvae (present work).

## Methods

### Insect rearing conditions

All larvae and flies used in the present study were of the sub-species *Lucilia cuprina cuprina*, called “*L. cuprina*” to ease the reading of the manuscript. The *L. cuprina* LA07 wild-type (*wt*) colony is currently at approximately 225 generations and was established in 2010 using 300 individuals kindly provided by Dr. Aaron Tarone (Texas A&M, TX, USA). The LA07 colony was established by Dr. Tarone from multiple collections of individuals (300 to 500) from the University of Southern California campus and the Miracle Mile neighborhood, in Los Angeles, CA, USA in 2007. All larvae and adult flies used for the experiments were reared using protocols previously described [9]. Adult flies and immature stages were kept at 23.5±1°C a non-controlled photoperiod (∼13:11 light/dark).

To complete the RNA-Seq analysis, *wt* and heterozygous individuals were used, the latter obtained from crossing *wt* and SLAM5X colonies. SLAM5X carries an X-linked constitutively expressed DsRed fluorescent protein gene plus the tetracycline transactivator coding sequence driven by the early embryo *Chslam* promoter [55]. For the behavioral assays, *LcupOrco* larvae or adult females were obtained from crossing heterozygous *LcupOrco* (*LcupOrco^+/-^*) siblings. The resulting *LcupOrco* mixture was ∼ 25% *wt*, 50% *LcupOrco^+/-^* and 25% *LcupOrco^-/-^*. In addition, adult males from the *NPF^-/-^*colony expressing the ZsGreen were used for the mating assay.

### RNA-Seq experiments

#### Experimental design and sample collection

Two RNA-Seq experiments, designated EXP-1 and EXP-2 were performed. EXP-1 addressed gene expression pattern across the three larval development stages using RNA isolated from whole first, second and late wandering third instar (L1, L2, L3). EXP-2 used RNA isolated from head, gut and whole body early feeding third instar. For EXP-1 there were five replicates for L1 and L2 and six replicates for L3. Samples one to three of L1 and L2, and all samples for L3, were *wt* larvae. Samples four and five of L1 and L2 were collected from the *wt*-SLAM5X cross. Each replicate was a pool including 10 to 15 larvae for L1 and L2, and three pooled individuals for L3. L1 samples were collected between 20 to 30 hours (h) after egg laying, 50 to 55 h for L2 and between 130 to 135 h for L3 (Fig. S1A-D). All samples were flash frozen using liquid nitrogen after collection and stored at -80 °C until use.

Samples included in EXP-2 were the first segment of the larva designated as head (H), the gut including the midgut plus the hindgut and without the crop and Malpighian tubules (MT) designated as gut (G), and the whole larva (WL) including all tissues (Fig. S1E-G). The crop was removed from the gut because the presence of meat from the larval diet withing this organ could bias the analysis. All tissues were collected from *wt* L3s of ∼ 75 h after egg laying. Four replicates were collected for H and G and three for WL, and each replicate was a pool including 50 first larval segments for H, 10 guts for G and five whole larvae per pool for WL. Tissues were dissected using dissection forceps and vannas scissors (WPI, Worcester, MA, USA), and immediately after placed in a 1.5 mL Eppendorf tube filled with 300 µl of cold RNAlater^TM^ (Thermo-Fisher, Waltham, MA, USA, Cat. # AM7020) and stored at -80°C until use.

#### RNA isolation

RNA extraction of samples associated with EXP-1 and EXP-2 was completed using two different protocols as follows. Whole larvae of EXP-1 were transferred using dissection forceps to 2.0 mL tubes prefilled with zirconium beads (Benchmark Scientific, Tempe, AZ, USA, Cat. # D1032-30). Subsequently, the larvae were resuspended in 250 µl of cold Trizol^TM^ (Thermo-Fisher, Cat. # 15596026) and disrupted using a benchtop homogenizer (Benchmark Scientific, Cat. # Z742475). The homogenizer was set to 6.5 m/s and samples were disrupted for 1 min, another 250 µl of cold Trizol^TM^ were added to the tubes and the homogenization process was repeated once more. After homogenization, total RNA was extracted using the RNeasy Kit (Qiagen, Hilden, Germany, Cat. # 74104), and genomic DNA (gDNA) was removed using the DNase kit (Qiagen, Cat. # 79254) following manufacturer’s specifications.

Before RNA extraction of samples belonging to EXP-2, the RNAlater^TM^ was removed from tubes and tissues were rinsed two times with 1 mL of cold 50% ethanol. After rinsing, pools of heads, guts and whole larvae were resuspended in 250, 500 and 1000 µl of cold Trizol^TM^ respectively. Samples were kept on ice and tissues were disrupted using pellet pestles (Bel-Art, Wayne, NJ, USA, Cat. # F65000-0002) with a homogenizer (Bel-Art, Cat. # F65100-0000) for 90 seconds (sec). Subsequently, samples were centrifuged at 16000 x g and 4°C for 5 min to pellet part of lipids, cuticle and other debris. After centrifugation the supernatant was transferred to a clean 1.5 mL tube. Total RNA was extracted using the Quick-RNA^TM^ kit (Zymo Research, Irvine, CA, USA, Cat. # R1050) according to the manufacturer’s specifications. The exception was the addition of a second deoxyribonuclease 1 (DNase 1) step to ensure the absence of gDNA contamination in the samples, as follows: 1) during the RNA extraction 30 U of DNAse1 plus 75 µl of digestion buffer (DB) were added to the crude samples followed by 15 min of incubation at RT; 2) after elution of the clean RNA in nuclease-free (NF) water, another 5 U of DNAse1 + 2.5 µl of DB were added followed by an incubation time same as above. After DNAse1 treatments, the RNA Clean & Concentrator^TM^-5 kit (Zymo Research, Cat. # R1013), was used to clean the samples following the manufacturer’s specifications.

Clean RNA belonging to samples from EXP-1 and EXP-2 was eluted in 30 and 12 μl of NF-water, respectively, and was quantified (1:10 dilution) using a Qubit-4^TM^ fluorometer and the HS kit (Thermo-Fisher, Cat. # Q33120). Samples were kept at -80°C until use.

#### RNA-Seq data analysis

Sample quality control, library construction and sequencing services were provided by Novogene Inc. (Sacramento, CA, USA), following protocols described by the provider (https://www.novogene.com/us-en/resources/downloads/). The sequencing platform was Illumina NovaSeq 6000 (Illumina, San Diego, CA, USA), using paired end 150 bp reads, and the sequencing depth coverage was ∼ 50 and 400 million raw reads per library for EXP-1 and EXP-2, respectively.

The Geneious Prime® software v2023.2.1 (https://www.geneious.com) and associated plugin packages, were used to complete all the bioinformatic analysis and generate the RNA-Seq data output. BBDuk plugin v38.88 was utilized to remove Illumina adaptors and trim off low-quality bases at the 5′ and 3′ends using the following parameters: kmer length: 27; trim both ends: minimum quality Q20; trim adapters based on paired reads overhangs: minimum overlap 22; discard short reads: minimum length 20 bp. Trimmed reads were mapped to the reference *L. cuprina* genome assembly NCBI ID ASM2204524v1, using the default Geneious Prime mapper set to detect all types of RNA sequences and low-medium sensitivity. A second *L. cuprina* genome assembly, NCBI ID GCA_001187945.1, was used to analyze specific genes with annotation errors in the first assembly. Gene expression was calculated only for identified coding sequence (CDS) and the normalization method used was transcript per million (TPM), but RPKMs (Reads Per Kilobase per Million mapped reads) FPKMs (Fragments Per Kilobase per Million mapped fragments) were also provided with the raw data.

A principal components analysis (PCA) was performed to compare EXP-1 libraries with the R packages FactoMiner [56] and ggcorrplot [57] used for analysis and plotting, respectively. Gene expression in TPMs was plotted using GraphPad Prism v9 software (San Diego, CA, USA), showing the mean of each analyzed gene and the standard deviation (SD) between libraries for the same gene. Differential gene expression between groups was calculated using DESeq2 [58]. The EXP-1 included five libraries for groups L1 and 2, and six for L3, and EXP-2, four libraries for groups H and G, and three for WL. A Benjamini-Hochberg (BH) adjusted value < 0.05 and a FC > 0 were used to define differentially expressed (DE) transcripts between groups. Sequences with a TPM expression ≥ 5 were separated from all DE transcripts and were used for all analyses. Transcripts with low expression may not be translated into proteins in the tissues analyzed or may be associated with pseudo-genes or isoforms [59]. Consequently, DE ABC and OBP gene families surpassing 5 TPMs were plotted. Results for other sequences outside this selection were provided as supplementary data tables.

#### Phylogenetic analysis

The phylogenetic analysis was completed for the ABC and OBP sensory families. ABC protein sequences were identified by running a NCBI BLASTp against the *L. cuprina* genome assembly, NCBI ID ASM2204524v1, and using *D. melanogaster* ABC protein sequences as query, the latter obtained from the FlyBase database (https://flybase.org/). *L. cuprina* OBP curated protein sequences were obtained from a previous study performed in the same species [25]. Only *L. cuprina* sensory receptors showing a TPM expression ≥ 5 in the present work were included in the final phylogenetic analysis. If there was more than one isoform per gene, only isoforms with different transcript expression were added. Further, OBPs and ABC transporters previously identified as larval biased in *L. sericata* [30] were added to the analysis.

Prior to phylogenetic analysis, a sequence alignment was performed using the G-INS-i method in MAFFT v7 [60]. A preliminary phylogenetic analysis including all ABC receptors identified for *L. cuprina* with a TPM expression > 0, using the Neighbor-Joining (NJ) algorithm [61] and the software MEGA v11 [62] was performed to rename *L. cuprina* ABCs based on direct orthology to *D. melanogaster* sequences. Subsequently, the Maximum likelihood (ML) method set for 1000 bootstrap replications (using the same software) was utilized to improve a second NJ tree completed including only ABC receptors with a TPM expression ≥ 5. The best substitution model for each ML run was determined by using the tool “Find Best DNA/Protein Models” of the same software, prior to running the phylogenetic test. Tree figures were produced using the interactive Tree of Life (iTOL) v6.8.2 software (https://itol.embl.de/). The analysis for the OBP sequences was completed using the same protocol except for the renaming process.

### Odorant coreceptor (*Orco*) experiment

#### Gene editing using CRISPR/Cas9

##### LcupOrco gene sequence, guide RNA and Cas9 in vitro assay

The guide RNA (gRNA) selected for gene editing was the ortholog of a gRNA previously used to edit the *ChomOrco* gene in *C. hominivorax* [63]. Note S1 provides further details about the *LcupOrco* gene sequence including exonic regions, gRNA position, primers used to amplify the *LcupOrco* gene fragment encompassing the Cas9 cutting site, and the Polymerase Chain Reaction (PCR) cycling parameters. The *LcupOrco* gene model showing the gene editing site was edited with Adobe Illustrator (San Jose, CA, USA), and the LcupOrco protein secondary structure generated with the same purpose was plotted with the Protter software (https://wlab.ethz.ch/protter/start/). The gRNA used was CRISPR-Cas9_crRNA [Integrated DNA Technologies (IDT), Coralville, IA, USA]. The latter was mixed with the CRISPR-Cas9_tracrRNA (IDT, Cat. # 1072532), using an equal volume of both RNAs after diluting them to the same concentration. Subsequently, the mix was heated at 95°C for 5 min, cooled to ∼ 15°C for 5 min and kept at -20°C until use.

The gRNA cutting efficiency was tested *in vitro* using Cas9 protein [New England Biolabs (NEB), Ipswich, MA, USA, Cat. # M0646T]. Cas9 and gRNA were mixed for a final concentration of 66.6 and 160 nM, respectively, and incubated for 20 min at 25°C to form the Cas9 ribonucleoprotein (RNP) complex. A PCR-generated fragment containing the targeted sequence (Note S1), was added to the mix for a final concentration of 3 nM and incubated for 60 min at 25°C. The reaction (RXN) was ended by adding 1 μl of Proteinase K 10 μg/μl (Zymo, Cat. # D3001-2-20) and incubated for 10 min. The RXN mix was then loaded into a 1.5% agarose gel in parallel to uncut gDNA used before as a template (same concentration), electrophoresed at 60 volts for 90 min and visualize using a Gel Doc™ EZ System (Bio-Rad, Hercules, CA, USA).

##### Insert construction, CRISPR/Cas9 mix and embryo injections

A Synthetic DNA fragment including two sections of the *LcupOrco* gene, plus restriction cutting sites for the enzymes XhoI (NEB, Cat. # R0146S) and NotI (NEB, Cat. # R0189S) was acquired cloned within the plasmid pUCIDT-AMP GoldenGate (IDT). The ZsGreen fluorescent marker later added to the same construct was obtained and purified from a donor plasmid [64]. The pUCIDT-AMP GoldenGate plasmid and ZsGreen fragment were ligated using T4 Ligase (NEB, Cat. # M0202S) and then used for transformation of commercial competent *Escherichia coli* cells (NEB, Cat. # C3019H) following the manufacturer’s specifications. Plasmids were obtained from 10 single clones using the ZR Miniprep Kit (Zymo, Cat. # D4016) following the manufacturer’s specifications. Subsequently, plasmids were evaluated by restriction enzyme digestion analyses. Plasmid DNA sequencing was performed for plasmids that showed the correct banding patterns (MGH CCIB DNA core, Boston, MA). For embryo microinjection, plasmid DNA was obtained by using a Midiprep Kit (Zymo, Cat. # D4200) and further purified using the DNA Clean Kit (Zymo, Cat. # D4004) following the manufacturer’s specifications. Prior to injection the DNA was diluted in NF-water to 1250 ng/μl concentration. For further details refer to Fig. S2A-F.

The Cas9 (IDT, Cat. # 1081058) was used for the *in vivo* embryo injections; the same was mixed with the duplex crRNA::tracrRNA, 1M Potassium chloride (KCl) and 1 μl of 3.1 buffer (NEB, Cat. # B6003S) and incubated at 25°C for 20 min to form the RNP complex. The mixture was cooled to room temperature for 5 min and the plasmid carrying the *LcupOrco* insert was added to the mix to reach a final plasmid concentration of 500 ng/μl. Final concentrations for the other compounds were as follows: Cas9 = 3.1 μM (500 ng/μl); crRNA::tracrRNA = 7.5 μM (250 ng/μl) and KCl = 250 mM. The mix was cleaned before injections by 2 min centrifugation at 4°C using a 0.45 μM filter column (Sigma-Aldrich, Burlington, MS, USA, Cat. # UFC30HV) and keep on ice all the time.

The needles used for microinjections were generated by pulling filamented quartz capillaries (Sutter Instrument, Novato, CA, USA, Cat. # QF100-70-10). The puller used was a Sutter Instrument P-2000 set for: Heat = 690; Filament = 4; Velocity = 60; Delay = 160; Pull = 170. After pulling, needle’s tips were beveled for ∼ 10 seconds (sec) using a BV-10 micropipette beveler (Sutter Instrument), a fine abrasive plate and an angle of 22.5°.

Before injections, microinjection mix was loaded into the needles using a Microloader™ (Eppendorf, Hamburg, Germany, Cat. # 930001007). The microinjector used was a XenoWorks Digital Microinjector (Sutter Instrument), set for continuous flow, 1000 hectopascal (hPa) of pressure and +5 hPa of transfer pressure, together with a motorized XenoWorks Micromanipulator (Sutter Instrument) and a stereomicroscope M125 (Leica, Wetzlar, Germany). Flies used for egg collection were LA07 8-day-old gravid females kept with males of the same age in a plastic container (Fig. S3A-B). The flies were stimulated to begin ovipositing by adding 93/7% lean-fat fresh grounded beef placed in a 35 mm Petri dish for 15 min. Following, another 35 mm-plate with fresh meat was added to collect new eggs for 5 min. An egg cluster was transferred to a piece of tissue paper (Kimberly-Clark, Roswell, GA, USA), moistened with distilled (DI)-water and located in one of the depressions of a double-depression microscope slide (VWR, Radnor, PA, USA, Cat. # 470003-480). Twenty eggs per round of injections were separated under an Olympus stereomicroscope SZX10 (Olympus, Center Valley, PA, USA), using a 00-paint brush and accommodated onto a Scotch^®^ double-sided film tape (3M, Saint Paul, MN, USA), located in the other depression of the same slide. The eggs were then desiccated for 5 min using a desiccation chamber of 150 mm in outer diameter (Nalgene, Rochester, NY, USA) containing a desiccant cartridge (SP Bel-art, Wayne, NJ, USA, Cat. # F42049-0100). Subsequently eggs were covered with oil (Sigma-Aldrich, Cat. # H8773), rest for 3 min and injected at the posterior end for ∼ 1 sec. The developmental time for the injected embryos was between 18 to 24 min, for the first and last injected embryo, respectively. After injection, embryos were kept overnight in an incubator chamber (Billups-Rothenberg, Del Mar, CA, USA, Cat. # MIC-101) filled with oxygen and protected from light using a carboard cover until eggs hatched, *i.e*. between 19 to 27 h at 23.5°C. After hatching, newly emerged L1s were transferred using a 00-paint brush to moistened tissue paper (Kimberly-Clark) and then moved to a 3-ounces clear plastic cup filled with 50 grams of the same meat mentioned above and let larvae develop until tested at L3 stage.

##### Fly screening, crosses and genotyping

Injected embryos were screened 7 days after injections (L3 stage) using a stereo microscope M205 FA (Leica) and a green filter. Only mosaic embryos from generation (G)0 showing transient expression of the plasmid were retained to obtain adult flies. Emerged adult flies were crossed following the protocol described in Fig. S4. G1 and G2 eggs obtained from G0 and G1 crosses, respectively, were tested for positive fluorescence using the same microscope, and only fluorescent eggs were kept.

At G3, a middle leg was collected using dissection forceps and scissors from 2-day-old flies for genotyping. gDNA was extracted by placing legs in a DNA extraction buffer (Biosearch Technologies, Hoddesdon, UK, Cat. # QE09050) for 15 min at 65°C and 2 min at 98°C. Three strategies were used to determine if the *LcupOrco* insert was landed at the correct *L. cuprina* genome position, *i.e*. withing the *LcupOrco* coding region: 1) amplification of left side of the insert + upstream genome; 2) amplification of right side of the insert + downstream genome; and 3) amplification of the whole insert including upstream and downstream genome regions; for further details refer to Note S2. In parallel, potential indels and point mutations at the Cas9 cutting site were investigated using the same primers described in Note S1. After PCR amplification, the gDNA was purified using the DNA Clean Kit and sequenced by Sanger Sequencing. Amplicons >1 Kb were cloned using the pGEM®-T system (Promega, Fitchburg, WI, USA, Cat. # A1360) together with competent cells following the manufacturer’s specifications. Plasmids were obtained, purified and sequenced following the same protocols described in Fig. S2.

#### Behavioral assays

##### Larval diet preference test

The larval diet preference test was previously used to evaluate *L. cuprina* larvae olfaction behavior [26]. The larvae used for the assay were obtained using protocols described at the *Insect rearing conditions* section. Before the assay, ∼ 250 L1s were transferred to 50 g of fresh ground beef 1 day after egg hatching. Two days after this step a total of 120 3-day-old larvae were tested following protocols described at [26]. The options offered to the larvae were rotten (Ro) and fresh (Fr) ground beef, either at 25±1 or 33±1°C, designated as cold (Co) and hot (Ho) respectively, and for further details about beef conditions refer to [26]. Those larvae that did not choose any option after 10 min were considered as non-choice (NC). After the assay, larvae were placed in separated 1.5 tubes filled with 300 µl of DNA preservative buffer (Zymo, Cat. # R1100-50) and kept at -80°C until DNA extraction. Subsequently, all larvae were dried using tissue paper and separately weighed using a LA204E analytical balance (Metter Toledo, Columbus, OH, USA). Following, a small section of each larva was collected using dissection forceps and scissors and placed back in the same buffer. Approximately one third of the larvae associated with each diet preference, giving a total of 48 larvae, were genotyped using same protocols described at the *Fly screening, crosses and genotyping* section and Note S1.

##### Adult female olfaction assay

The adult female olfaction assay was completed using the same protocol, spatial olfactometer and room conditions previously used to evaluate *L. cuprina* female olfaction behavior [25]. The olfactometer consists of four separate chambers (replicates) where the flies were released. Each of these chambers are connected with two smaller collection chambers A and B, where meat samples were placed to attract and trap the flies. The attractants used to lure the flies were 2 g of fresh (Fr) and rotten (Ro) beef at room temperature, placed in chambers A and B, respectively.

The females used for the olfaction assay were from the same cohort of individuals used for the larval diet preference test (see above). In addition, LA07 *wt* females were tested using the same assay conditions, but on different days without mixing them with the *LcupOrco* flies. All flies used for the olfaction assay were 10-day-old gravid females and were separated from males the day before the assay using CO_2_ to anesthetize the flies under a stereomicroscope SMZ745 (Nikon, Melville, NY, USA). After sexes separation, females were kept in the same room where the assay was performed for habituation, provided with tap water and cane sugar. After the assay, females were kept in separated 1.5 mL empty tubes at -80°C until DNA extraction. A total of 80 *wt* and 174 *LcupOrco* females were tested, and 72 of the latter, randomly selected from all assayed *LcupOrco* females, were chosen for genotyping using the same protocols described in the previous section. After the assay, females were separated into three groups: 1) attracted to fresh (Fr) beef; 2) attracted to rotten (Ro) beef; and 3) flies that did not respond to any stimulus, designated as non-choice (NC).

##### Mating assay

To evaluate the mating performance of the *LcupOrco* homozygous females (*LcupOrco*^-/-^), 5-day-old females previously determined as heterozygous or homozygous females by genotyping protocols described above, were mixed with twice as many adult *NPF*^-/-^ males of approximate same age, constitutively expressing the ZsGreen fluorescent protein marker. Fifty grams of fresh grounded beef was offered O/N to females at 3, 10 and 17 days after mixing them with the *NPF*^-/-^ males. After oviposition, eggs were visualized and photographed using a stereo microscope M205 FA (Leica), a green filter and a Leica K5C camera.

## Results and Discussion

### RNA-Seq overview

To identify genes that are differentially expressed at the very anterior end of the larvae, the first segment (excluding the brain) was dissected from early feeding third instar (Fig. S1F). RNA was isolated from gut dissected from the same stage to identify genes important for meat digestion. Lastly RNA was obtained from whole first, second and third instar (early and late) for reference for the two tissues, and for a larval developmental series. RNA-Seq output including reads per library, total number of sequences mapped and subtotals per type of RNA is provided in Table S1A-B. Considering only transcripts surpassing 5 TPMs, the number of transcripts ranged between 5,617 and 6,642 for different larval stages and tissues (Data S1A-C and S2A-C). Further, a PCA analysis showed that samples belonging to each larval stage clustered together, highlighting the differences between larval stages over those between the samples within each group (Fig. S5). An RNA-Seq analysis of different larval tissues of *L. sericata* showed a similar number of transcripts expressed over the same expression threshold, between ∼ 4,400 and 6,000 [30]. Transcripts below this expression threshold for each library are provided in Data S1D-S and S2D-N.

### *Lucilia cuprina* larval sensory receptors

ABC transporters and OBPs compromised about 80% of the sensory receptors/carriers showing a TPM expression ≥ 5, and the number of receptors/carriers per gene family was as follows: 49 ABCs; 25 OBPs; eleven CD36/SNMPs; six ML/NPC2s; three CSPs and one for each of the following gene families, AMTs, GRs and TRPs (Data S3 and S4). Furthermore, OR, GR, IR and PPK receptors were not observed above the expression threshold of 5 TPM, except for *LcupGR94a* in the larval gut (Data S4A). The expression level of ORs, GRs and IRs suggests that the same may not have a central role in the sensory system of *L. cuprina* larvae. For instance, *LcupOrco,* a highly expressed sensory coreceptor in the *L. cuprina* antenna [25] showed a TPM expression < 0.5 in whole larvae and in head and gut (Table 1). However, in *D. melanogaster* larvae, *Orco* expression is restricted to the olfactory dorsal organs at the anterior tip [65] and is essential for chemotaxis [66]. Furthermore, the role of many ORs, GRs and IRs has been confirmed by *in situ* hybridization and functional assays [33–36,41–44]. Consequently, more experiments are required to elucidate the role of the specified receptors within the sensory system of the *L. cuprina* larvae.

**Table 1.**
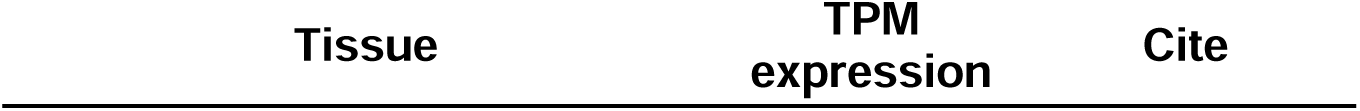
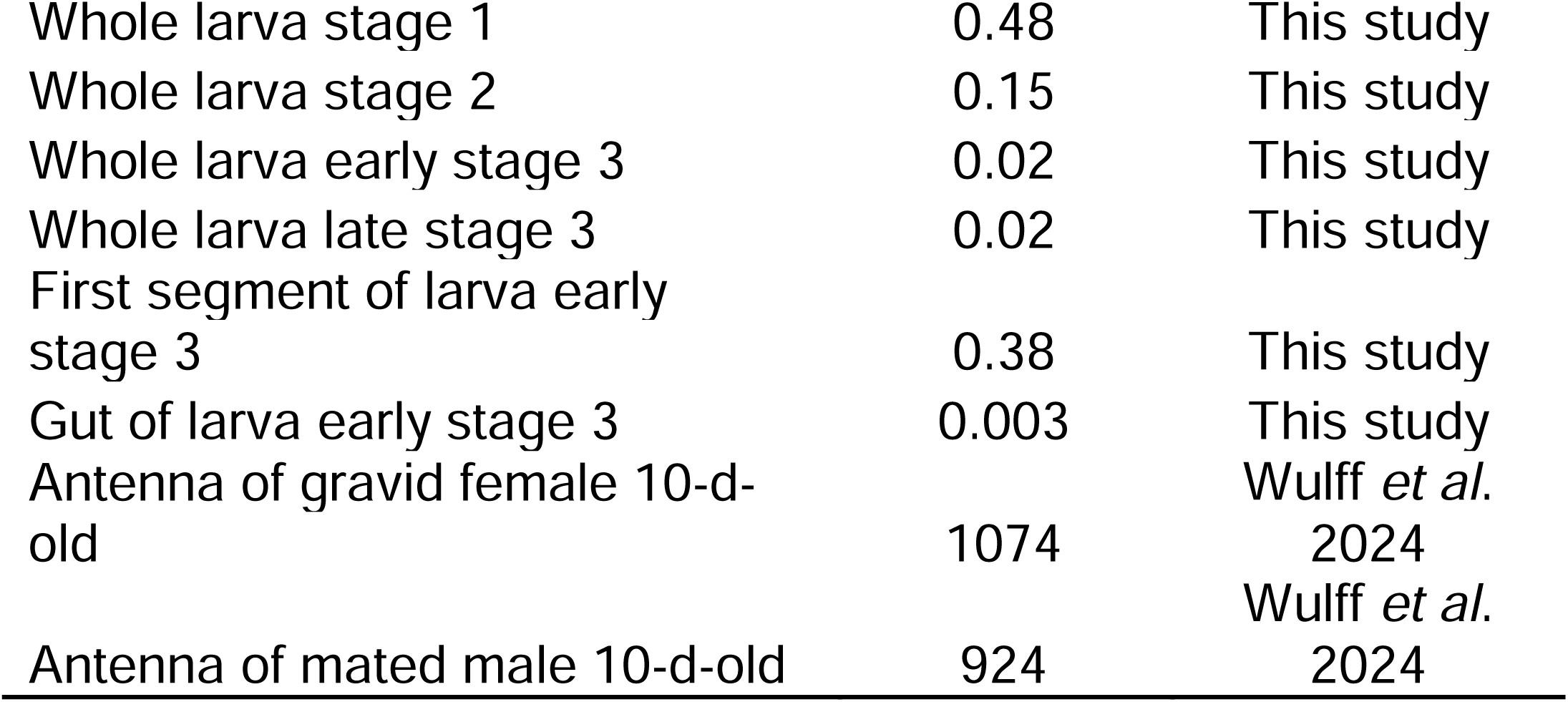
*Lucilia cuprina Orco* expression in adult and larval tissues

| Tissue | TPM<br>expression | Cite |
| --- | --- | --- |
| Whole larva stage 1 | 0.48 | This study |
| Whole larva stage 2 | 0.15 | This study |
| Whole larva early stage 3 | 0.02 | This study |
| Whole larva late stage 3 | 0.02 | This study |
| First segment of larva early stage 3 | 0.38 | This study |
| Gut of larva early stage 3 | 0.003 | This study |
| Antenna of gravid female 10-d-old | 1074 | Wulff *et al.* 2024 |
| Antenna of mated male 10-d-old | 924 | Wulff *et al.* 2024 |

The ABC transporters are classified into eight subfamilies [67]. A phylogenetic analysis showed a possible clade expansion for the subfamily G in *L. cuprina* (Fig. 1 and Fig. S6). Similar expansions for the same subfamily were observed in hematophagous dipterans [68]. In *L. sericata*, orthologs of LcupABCGs 4, 6 and 7 were highly expressed in larval Malpighian tubules (MT) and fat body (FB) [30]. Here we found that members of this subfamily were highly enriched in the larval anterior end (see below). In contrast to the apparent expansion of subfamily G, subfamily A showed a possible clade contraction compared to *D. melanogaster* (Fig. 1), with only four receptors identified for *L. cuprina* (Data S5B). In line with these findings, clade contractions and expansions are common for subfamilies A and G along insect species [69]. Sequences used in the analysis are provided in Data S5A-D; annotation of two sequences was corrected and transcript expression and differential expression calculated using genome assembly GCA_001187945.1 (Data S6).

**Figure 1.**
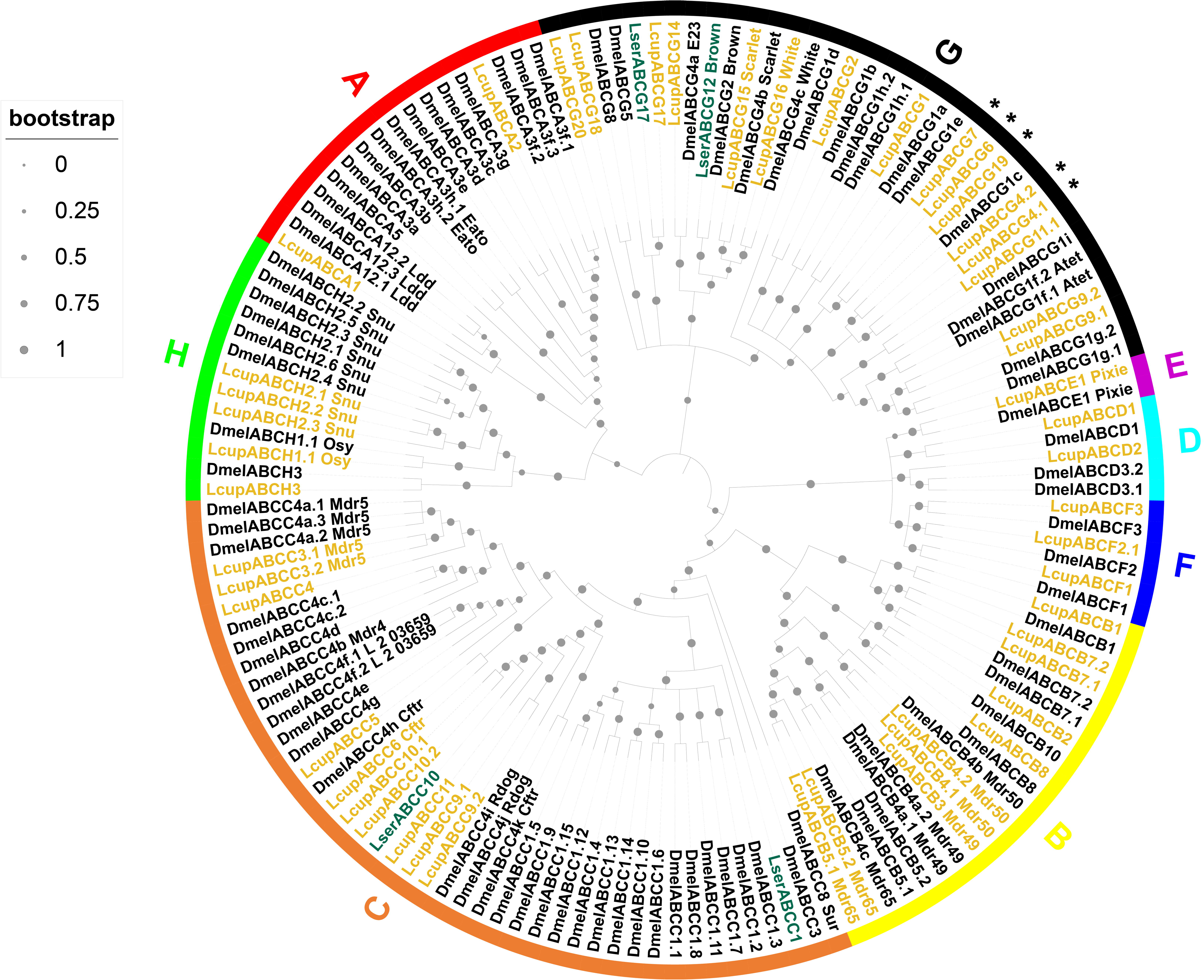
Phylogenetic analysis of the ABC transporters. Three Diptera species were used for the phylogenetic analysis as follows: *L. cuprina* (Lcup, gold), *L. sericata* (Lser, green) and *D. melanogaster* (Dmel, black). ABC subfamilies from A to H and node bootstrap supporting values were detailed. Only *L. cuprina* receptors with a TPM expression ≥ 5 in larval head, gut or whole body were used for the analysis, but all ABC transporters of *D. melanogaster* were included. In addition, ABC transporters previously identified as larval biased in *L. sericata* [30] were added to the analysis. If there was more than one isoform per gene, only isoforms producing a different amino acid sequence and showing a dissimilar transcript expression were added. Potential expanded *L. cuprina* ABC transporters belonging to subfamily G were detailed with an asterisk. For more details about sequences used in the analysis refer to Data S5A-D.

*Lcuppixie* was the most highly expressed ABC transporter (Data S3A and S4B) and upregulated in the first two instar compared with third (Fig. 2A). This gene belongs to subfamily E and is essential for protein translation [70]. Receptors belonging to subfamily F, associated with protein translation and immune response [71], and B, associated with xenobiotic detoxification and insecticide resistance [67], were also upregulated in the first two instar larva vs. L3 (Fig. 2A, Data S3A). No member of these subfamilies was head-biased, but five ABCs of subfamily B were gut-biased (Fig. 3A-C, Data S4B). Subfamily G showed higher expression in the latter instars (L2 or L2/L3) compared with the first instar (Fig. 2B-D). In addition, this subfamily had the highest number of head-biased genes (Fig. 3A and Data S4B), with some showing a very high enrichment compared with gut or whole body (Data S7A). This subfamily includes orthologs of the well-studied Drosophila pigmentation genes *white* and *scarlet* [67], which are L2-biased in *L. cuprina* (Fig. 2B) but do not show any enrichment in head or gut (Fig. 3A-B).

**Figure 2.**
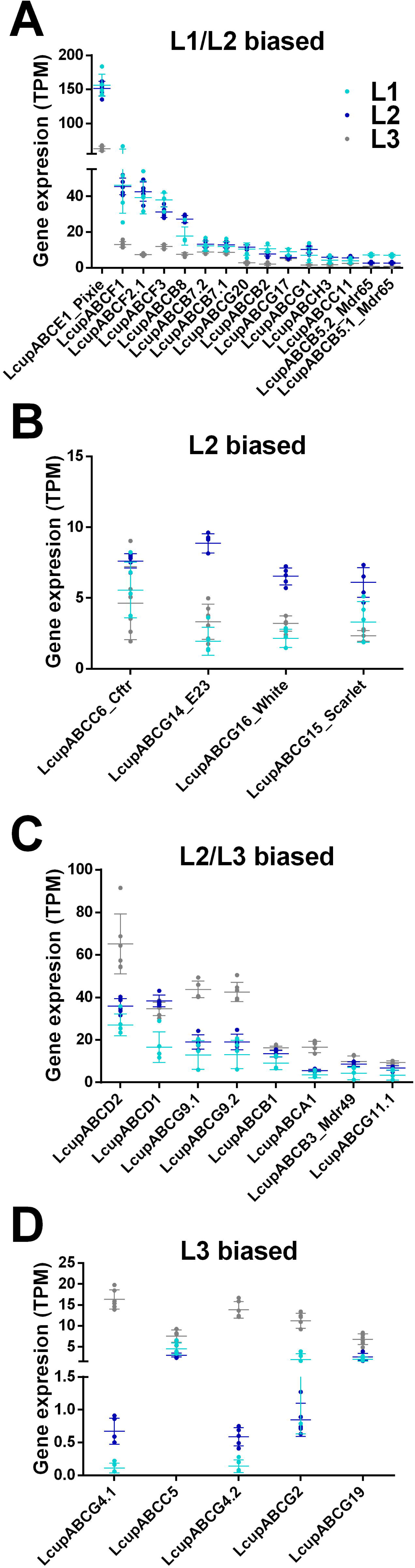
Expression and differential expression of the ABC transporters in *L. cuprina* larval stages. The TPM expression of ABC receptors was plotted clustering the same according to differential expression between the three larval stages as follows: (**A**) includes receptors biased to the first two larval stages (L1 and L2) vs. L3; (**B**) shows receptors only biased to the L2 stage; (**C**) groups receptors biased to the last two stages (L2 and L3) vs. L1; and (**D**) the receptors only biased to the last stage (L3). Only receptors with a TPM expression ≥ 5 were added. Dots represent the gene TPM expression of each library; the average mean expression per gene, plus the SD between libraries were also plotted. For more details about the expression level and differential expression of each receptor refer to Data S3A.

**Figure 3.**
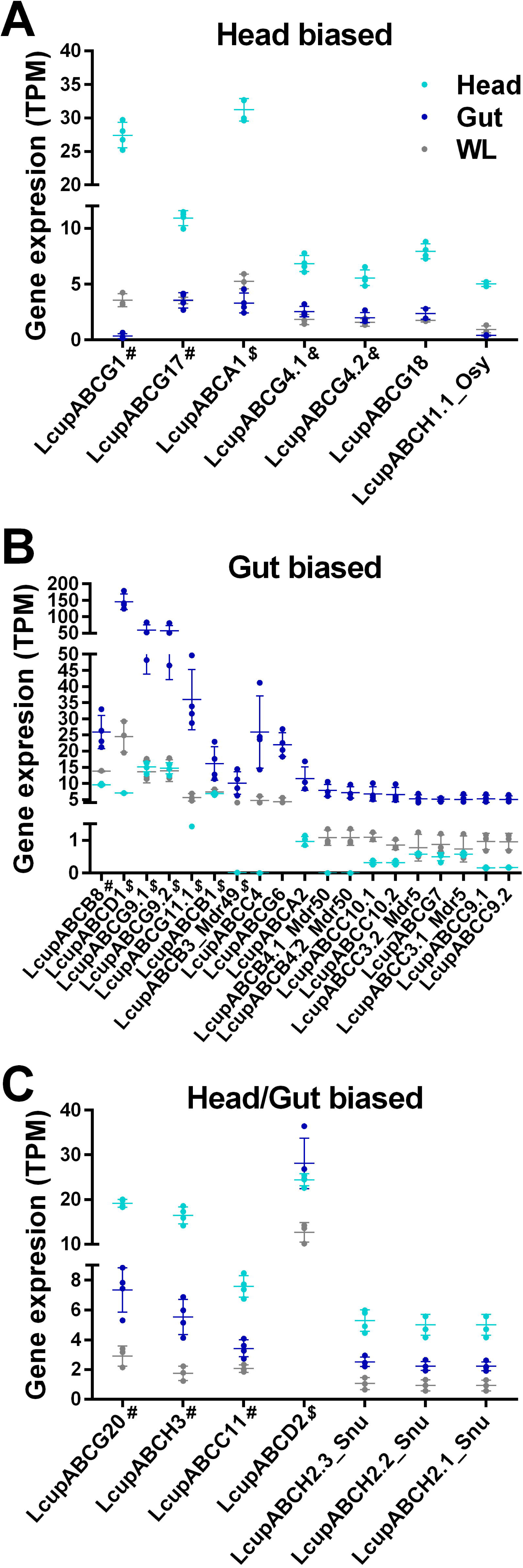
Expression and differential expression of the ABC transporters in *L. cuprina* third instar larval tissues. The TPM expression of ABC receptors was plotted clustering the same according to differential expression between different tissues as follows: (**A**) includes receptors biased to the larval head (first segment of the larval body) vs. whole body; (**B**) shows receptors biased to the gut vs. whole body; and (**C**) groups receptors biased to head plus the gut vs. whole body. Only receptors with a TPM expression ≥ 5 were added. Dots represent the gene TPM expression of each library; the average mean expression per gene, plus the SD between libraries were also plotted. Symbols next to receptor’s names represent larval stage biased expression (plotted in Fig. 2) as follows: # L1/L2-biased; $ L2/L3-biased; & L3-biased. For more details about the expression and differential expression of each receptor refer to Data S4B.

Subfamily H was also well represented in the larval head, with the *L. cuprina* ortholog of the Drosophila *oskyddad* (*osy*) showing head-biased expression (Fig. 3A) and the ortholog of *snustorr* (*snu*) enriched in head and gut (Fig. 3C). In *D. melanogaster*, *osy* and *snu* are associated with lipid deposition on the cuticle [72,73]. In the larval gut, the best represented subfamily was C followed by B (Fig. 3B). Both subfamilies share a role in xenobiotics detoxification, multidrug resistance and protein translation [67]. ABC-Cs are also involved in the binding of sulfur-rich compounds [74], and ABC-Bs in ion and lipid transportation [67] and iron/heme metabolism in hematophagous insects [75]. Sulfur-rich compounds are common blowfly attractants [13] and iron and heme are both present in the *L. cuprina* larval meat diet [76]. Other genes engaged in processing these compounds, such as *heme oxygenase 1* and *ferritin* were also highly expressed and biased to the larval gut (Data S2B and S8A-B).

The second highest number of chemosensory proteins identified in all *L. cuprina* larval stages and tissue transcriptomes were grouped by OBPs (Data S3B and S4C). This gene family is classified based on a cysteine (C) conserved motif, and modifications of the same change the OBP-ligand binding affinity [54]. The classic subfamily includes a six C-motif and in the Dimer subfamily, the whole motif has been duplicated [54]. Other subfamilies are Minus-C, where the C2 and 5 from the classic motif have been lost, and Plus-C, that includes the classic motif plus three extra conserved Cs and a conserved proline immediately after the Cs [54]. The OBP phylogenetic analysis suggested a clade expansion for *L. cuprina* and *L. sericata* classic OBPs 39 to 44 (Fig. 4). The *L. sericata* OBPs included in the expanded clade were highly expressed in the FB followed by the MT [30], suggesting they may be associated with lipid binding and excretion. OBPs grouped within the clade LcupOBPs 39-44 and 49, were the highest expressed genes among the L3 biased (Fig. 5C-E and Data S3B), and head-biased (Fig. 6, Data S4C and S8C-D). Sequences used for the phylogenetic analysis are provided in Data S9A-C.

**Figure 4.**
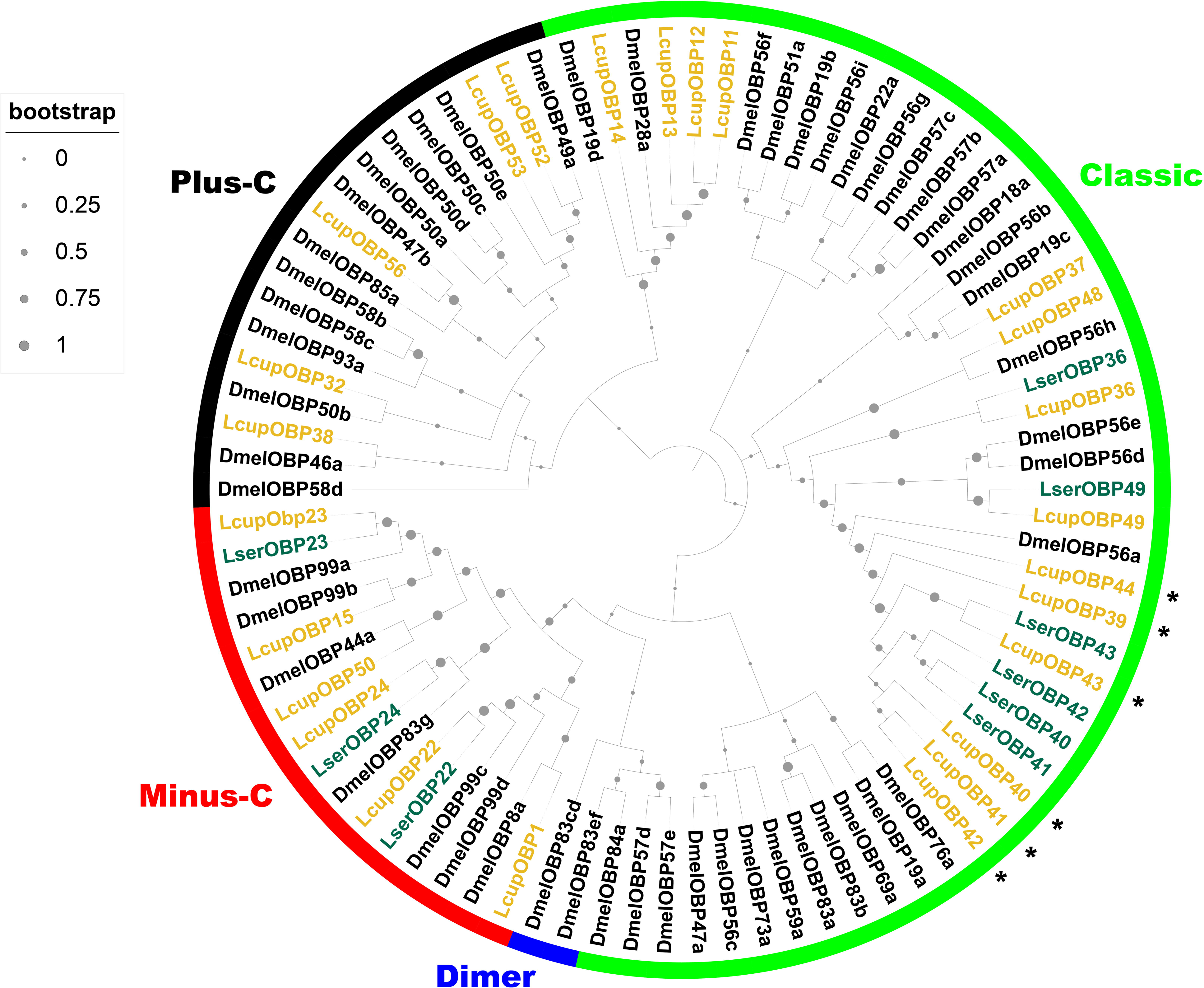
Phylogenetic analysis of the OBPs. Three Diptera species were used for the phylogenetic analysis as follows: *L. cuprina* (Lcup, gold), *L. sericata* (Lser, green), *D. melanogaster* (Dmel, black). OBP subfamilies classified according to the cysteine motif (Classic, Minus-C, Plus-C and Dimer), and node bootstrap supporting values were detailed. Only *L. cuprina* OBPs with a TPM expression ≥ 5 in larval head, gut or whole body were used for the analysis, but all *D. melanogaster* OBPs were included. In addition, OBPs previously identified as larval biased in *L. sericata* [30] were added to the analysis. If there was more than one isoform per gene, only isoforms producing a different amino acid sequence and showing a dissimilar transcript expression were added. Potential expanded *L. cuprina* OBPs 39 to 44 belonging to subfamily Classic were detailed with an asterisk. For more details about sequences used in the analysis refer to Data S9A-C.

**Figure 5.**
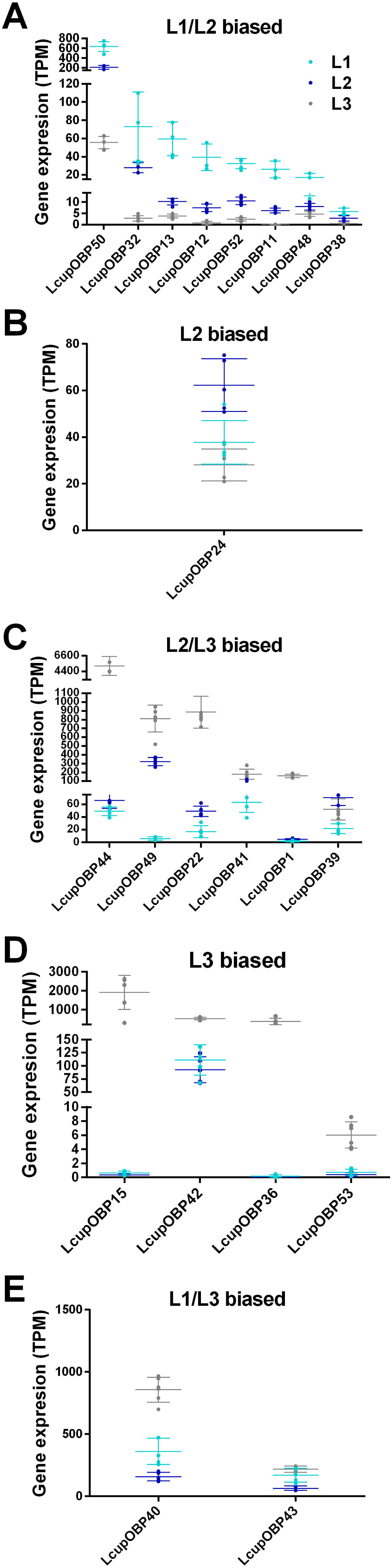
Expression and differential expression of the OBPs across *L. cuprina* larval stages. The TPM expression of OBPs was plotted clustering the same according to differential expression between the three larval stages as follows: (**A**) includes OBPs biased to the first two larval stages (L1 and L2) vs. L3; (**B**) shows OBPs only biased to the L2 stage; (**C**) groups OBPs biased to the last two stages (L2 and L3) vs. L1; (**D**) OBPs only biased to the last stage (L3); and (**E**) OBPs biased to the first and last stage (L1 and L3). Only receptors with a TPM expression ≥ 5 were added. Dots represent the gene TPM expression of each library; the average mean expression per gene, plus the SD between libraries were also plotted. For more details about the expression and differential expression of each receptor refer to Data S3B.

**Figure 6.**
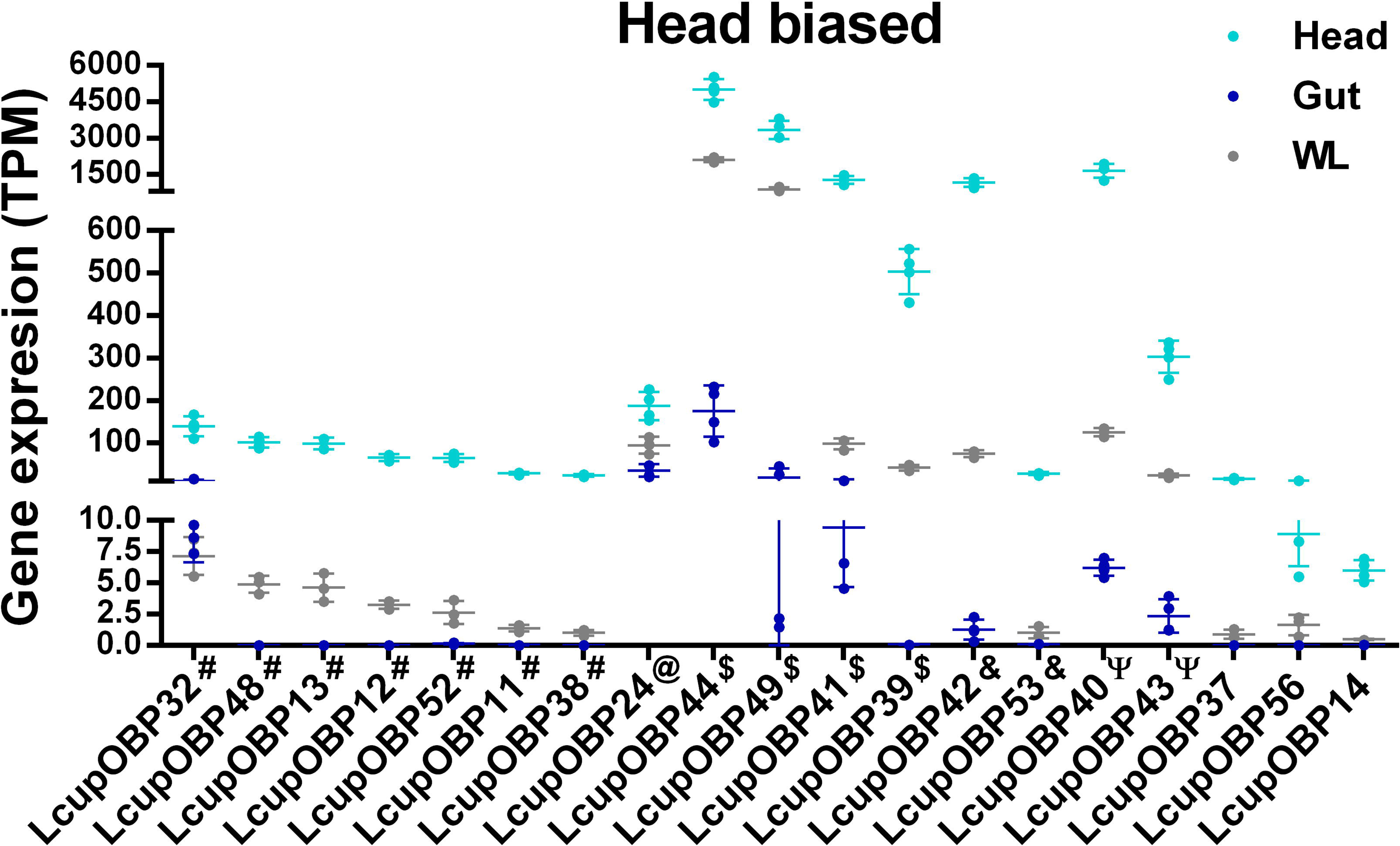
Expression and differential expression of larval head-biased OBPs. Only the TPM expression of head-biased OBPs was plotted. Dots represent the gene TPM expression of each library; the average mean expression per gene plus, the SD between libraries were also plotted. Only receptors with a TPM expression ≥ 5 were added. Symbols next to receptor’s names represent larval stage biased expression (plotted in Fig. 5) as follows: # L1/L2-biased; @ L2-biased; $ L2/L3-biased; & L3-biased; Ψ L1/L3-biased. For more details about the expression and differential expression of each receptor refer to Data S4C.

In insects, OBPs are also involved in lipid deposition on the cuticle [77], and some OBPs expressed at the MT and cuticle were associated with xenobiotics detoxification [78–80]. Based on their expression in the *L. cuprina* larvae, OBPs with a lipid biding affinity may be involved in energy homeostasis, avoiding larval desiccation and assisting detoxification. Remarkably, ABC transporters with similar functions, such as receptors from subfamilies G and H, shared a common expression pattern in the *L. cuprina* larvae. Part of the Minus-C and most of the Plus-C OBPs were L1/L2 biased vs. L3 (Fig. 5A-B and Data S3B). The Minus-C *LcupOBP50* and Plus-C *LcupOBP52* were among the most highly expressed sequences within these groups and biased to L1/L2 (Fig. 5A-B and Data S3B). Orthologs of these receptors, namely DmelOBPs 44a and 49a respectively, were studied in *D. melanogaster*. *DmelOBP44a* is involved in lipid binding [81] and *DmelOBP49a* associated with taste in the same species [82]. Further, Minus-C and Plus-C OBPs have been associated with binding of sesquiterpenes and derived sesquiterpenoids such as Juvenile Hormone (JH) in other insects [83–85], and they could have a similar role in *L. cuprina*. Only a few OBPs were expressed in the larval gut, and none of them biased to the same tissue (Data S4C). The Minus-C *LcupOBP22* and *LcupOBP24* were among the highest expressed in the gut (Data S4C). In insect, JH regulates the function of the adult and larval gut [86,87], and in *D. melanogaster* this hormone is synthesized in the *corpora allata* but also in the adult gut [86]. Consequently, the above OBPs could be involved with the JH-gut signaling system.

*Lcuppainless* was the only TRP above a 5 TPM expression threshold and biased to the larval head (Data S3C and S4D). In *D. melanogaster* this gene has been associated with taste, thermal sensing and proprioception [88–91]. Accurate temperature detection is indispensable to ensuring *L. cuprina* larval survival [17,19], and this receptor may be involved in thermal sensing. Of the CD36/SNMP genes, the scavenger receptor class B1 (SR-BI, XP_023294991.1) was upregulated in the L2/3 stages when compared to L1 (Data S3D), and the only member of this family showing a head bias (Data S4E). Interestingly this gene was highly expressed in the *L. cuprina* adult antenna [25], indicating a possible role in larval and adult olfaction. Other members of the same family, *e.g*. *protein croquemort-like* were biased to the larval gut (Data S4E). In *D. melanogaster* members of the CD36/SNMP gene family were associated with lipid binding and their expression in the gut of the same species [92] suggest a role in feeding and metabolism.

ML/NPC2 transporters have been associated with lipid binding, immune response and olfaction in different insect species [25,93–95]. Members of this family were highly expressed in the *L. cuprina* L2 stage and the larval gut (Data S3E and S4F). The *L. cuprina ecdysteroid-regulated 16 kDa* receptor (*LcupESR16*) was the only head-biased gene within the family (Data S4F). The same gene was the highest expressed in the *L. cuprina* adult antenna among this family [25]. Further, *ESR16* orthologs were found expressed in the antenna of many insect species [25,94,95] suggesting a possible role in olfaction.

Three CSPs were expressed in *L. cuprina* larvae, and two of them were biased to the head (Data S3F and S4G). In *Bradysia odoriphaga* Winnertz (Diptera: Sciaridae), a CSP protein was associated with sulfur-rich compounds detection [96], a common blowfly attractant [13]. It is possible that CSP proteins may have a role in detection of sulfur-rich compounds in *L. cuprina* larvae. The *ammonium transporter Rhesus type B* (*RHBG*) was biased to the *L. cuprina* L2 stage and the larval gut (Data S3G and S4H). This receptor is involved in ammonia detection [52,97] and excretion at the MT [98] in *D. melanogaster*. Ammonia-rich compounds are attractants to adult females of *L. cuprina* and *L. sericata* [13]. In adult antennae, *LcupRHBG* was highly expressed in both sexes with a female-bias [25]. In the larva of *L. sericata*, *LserRHBG* showed the highest expression in salivary glands followed by MT [30]. Consequently, *LcupRHBG* might be related to olfaction of adult flies, but involved in other processes in *L.cuprina* larvae, such as feeding and excretion.

#### Odorant receptors are not essential for L. cuprina larval response to rotten beef

To determine the importance of ORs in larval behavior, CRISPR/Cas9 was used to obtain a null mutation of the *LcupOrco* gene following previous protocols used in blowflies [63,99]. We identified a deletion mutation that introduced a premature stop codon at the end of the first exon, 265 nucleotides from the start codon (Fig. 7 and Figs. S2-S4, S7-S12). Without the presence of the odorant coreceptor (*Orco*), no odorant receptor can complete its olfactory function [54]. Larvae that were a mix of wild-type (*wt*), *LcupOrco* heterozygotes and homozygotes, were tested for diet preference in a choice assay. Overall, there was no major change in preference, with larvae preferring rotten beef that was at room temperature as previously reported [26] (Fig. 8A). No correlation was found between diet preferences and larval sizes, either for the genotyped (Fig. 8B) or non-genotyped (Fig. 8C) larvae, ANOVA, P = 0.22 and P = 0.25 respectively. This lack of correlation between genotype and diet preference was confirmed by a Chi-Square analysis, P = 0.66 (Fig. 8D and Table S2A). In the blowfly *C. hominivorax, ChomOrco* expression was minimal compared to adult flies showing the higher expression at the L1 stage [63], as observed here for *L. cuprina* larvae (Table 1). The authors were able to rear a *ChomOrco* silenced strain indicating that this gene may not be essential for *C. hominivorax* to complete the larval cycle. Our results suggest that *LcupOrco* is not necessary for *L. cuprina* larvae to respond to the complex mixture of odors emitted by decomposing beef, in contrast to observations in *D. melanogaster* larvae, where *Orco* is essential for the response to odors [33–36].

**Figure 7.**
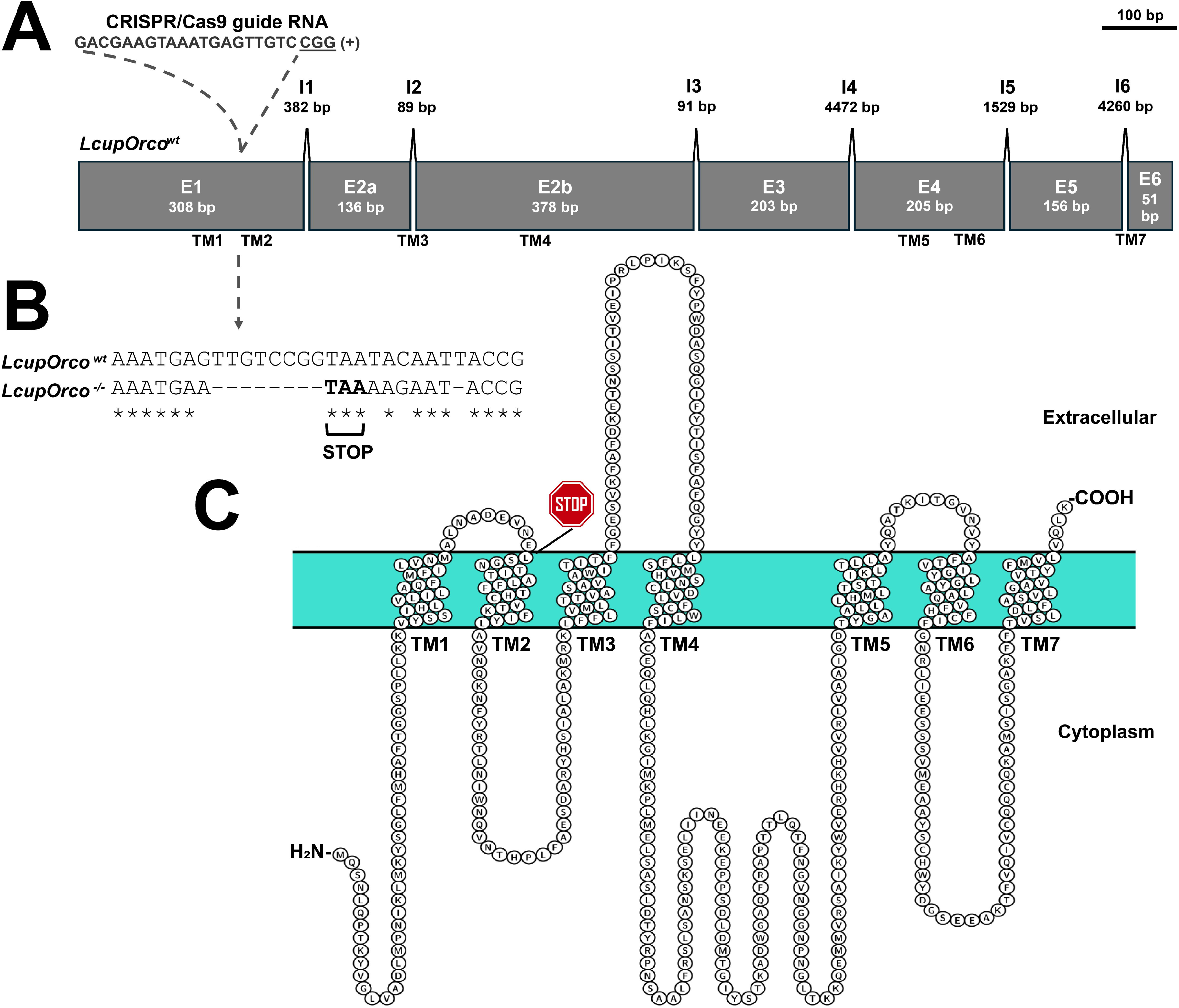
CRISPR/Cas9-mediated loss-of-function mutation of the *L. cuprina* odorant coreceptor (*LcupOrco*) gene. A deletion mutation in the first exon resulted in the introduction of a premature stop codon at the end of the first exon. (**A**) *LcupOrco* gene detailing exons (E) and introns (I) with lengths in base pairs (bp), the guide RNA target site, and PAM sequence (underlined); (**B**) alignment between the *wt* and mutated *LcupOrco* DNA sequences, detailing deletions and insertions, including a 8 bp gap that introduces a premature stop codon at 265 bp from the start codon, producing a 73 amino acids (aa) truncated protein of 478 aa for the *wt* protein; and (**C**) LcupORCO protein sequence detailing the premature stop codon position located after the glutamic acid (E) 73, the seven transmembrane (TM) regions and cytoplasmic and external loops. The interaction between Orco protein and any odorant receptor to produce a functional dimer occurs at the Orco C-term region (-COOH in the figure), which is absent in the mutated version of LcupOrco protein. For more details about *LcupOrco* silencing refer to Figs. S2-S4, S7-S12 and Notes S1 and S2.

**Figure 8.**
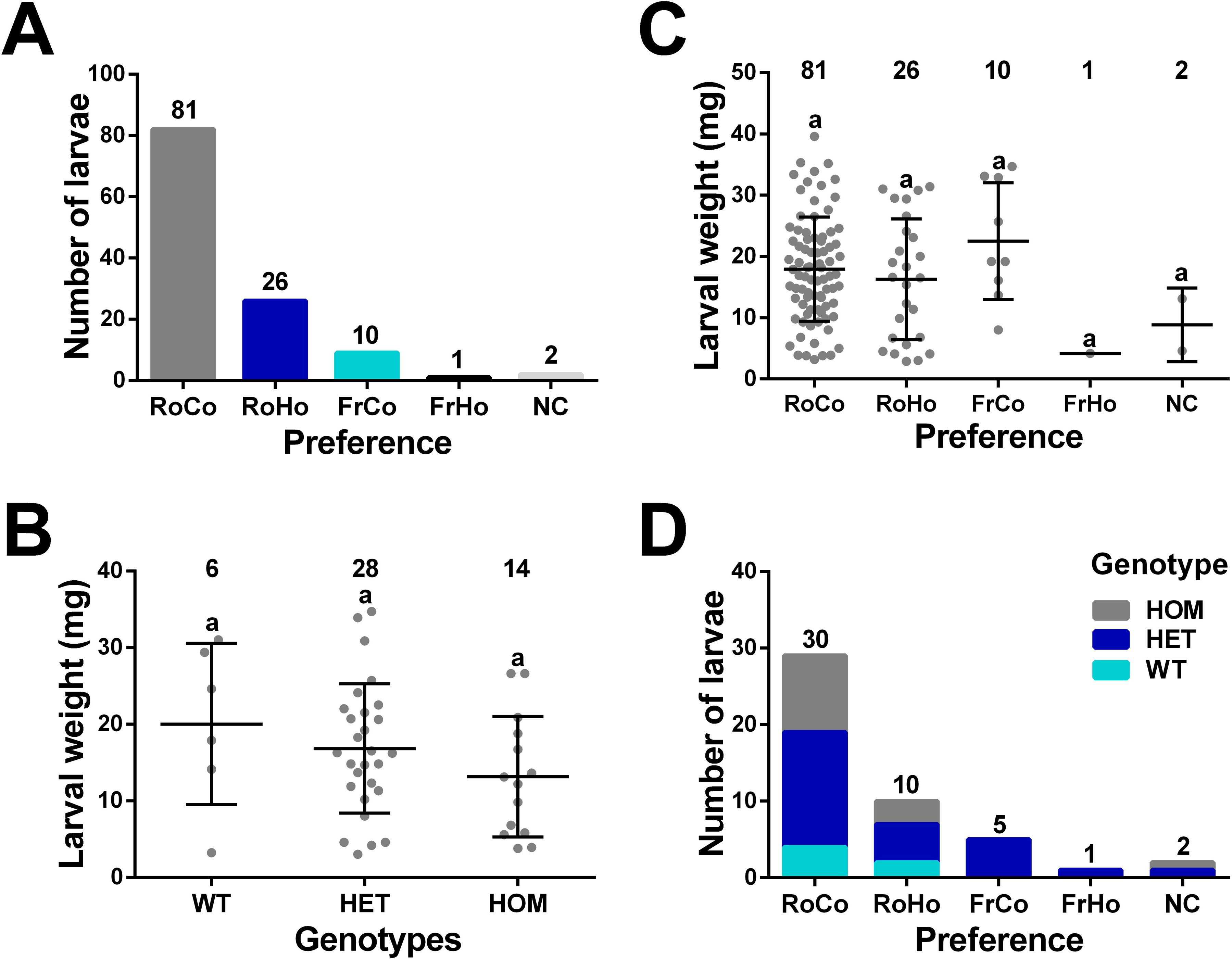
Odorant receptors are not essential for *L. cuprina* larval response to rotten beef. A larval choice assay was conducted to evaluate the larval response to fresh (Fr) and rotten (Ro) beef at two temperatures, cold 25±1°C (Co) and hot 33±1°C (Ho). A total of 120 early L3 (4 days after oviposition) were tested, and 48 larvae randomly collected from individuals attracted to each diet preference were genotyped. Larvae were obtained from crossing *LcupOrco* heterozygous flies giving an approximate 3:1 ratio of *LcupOrco* non-silenced (wild-type plus heterozygous, *wt* + HET) and silenced larvae for *LcupOrco* (homozygous, HOM). (**A**) the numbers on top of the bars represent total larvae attracted to each diet preference, *i.e*. rotten cold (RoCo), rotten hot (RoHo), fresh cold (FrCo), fresh hot (FrHo) and non-choice (NC); (**B**-**C**) show the weight of larvae in milligrams (mg) attracted to each diet preference, either for genotyped (**B**) or non-genotyped (**C**) larvae. Dots represent single larvae, numbers on top of each preference or genotype mean total number of larvae, and equal letters indicate no statistically significant difference (ANOVA, P = 0.22 for B and P = 0.25 for C); (**D**) shows how genotypes were randomly associated to each diet preference (Chi-Square analysis, P = 0.66). Numbers on top of the bars represent total larvae per diet preference. For more details about results obtained in this assay refer to Figs. 9A, 10A and Table S2A.

Disruption of *LcupOrco* affected the *L. cuprina* female oviposition behavior with egg laying significantly delayed compared to heterozygous siblings (Fig. S11A-G). This was not due to a feeding disorder as *LcupOrco*^-/-^ homozygotes showed a fully developed ovary (Fig. S12A-O). Further, a mating assay using transgenic males expressing a green fluorescent protein, showed that only *LcupOrco* heterozygous females laid eggs the first week after crossing (Fig. S11A). After a second week, *LcupOrco* silenced females (homozygous) started to lay non-fertilized eggs (Fig. S11B-C), which did not produce any offspring. *LcupOrco* silenced females needed a third week to start to lay fertilized eggs giving healthy progeny (Fig. S11D-G). Notably, *LcupOrco* silenced males showed an average fertility of 88% including 30 males tested across four generations. These results were in line with previous observations in *Harpegnathos saltator* T. C. Jerdon, 1851 (Hymenoptera: Formicidae), where silencing of *Orco* produced males with a normal mating behavior and females with a delayed and decreased oviposition [100]. However, silencing of *Orco* in *Helicoverpa armigera* Hübner, 1808 (Lepidoptera: Noctuidae), produced the opposite phenotype, *i.e*. sterile males and fertile females [101], indicating that the function of *Orco* may change among species.

We next asked if *LcupOrco* is important for detecting rotten beef in adult females using a spatial olfactometer. A mix of *wt*, *LcupOrco* heterozygotes and homozygotes was obtained from crossing *LcupOrco* heterozygous flies. This mix included an approximate 3:1 ratio of *LcupOrco* non-silenced to silenced females. Also, a different group of *L. cuprina wt* females were used for a further comparison with *LcupOrco* females. As previously reported [25], *wt* females were more attracted to the rotten beef vs. fresh beef, or did not respond to any of the stimulus (non-choice, NC), ANOVA, P = 0.0001 (Fig. 9A). However, for the *LcupOrco* mix, the number of non-responding (NC) females was higher than any of other two preferences, ANOVA, P = 0.0001 (Fig. 9B). Genotyping of randomly selected females from tested groups, showed that *LcupOrco* silenced females were overrepresented among the non-responding flies (NC), Chi-Square, P = 0.0001 (Fig. 9C and Table S2B). These observations showed that *LcupOrco* is essential for odor detection in adult females, and our results are consistent with previous observations in *C. hominivorax*, where disruption of *ChomOrco* impaired the foraging and host-seeking behaviors of adult flies [63].

**Figure 9.**
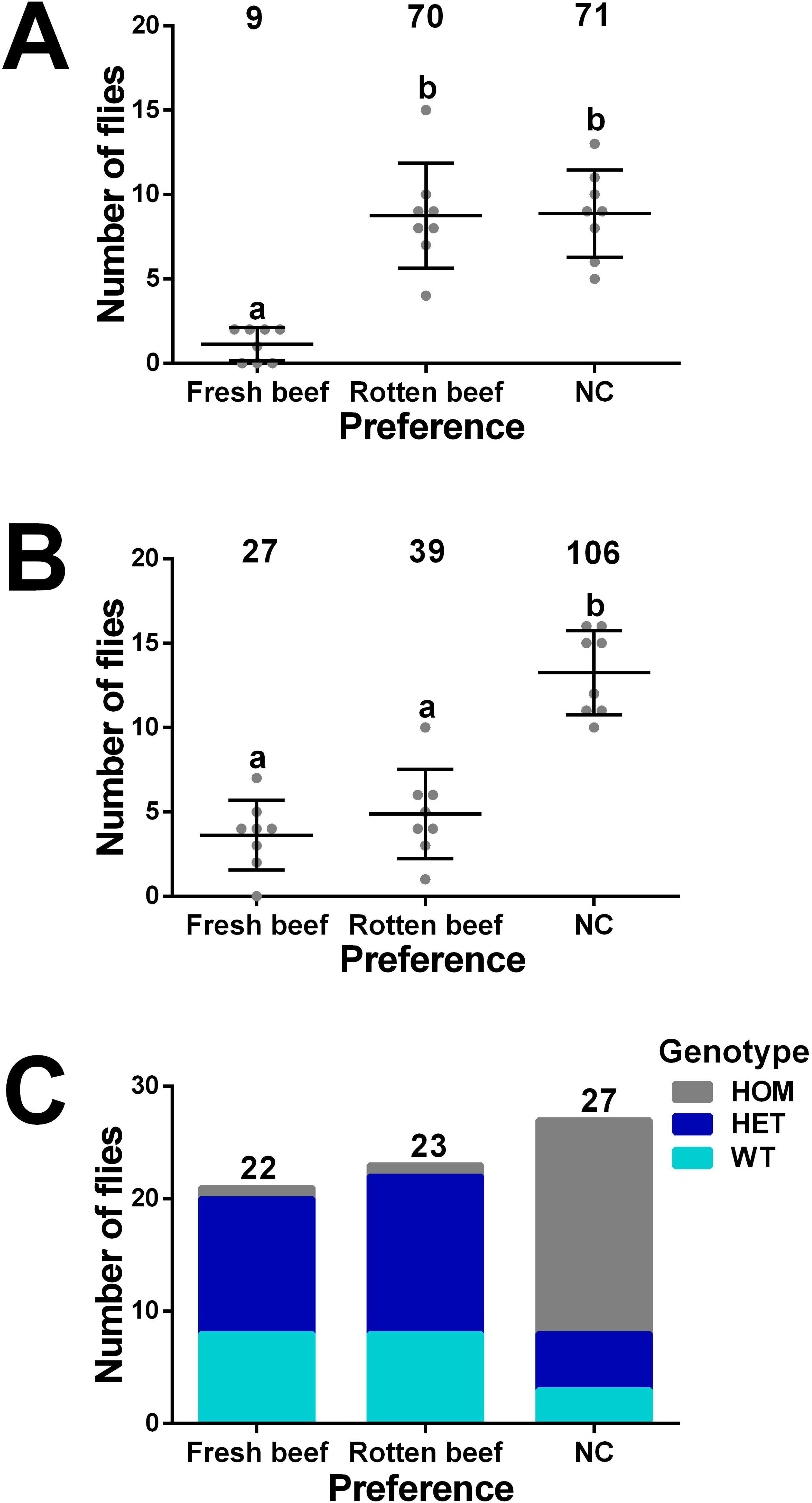
Disruption of *LcupOrco* impairs *L. cuprina* adult female olfaction. Females were obtained from crossing *LcupOrco* heterozygous flies giving an approximate 3:1 ratio of *LcupOrco* non-silenced (wild-type plus heterozygous, *wt* + HET) and silenced females for *LcupOrco* (homozygous, HOM). A different group of *L. cuprina wt* females were used for a further comparison with *LcupOrco* females. All flies used for the olfaction assay were ∼ 10-day-old gravid females and were separated from males the day before the assay using CO_2_ to anesthetize the flies. The attractants used to lure the flies were 2 grams of fresh (Fr) and rotten (Ro) beef at room temperature. A total of 80 *wt* and 174 *LcupOrco* females were tested, and 72 of the latter were genotyped. (**A**) the *wt* females were more attracted to the rotten beef vs. fresh beef, or did not respond to any of the stimulus (NC), ANOVA, P = 0.0001; (**B**) in females belonging to the *LcupOrco* mix, the number of non-responding (NC) females was higher than any of other two preferences, ANOVA, P = 0.0001; (**C**) genotyping of randomly selected females from tested groups, showed that *LcupOrco* silenced females were overrepresented among the non-responding flies (NC), Chi-Square, P = 0.0001. Different letters indicate statistically significant difference. For more details about results obtained in this assay refer to Figs. S9B, S10B and Table S2B.

#### Non-sensory genes expressed and biased to the larval head

Transcripts encoding the *L. cuprina* orthologs of the *D. melanogaster yellow*, *yellow-h* and *ebony* genes, with roles in cuticle pigmentation and hardening (tanning) [102–104], were among the highly expressed head-biased genes vs. the whole body (Data S7A, S8C-D). The *L. cuprina eclosion hormone* gene (*LcupEH*) was also enriched in the larval head in comparison to the whole body (Data S7A). In insects, *EH* is associated with cuticle tanning through the control of *Bursicon* (*bur*) and *Partner of Bursicon* (*pbur*) [105]. The presence of these transcripts suggests a role in cuticle and/or mouthparts tanning of the *L. cuprina* larvae. In *D. melanogaster*, *yellow* controls the tanning of the larval mouth parts [106]. Remarkably, the *tan* gene which display an antagonistic role to *ebony* in cuticle tanning in *D. melanogaster* [102,107] was downregulated in the head vs. the whole body (Data S8D).

Transcripts encoding for mucins such as *Lcupmucin-5AC* were highly expressed in the head compared to whole body, and head-biased (Data S7A and S8C). Mucins have been studied in the insect gut, playing a role in food processing and supporting the immune system [108]. Enrichment of mucin transcripts in the larval head could suggest a role at the anterior part of the digestive system. We previously found that members of this gene family were highly expressed in the *L. cuprina* adult antenna [25], and suggested that mucins may be important for olfaction, a role confirmed in vertebrates [109] but not yet addressed in insects. It is possible that mucins may also be important for olfaction in *L. cuprina* larvae.

#### Genes potentially associated with L. cuprina larva feeding and digestion

Proteases and lipases such as *trypsin delta* and *lipase 3* respectively, were among the genes highly enriched in the larval gut compared to whole body and biased to the larval gut of *L. cuprina* (Data S7B and S8A-B). These findings are consistent with a larval diet composed mainly of meat with a small fat content [76]. Neuropeptides *Allastostanine*, *CCHamide-2* and *long neuropeptide F* (*NPF*), that modulate food intake and digestion in *D. melanogaster* [110,111], were also expressed and biased to the larval gut (Data S7B and S8A-B). Neuropeptide *Diuretic hormone class 2* (*DH31*), which play a role in diuresis, gut contractions and elimination of virulent bacteria in *D. melanogaster* [112] showed the same expression pattern as the above neuropeptides (Data S7B and S8A-B). In addition, transcripts encoding for mucins and cytochrome P450s (CYPs) were overrepresented in the larval gut (Data S7B and S8A-B). CYPs play a crucial role in the insect gut metabolizing endogenous and exogenous substances [113], and mucins in the aggregation of harmful elements [108]. These enzymes may be involved together in detoxifying harmful compounds and metabolizing dietary components in the gut of *L. cuprina* larva.

#### Gene expression and differential expression across larval stages

The most highly expressed sequences in the whole larva of any stage were genes associated with cuticle (*e.g*. *larval cuticle-protein 65Ag1-like, Lcuplcp65Ag1*), muscle formation (*e.g*. *myosin-regulatory light chain 2*), and RNA translation factors (Data S1A-C). These transcripts were also abundant in larval head, but with an overrepresentation of sequences coding for cuticle proteins and sensory receptors (Data S2A). The larval cuticle is essential for larval life, acting as both skin and exoskeleton, with the muscles linked directly to the same [114], and the interaction between cuticle and muscles allows larval locomotion [114]. The cuticle also protects the larva from desiccation and abrasion, includes sensory and feeding organs and constitutes the first barrier of the immune system [115].

Genes associated with cuticle formation, *e.g*. *Lcuplcp65Ag1*, showed the highest fold-change (FC) increases in L2 vs. L1 (Data S10A). Other genes potentially associated with cuticle tanning such as the orthologs of the *D. melanogaster yellow, yellow-h* and *EH* genes were also upregulated in L2 vs. L1 (Data S10A). An ortholog of the *Adipokinetic hormone* (*AKH*), which is involved in mobilizing lipids from storage tissues in different insects [116], showed a high FC increase in L2 vs. L1 (Data S10A). Other genes involved in protein and fat digestion, such as *Lcuptrypsin-1* and *Lcuplipase 3* respectively, were upregulated in the L2 stage (Data S10A). Interestingly, orthologs of the antimicrobial peptides *Lucifensin* [117] and *defense protein-l(2)34Fc* (*l(2)34Fc*) [118], were also increased in L2 vs. L1 (Data S10A). Larvae are reared on fresh meat under non-sterile conditions [9] and thus could be exposed to a higher bacterial load at later stages. On the contrary, the expression of orthologs of feeding modulators such as *short neuropeptide F* (*sNPF*) and *SIFamide* [119,120] was downregulated in L2 vs. L1 (Data S10A). In *D. melanogaster* larvae, *sNPF* increases food intake but does not prolong the feeding period or modulate food preferences, unlike *NPF* [120]. Both, *NPF* and *sNPF* are major modulators of feeding and antimicrobial immune response [121,122].

In the late L3, arylphorin genes showed the highest FC increases compared to previous larval stages (Data S10B-C). In insects, these proteins have a role in storage and cuticle formation during the prepupal period [123]. Other genes associated with cuticle formation, such as *Lcuplcp65Ag1*, were also upregulated in L3 (Data S10B-C). Genes potentially involved in larval energy homeostasis such as *LcupAKH* and *LcupCCHAmide-2*, also showed a biased expression to L3 (Data S10B-C). In contrast, genes potentially involved in cuticle tanning of *L. cuprina* larva such as *Lcupyellow* and *LcupEH*, and orthologs of feeding modulators such as *sNPF*, *NPF* and *SIFamide* were downregulated in L3 vs. L1/L2 (Data S10B-C). In addition, orthologs of the antimicrobial peptides *Lucifensin* and *l(2)34Fc*, and chymotrypsin inhibitors associated with protease inactivation [124] were also downregulated in L3 vs. L1/L2 (Data S10B-C).

## Conclusions

Based on their expression patterns, ORs, GRs, IRs and PPKs may not play a central role in the *L. cuprina* larval sensory signaling and digestive systems, but further studies are needed to confirm this observation. In contrast, ABCs were highly enriched in head and gut RNA and OBPs only in the head, indicating that both gene families are essential to the physiology of the *L. cuprina* larva. Notably, ABCs were also highly enriched in the *L. cuprina* adult antenna [25], suggesting a role in olfaction across all *L. cuprina* stages for this gene family. Behavioral assays showed that ORs were essential for the adult female response to rotten beef, but not for the larval response. These findings are consistent with high levels of expression of *LcupOrco* in the adult female antenna [25] but very low expression in larvae (present work). Among other relevant genes in *L. cuprina* larvae, the TRP *Lcuppainless* may play a role in environmental sensing such as thermal, taste or proprioception, and the ammonia transporter *LcupRHBG* in food digestion and excretion. Future studies using gene editing techniques in combination with behavioral assays will be conducted to assess the function of the genes highlighted in the present study.

## Availability of data and materials

All relevant data are in the manuscript. Raw sequences were uploaded to the NCBI database under Bioproject PRJNA1195419.

## Abbreviations

ABC: ATP-binding cassette transporter
AKH: Adipokinetic hormone
AMT: ammonium transporter
BH: Benjamini-Hochberg
BLAST: basic local alignment search tool
C: cysteine
CDS: coding sequence
CD36/SNMP: CD36 and sensory neuron membrane proteins
CSP: chemosensory protein
Co: cold
CYP: cytochrome P450
DE: differential expressed
DGE: differential gene expression
DH3: diuretic hormone class 2
DNA: deoxyribonucleic acid
DMDS: dimethyl disulfide
DMTS: dimethyl trisulfide
E: exon
EH: eclosion hormone
FB: fat body
FC: Fold-Change
Fr: fresh
G: gut
g: gram
gDNA: genomic DNA
GR: gustatory receptor
H: head
h: hour
Ho: hot
hPa: hectopascal
I: intron
IPM: integrated pest management
IR: ionotropic receptor
iTOL: interactive Tree of Life
JH: juvenile hormone
KB: kilobase
KCl: potassium chloride
L: larva
LS: left side
ML: Maximum likelihood
ML/NPC2: Myeloid lipid-recognition (ML) and Niemann-Pick C2 disease proteins
mRNA: messenger RNA
MT: malpighian tubules
NCBI: national center for biotechnology information
NC: non-choice
NF: nuclease free
NJ: Neighbor-Joining
NPF: long neuropeptide F
OBP: odorant-binding protein
OR: odorant receptor
*Orco*: odorant receptor coreceptor
PCA: principal components analysis
PE: paired end
PPK: pickpocket receptor
RH: relative humidity
RHBG: ammonium transporter Rhesus type B
RNA: Ribonucleic Acid
RNA-Seq: RNA-sequencing
Ro: rotten
RS: right side
RT: room temperature
SD: standard deviation
SE: secretions and excretions
SIT: sterile insect technique
sNPF: short neuropeptide F
SRB1: Scavenger Receptor Class B Type I
TM: transmembrane
TPM: transcripts per million
TRP: Transient receptor potential
UTR: untranslated
WL: whole larva
*wt*: wild-type

## Declarations

### Ethics approval and consent to participate

Not applicable.

### Consent for publication

Not applicable.

### Author contributions

MJS and JPW conceptualized the experiments. JPW and RKL collected the samples and performed the experiments, JPW completed the analyses and produced the figures. JPW and MJS wrote the manuscript and RKL reviewed and edited the manuscript. All authors read and approved the final manuscript.

## Supplementary information

**Additional file 1: Fig. S1:** larval stages and tissues used for the RNA-Seq experiment. **(A)** whole larva stage 1 (L1); **(B)** whole larva stage 2 (L2); **(C)** lateral view of a whole late larva stage 3 (L3) and **(D)** ventral view; **(E)** whole early larval stage 3 (WL); **(F)** first segment designated as “head” (H) from an early L3, detailed within a dashed white circle; and **(G)** gut from an early L3, where crop have been separated from the cardia originally connected by the foregut (not present in the picture); **Note S1:** *LcupOrco* gene, guide RNA (gRNA) sequence, and primers used to determine potential indels and point mutations at the gRNA cutting site (genotyping); **Fig. S2:** *LcupOrco* insert construction. **(A)** *LcupOrco* gene fragment including two sequences corresponding to the left (LHA) and right (RHA) homology arms of 1000 bp in length each one, necessary for the CRISPR homology directed repair (HDR) protocol cloned using plasmid pUCIDT-AMP GoldenGate (ordered from IDT). In addition, a DNA section including the restriction cutting sites (RCS) for enzymes XhoI (NEB) and NotI (NEB) was added in between of both sequences in the same plasmid; **(B)** donor plasmid including the ZsGreen Marker followed by the *Hsp83* promoter located between positions 1891 to 7441 [2]; **(C)** ZsGreen Marker obtained from the donor plasmid. The plasmid was linearized using restriction enzymes BpmI (NEB), XhoI and NotI at 37°C for 60 min and the reaction was finished at 65°C for 20 min. After restriction digestion the mix was electrophoresed in a 1.5% agarose gel at 60 volts for 120 min. The band with the size corresponding to the marker (∼5 Kb) was recovered from the gel and purified using the Zymoclean Gel DNA Recovery Kit (Zymo) following the manufacturer’s specifications; **(D)** to obtain the final construct, 50 ng of linearized pUCIDT-AMP GoldenGate plasmid + 175.5 ng of the marker fragment (1:3 ratio) were ligated overnight (O/N) at 16°C using the T4 DNA Ligase (NEB). After ligation, 1 μl of the reaction mix was used to transform 10-beta competent *Escherichia coli* cells (NEB) following the manufacturer’s specifications. Seventy-five μl of undiluted transformed 10-beta cells were poured into an Ampicillin 20 mg/ml (+Amp) Lysogeny Broth (LB) plate and incubated O/N at 37°C; **(E)** ten clones were recovered from the LB plate and grow in 3 mL of liquid LB +Amp O/N at 37°C and 250 rpm. To obtain the plasmids from clones the ZR Plasmid Miniprep Kit (Zymo) was used following the manufacturer’s specifications and evaluated with the BpmI, XhoI and NotI restriction enzymes following the same protocols described above for restriction digestion and electrophoresis; **(F)** three positive plasmids for the selected restriction cutting sites were sequenced using Oxford Nanopore and aligned to the *in-silico* construct to confirm the sequence identity. Abbreviations: Kb = kilobase; MW = molecular weight; **Fig. S3:** containers used for fly rearing and crosses. **(A)** plastic bottle used for fly rearing; **(B)** sandglass shaped arrangement made using two five ounces clear-plastic cups joined from removed bottoms used for fly crosses after embryo injections. These containers also included: a top holed cap and a bottom cap with an opening of ∼ 1 inch in diameter; a rounded piece of white paper towel; a small glass vial filled with tap water and a small protein cookie made with yeast, milk and egg powder and cane sugar; **Fig. S4:** fly crosses completed after embryos microinjections. Abbreviations: G = generation; *wt* = wild-type; **Note S2:** primers used for *LcupOrco* insert detection of landing site. Three types of pair of primer associated with three different strategies were used to determine if the landing site of the *LcupOrco* insert was withing the *LcupOrco* locus in the *L. cuprina* genome. First one was associated with the left side (LS) of the insert including the upstream genome region, the *Hsp83* promoter and part of the green marker; the second one was designed to amplify the right side (RS) of the insert including the part of Tub3’-UTR and the downstream genome region; and third one was to amplify the whole *LcupOrco* insert (All) including from upstream to downstream *L. cuprina* genome regions flanking the *LcupOrco* locus; **Table S1:** RNA-Seq data overview. **(A)** each sample represents an RNA-Seq library. Sample’s names, description and number of raw, trimmed and mapped reads per library are provided; **(B)** total sequences identified for the RNA-Seq experiment, divided by RNA type; **Fig. S5:** principal component analysis (PCA) completed using transcripts with a TPM expression ≥ 5 for all libraries (samples) associated with EXP-1 (larval stages). Samples WL1_1 to 3, WL2_1 to 3 and WL3_1 to 6 were collected from the LA07 colony. Samples WL1_4 and 5 and WL2_4 and 5 were collected from heterozygous individuals obtained from the crossing between the LA07 and the SLAM5X colonies. The PCA analysis did not show differences between samples of the same larval stage, collected from different colonies. Abbreviations: L1 = first larval state; L2, second larval stage; L3 = third larval stage; WL = whole larva sample; TPM = transcript per million; **Fig. S6:** ABC transporters preliminary Neighbor Joining phylogenetic analysis. Abbreviations: Dmel = *Drosophila melanogaster*; Lcup = *Lucilia cuprina*; Lser = *Lucilia sericata*; **Fig. S7:** guide RNA (gRNA) cutting efficiency tested *in vitro* using the EnGen Spy Cas9. The amplicon size of *LcupOrco* gene fragment without cutting was 439 bp and after Cas9 *in vitro* cutting generated two overlapped amplicons of 224 and 215 bp, respectively. Abbreviations: C = control, uncut genomic DNA; MW = molecular weight; T = treated, gDNA + Cas9; **Fig. S8:** *Lucilia cuprina* eggs, larva stage 3 (L3) and adult flies expressing the ZsGreen marker before the genotyping analysis to determine *LcupOrco* insert landing site. **(A)** L3 showing transient expression of the marker 7 days after *wt* eggs injection; **(B)** eggs obtained from crossing a *LcupOrco* mosaic male with a *wt* female; **(C)** *wt* L3 on top vs. a heterozygous ZsGreen L3 at the bottom; **(D)** heterozygous ZsGreen L3 on top vs. homozygous for the same marker at the bottom; **(E)** dorsal view of a *wt* adult male (left) vs. G1 heterozygous ZsGreen male (right) under bright field; **(F)** same males showed in E under a green filter; ventral view of the same males using bright field **(G)** and a green filter **(H)**; **Fig. S9:** PCR amplification of gDNA of samples collected from *LcupOrco* larval diet preference test and female olfaction assay. **(A)** samples of larval olfaction assay; **(B)** samples of adult female olfaction assay. Primers, amplicon sequence and cycling parameters are detailed in **Note S1** (see Supplementary Materials and Methods section). Abbreviations: MW = molecular weight; **Fig. S10:** Sanger sequencing results for samples collected from *LcupOrco* larval diet preference test and adult female olfaction assay. (**A**) genotyped samples from # 1 to 48 of larval diet preference test; and (**B**) genotyped samples from # 1 to 72 of adult female olfaction assay. Synthego software was used to analyze sample’s chromatograms and indels % = 0 corresponds to *wt* samples, from 1 to 89% to heterozygous samples, and ≥ 90% to homozygous samples. Results were compiled along with single larva preferences in **Table S2 A-B**; **Table S2:** compiled data of samples associated with *LcupOrco* behavioral assays. **(A)** larval diet preference test; **(B)** adult female olfaction assay. The numbers one through eight after the letters in table B, refer to the main chambers of the olfactometer and numbers after dots, to single females from the same chamber. Abbreviations: Fr = fresh beef (room temperature); FrCo = fresh cold beef; FrHo = fresh hot beef; HET = heterozygous; HOM = homozygous; NC = non-choice; NG = non-genotyped; Ro = rotten beef (room temperature); RoCo = rotten cold beef (25±1°C); RoHo = rotten hot beef (33±1°C); *wt* = wild-type. Hyphen symbol means non-tested by Sanger sequencing; **Fig. S11:** eggs obtained from *LcupOrco* mutated females. **(A)** eggs obtained from 8-day-old *LcupOrco^+/-^* females 3 days after crossing them to *NPF^-/-^* males expressing the ZsGreen marker. The same marker was removed from *LcupOrco* females after confirming that the *LcupOrco* insert landing site was not located withing the *LcupOrco* gene locus in the *L. cuprina* genome. The insert was removed by selecting non-fluorescent larvae and crossing males obtained from them for two generations vs. *wt* females. In addition, males from each generation were genotyped using protocols described in **Note S1** (see Supplementary Materials and Methods section) to confirm the presence of indels and point mutations within the *LcupOrco* coding region. The fluorescence in eggs confirmed the mating between the *LcupOrco^+/-^* females and *LcupNPF^-/-^* males and egg fertilization. *LcupOrco^-/-^*females did not lay eggs at this time; **(B)** part of *LcupOrco^-/-^*females laid eggs 10 days after mixing them with *LcupNPF^-/-^* males but eggs were not fertilized as showed under green filter **(C)**, and no larvae emerged from them; part of the *LcupOrco^-/-^* females laid fertilized eggs 17 days after mixing with *LcupNPF^-/-^* males, as showed under bright field **(D-F)** and green filter **(E-G)**; these eggs produced healthy progeny; **Fig. S12:** ovaries from 10-day-old *L. cuprina wt* and *LcupOrco* mutated females. Three females per condition were dissected (columns). All females were provided with tap water and sugar, but the presence of protein in the diet and mating condition changed between groups; *wt* virgin females not fed **(A-C)** and fed with protein **(D-F)**; *wt* mated females not fed **(G-I)** and fed with protein **(J-L)**; *LcupOrco-/-* mutated females fed with protein **(M-O)**.

**Additional file 2: Data S1. TPM transcript expression for *L. cuprina* larval stages. A** to **C** compile all libraries from each stage, namely L1, 2 and 3, showing the average expression for each transcript, only for transcripts with a TPM expression ≥ 5. **D** to **S** show the expression of all transcripts for all libraries.

**Additional file 3: Data S2. TPM transcript expression for *L. cuprina* early L3 tissues. A** to **C** compile all libraries for the larval head, gut and whole larva, respectively. Same tabs also show the average expression for each transcript, only for transcripts with a TPM expression ≥ 5. **D** to **N** show the expression of all transcripts for all libraries.

**Additional file 4: Data S3. Sensory receptors expressed in *L. cuprina* larval stages. A**) ATP-binding cassette (ABC); **B**) odorant-binding protein (OBP); **C**) transient receptor potential channel (TRP); **D**) CD36-sensory neuron membrane proteins (CD36/SNMPs); **E**) ML (MD-2 (myeloid differentiation factor-2)-related Lipid-recognition and Niemann-Pick C2 disease proteins (ML/NPC2); **F**) chemosensory protein (CSP); **G**) ammonia transporter (AMT). Only receptors with a TPM expression ≥ 5 are displayed. Differential expressions between stages for each receptor were listed.

**Additional file 5: Data S4. Sensory receptors expressed in *L. cuprina* early L3 tissues. A**) gustatory receptor (GR); **B**) ATP-binding cassette (ABC); **C**) odorant-binding protein (OBP); **D**) transient receptor potential channel (TRP); **E**) CD36-sensory neuron membrane proteins (CD36/SNMPs); **F**) ML (MD-2 (myeloid differentiation factor-2)-related Lipid-recognition and Niemann-Pick C2 disease proteins (ML/NPC2); **G**) chemosensory protein (CSP); **H**) ammonia transporter (AMT). Only receptors with a TPM expression ≥ 5 are displayed. Differential expressions between stages for each receptor were listed.

**Additional file 6: Data S5. ABC protein sequences included in the phylogenetic analysis. A**) *D. melanogaster*; **B**) *L. cuprina*; **C**) *L. cuprina* receptors with a TPM expression ≥ 5; **D**) larval biased *L. sericata* ABC transporters vs. adult flies.

**Additional file 7: Data S6. TPM transcript expression and differential expression of the ABC transporter LcupABCC13_Sur and LcupABCG6.** Annotation of both receptors was corrected using assembly GCA_001187945.1, and the transcript expression and differential expression between A) larval stages and B) tissues was calculated.

**Additional file 8: Data S7. Transcripts enriched in A) the larval head or B) larval gut, considering only TPM expression ≥ 5.**

**Additional file 9: Data S8. Differential expressed (DE) transcripts between different *L. cuprina* early L3 tissues.** Head and gut biased transcripts with a TPM expression ≥ 5 are provided in tabs **A** and **C**. All DE expressed transcripts between the head or the gut vs. the whole larva are displayed in tabs **B** and **D**, respectively.

**Additional file 10: Data S9. OBP protein sequences included in the phylogenetic analysis. A**) *D. melanogaster*; **B**) *L. cuprina* receptors with a TPM expression ≥ 5; **C**) larval biased *L. sericata* OBPs vs. adult flies.

**Additional file 11: Data S10. Differential expressed (DE) transcripts between different *L. cuprina* larval stages. A** to **C** compile those DE transcripts for L2 vs. L1, L3 vs. L2 and L3 vs. L1, respectively. Only transcripts with a TPM expression ≥ 5 are displayed. From **D** to **N** same groups comparisons are shown but including all DE transcripts.

