## Supplementary Information for "Genetic and behavioral analyses suggest that larval and adult stages of *Lucilia cuprina* employ different sensory systems to detect rotten beef"

### Supplementary Results

**Supplementary Table 1: RNA-Seq data overview.** (**A**) each sample represents an RNA-Seq library. Sample's names, description and number of raw, trimmed and mapped reads per library are provided; (**B**) total sequences identified for the RNA-Seq experiment, divided by RNA type.

**A**

| Sample | Description | Input (raw reads) | Seqs after trimming | Seqs that mapped to the reference genome | % of mapped reads |
|---|---|---|---|---|---|
| L1_1 | Paired reads created from L1_1_R1 + L1_1_R2 | 52,830,504 | 52,205,318 | 48,807,734 | 93.5 |
| L1_2 | Paired reads created from L1_2_R1 + L1_2_R2 | 48,086,438 | 47,506,322 | 42,573,624 | 89.6 |
| L1_3 | Paired reads created from L1_3_R1 + L1_3_R2 | 42,877,238 | 42,330,090 | 37,676,550 | 89.0 |
| L1_4 | Paired reads created from L1_4_R1 + L1_4_R2 | 45,603,866 | 45,053,754 | 40,935,156 | 90.9 |
| L1_5 | Paired reads created from L1_5_R1 + L1_5_R2 | 41,328,256 | 40,821,682 | 37,217,254 | 91.2 |
| L2_1 | Paired reads created from L2_1_R1 + L2_1_R2 | 58,320,992 | 57,698,612 | 51,844,520 | 89.9 |
| L2_2 | Paired reads created from L2_2_R1 + L2_2_R2 | 41,450,914 | 40,964,162 | 36,594,770 | 89.3 |
| L2_3 | Paired reads created from L2_3_R1 + L2_3_R2 | 53,656,602 | 53,010,010 | 47,184,164 | 89.0 |
| L2_4 | Paired reads created from L2_4_R1 + L2_4_R2 | 42,815,302 | 42,325,130 | 38,215,588 | 90.3 |
| L2_5 | Paired reads created from L2_5_R1 + L2_5_R2 | 41,283,070 | 40,828,438 | 36,433,156 | 89.2 |
| L3_1 | Paired reads created from L3_1_R1 + L3_1_R2 | 49,724,376 | 49,257,306 | 43,834,497 | 89.0 |
| L3_2 | Paired reads created from L3_2_R1 + L3_2_R2 | 46,915,790 | 46,417,644 | 39,777,785 | 85.7 |
| L3_3 | Paired reads created from L3_3_R1 + L3_3_R2 | 47,115,312 | 46,591,974 | 42,029,106 | 90.2 |
| L3_4 | Paired reads created from L3_4_R1 + L3_4_R2 | 50,676,690 | 50,162,324 | 43,686,831 | 87.1 |
| L3_5 | Paired reads created from L3_5_R1 + L3_5_R2 | 48,385,720 | 47,878,954 | 41,611,198 | 86.9 |
| L3_6 | Paired reads created from L3_6_R1 + L3_6_R2 | 40,037,226 | 39,576,554 | 34,283,222 | 86.6 |
| G1 | Paired reads created from G1_R1 + G1_R2 | 376,305,232 | 363,255,256 | 330,378,422 | 90.9 |
| G2 | Paired reads created from G2_R1 + G2_R2 | 426,594,950 | 413,197,846 | 375,863,416 | 91.0 |
| G3 | Paired reads created from G3_R1 + G3_R2 | 389,811,044 | 376,623,516 | 342,584,417 | 91.0 |
| G4 | Paired reads created from G4_R1 + G4_R2 | 418,659,816 | 405,492,620 | 369,667,402 | 91.2 |
| H1 | Paired reads created from H1_R1 + H1_R2 | 368,202,822 | 356,051,540 | 327,903,784 | 92.1 |
| H2 | Paired reads created from H2_R1 + H2_R2 | 266,154,798 | 256,836,086 | 235,959,629 | 91.9 |
| H3 | Paired reads created from H3_R1 + H3_R2 | 421,364,048 | 405,731,648 | 373,815,153 | 92.1 |
| H4 | Paired reads created from H4_R1 + H4_R2 | 358,563,270 | 345,286,086 | 316,903,040 | 91.8 |
| WL1 | Paired reads created from WL1_R1 + WL1_R2 | 420,779,456 | 404,086,394 | 377,826,369 | 93.5 |
| WL2 | Paired reads created from WL2_R1 + WL2_R2 | 439,626,326 | 422,272,136 | 394,438,143 | 93.4 |
| WL3 | Paired reads created from WL3_R1 + WL3_R2 | 370,789,824 | 356,666,082 | 334,585,814 | 93.8 |

**B**

24207 sequences detailed as follows matched to the reference genome ID ASM2204524v1

- •21801 CDS (15547 annotated as genes)
- •1367 non-coding_RNA
- •420 misc_RNA (miscellaneous RNAs = small uncharacterized RNAs)
- •618 tRNA
- •1 uncharacterized RNA

**Percent GC:**

| Sample | GC% before trimming | GC% after trimming |
|---|---|---|
| L1_1 | 39.8 | 39.6 |
| L1_2 | 41.3 | 41.2 |
| L1_3 | 41.1 | 40.9 |
| L1_4 | 39.7 | 39.6 |
| L1_5 | 39.6 | 39.4 |
| L2_1 | 39.7 | 39.6 |
| L2_2 | 40.4 | 40.3 |
| L2_3 | 40.5 | 40.3 |
| L2_4 | 38.5 | 38.3 |
| L2_5 | 38.8 | 38.6 |
| L3_1 | 40.5 | 40.4 |
| L3_2 | 40.9 | 40.7 |
| L3_3 | 41.3 | 41.1 |
| L3_4 | 41.5 | 41.4 |
| L3_5 | 41.2 | 40.9 |
| L3_6 | 40.9 | 40.7 |
| G1 | 36.7 | 36.5 |
| G2 | 37.2 | 38.2 |
| G3 | 34.4 | 37.2 |
| G4 | 34.4 | 34.3 |
| H1 | 40.6 | 40.5 |
| H2 | 41.6 | 41.5 |
| H3 | 41 | 41 |
| H4 | 40.9 | 40.9 |
| WL1 | 40.4 | 40.5 |
| WL2 | 40.7 | 40.7 |
| WL3 | 40 | 40.1 |

**Supplementary Figure 2: principal component analysis (PCA) completed using transcripts with a TPM expression ≥ 5 for all libraries (samples) associated with EXP-1 (larval stages)**. Samples WL1_1 to 3, WL2_1 to 3 and WL3_1 to 6 were collected from the LA07 colony. Samples WL1_4 and 5 and WL2_4 and 5 were collected from heterozygous individuals obtained from the crossing between the LA07 and the SLAM5X colonies. The PCA analysis did not show differences between samples of the same larval stage, collected from different colonies. Abbreviations: L1 = first larval state; L2, second larval stage; L3 = third larval stage; WL = whole larva sample.

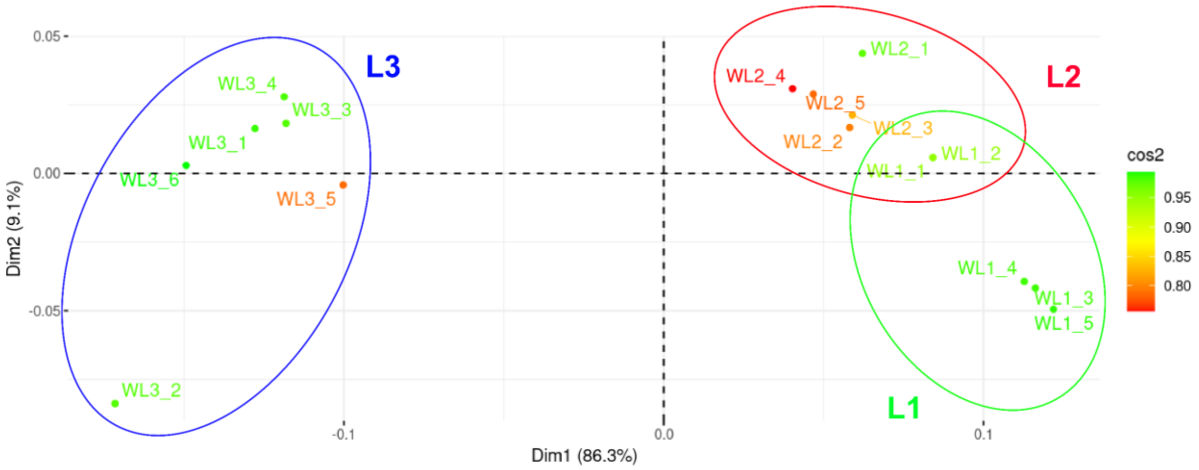

**Supplementary Figure 3: ABC transporters preliminary Neighbor Joining phylogenetic analysis**. Abbreviations Dmel = *Drosophila melanogaster*; Lcup = *Lucilia cuprina*; Lser = *Lucilia sericata*.

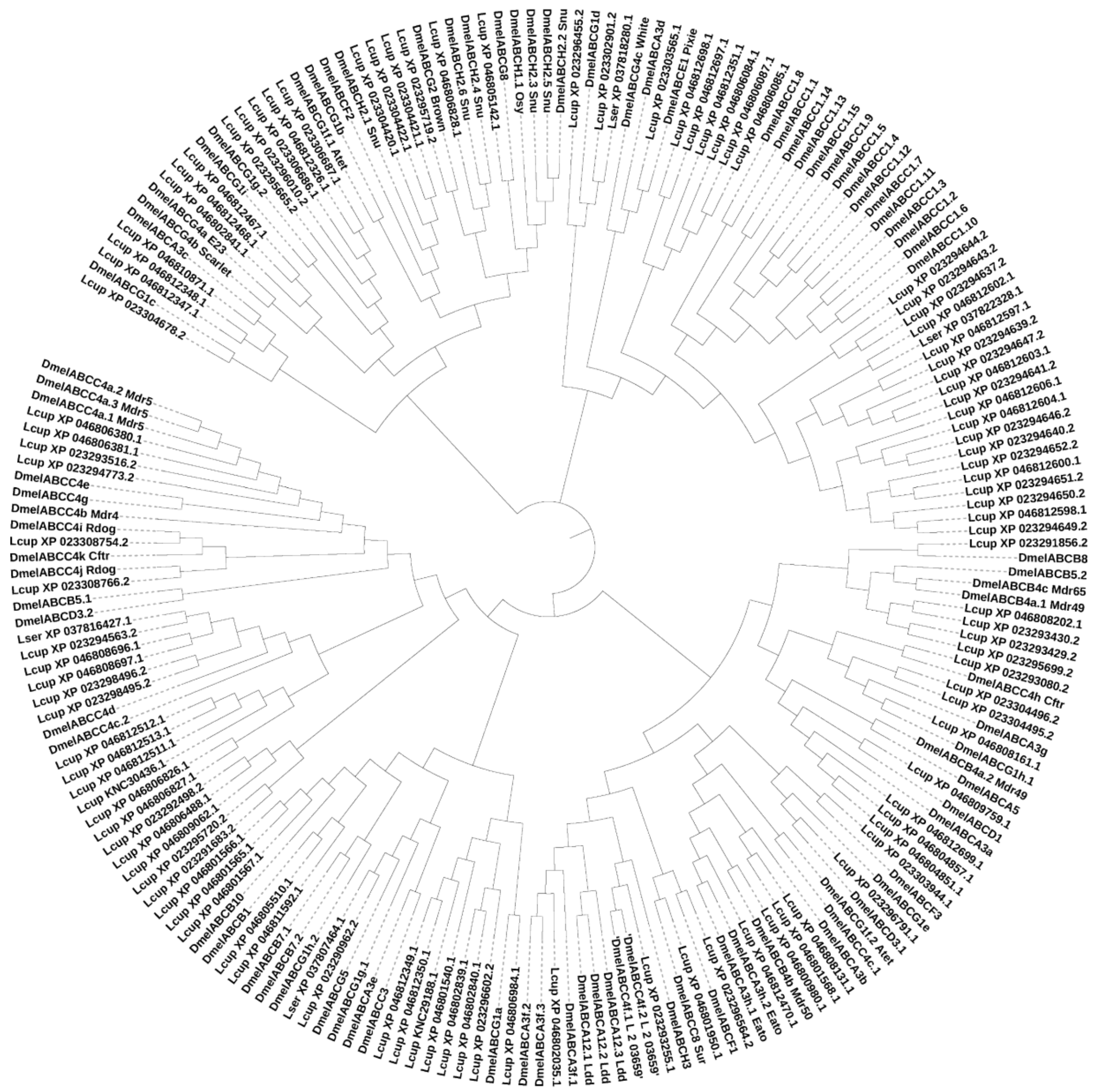

**Supplementary Figure 4: guide RNA (gRNA) cutting efficiency tested *in vitro* using the EnGen Spy Cas9.** Amplicon size without cutting was 439 bp and after Cas9 *in vitro* cutting generated two overlapped amplicons of 224 and 215 bp, respectively. Abbreviations: C = control, uncut genomic DNA; MW = molecular weight; T = treated, gDNA + Cas9.

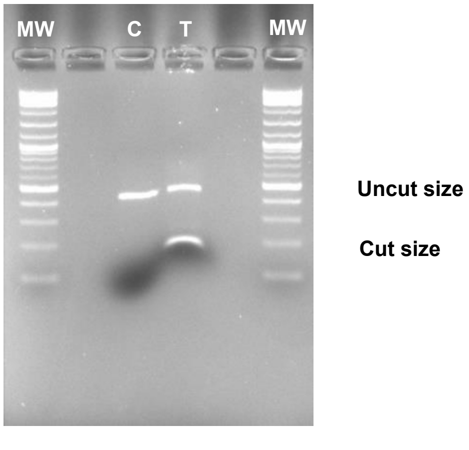

**Supplementary Figure 5: *L. cuprina* eggs, larva stage 3 (L3) and adult flies expressing the ZsGreen marker before the genotyping analysis to determine *LcupOrco* insert landing site.** (**A**) L3 showing transient expression of the marker 7 days after *wt* eggs injection; (**B**) eggs obtained from crossing a *LcupOrco* mosaic male with a *wt* female; (**C**) *wt* L3 on top vs. a heterozygous ZsGreen L3 at the bottom; (**D**) heterozygous ZsGreen L3 on top vs. homozygous for the same marker at the bottom; (**E**) dorsal view of a *wt* adult male (left) vs. G1 heterozygous ZsGreen male (right) under bright field; (**F**) same males showed in **E** under a green filter; ventral view of the same males using bright field (**G**) and a green filter (**H**).

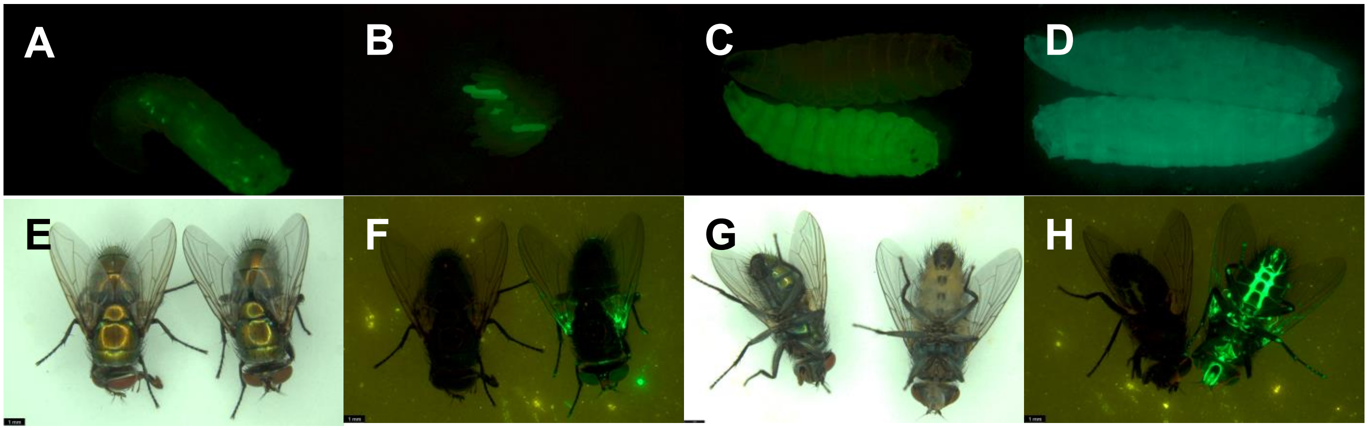

**Supplementary Figure 6: PCR amplification of gDNA of samples collected from *LcupOrco* larval diet preference test and female olfaction assay.** (**A**) samples of larval olfaction assay; (**B**) samples of adult female olfaction assay. Primers, amplicon sequence and cycling parameters are detailed in **Supplementary Note 1** (see Supplementary Materials and Methods section). Abbreviations: MW = molecular weight.

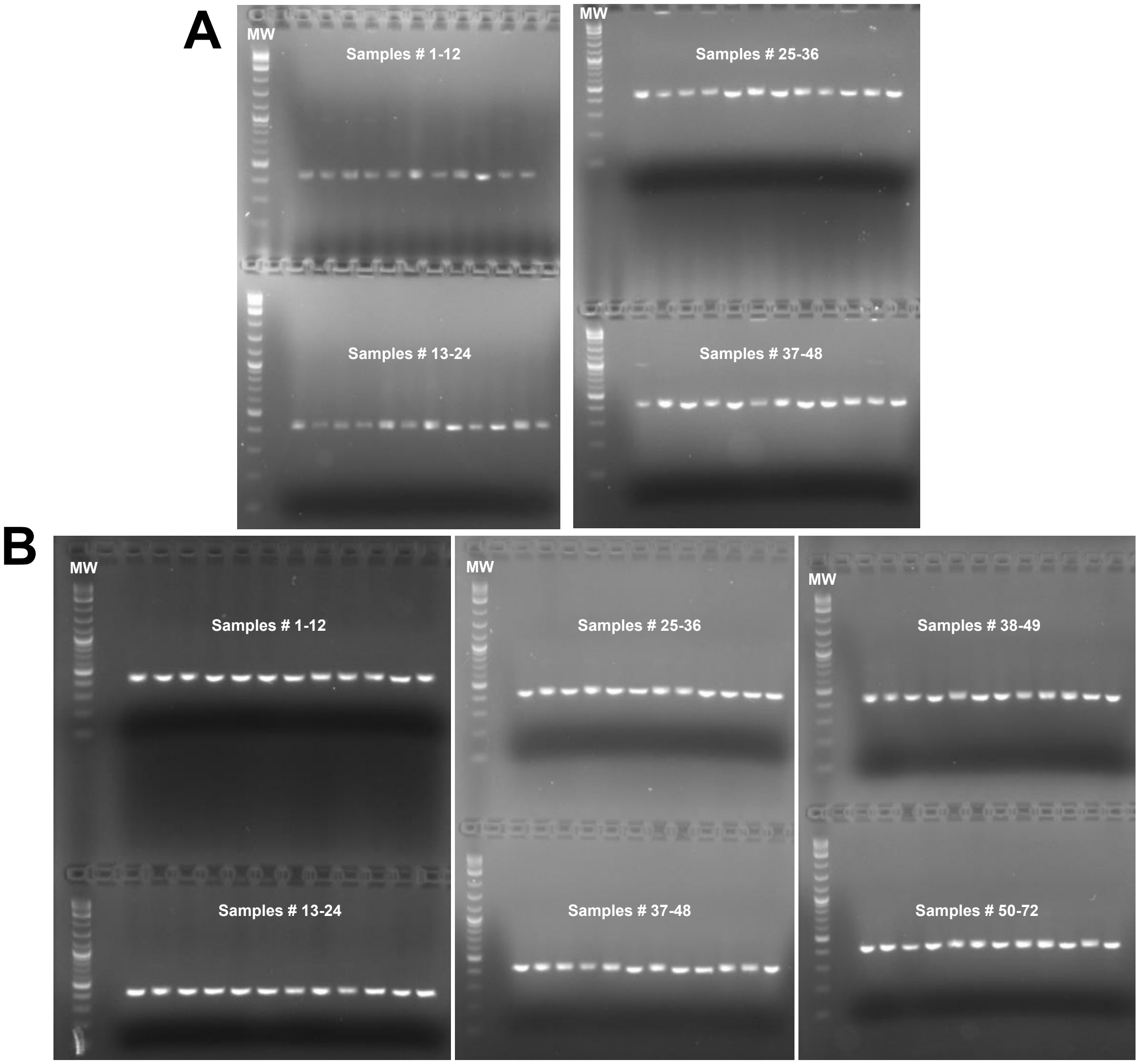

**Supplementary Figure 7: Sanger sequencing results for samples collected from *LcupOrco* larval diet preference test and adult female olfaction assay.** (**A**) genotyped samples from # 1 to 48 of larval diet preference test; and (**B**) genotyped samples from # 1 to 72 of adult female olfaction assay. Synthego software was used to analyze sample's chromatograms and indels % = 0 corresponds to *wt* samples, from 1 to 89% to heterozygous samples, and ≥ 90% to homozygous samples. Results were compiled along with single larva preferences in **Supplementary Table A** and **B** (see below).

# A

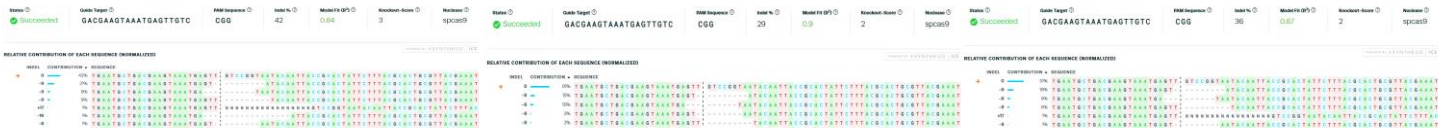

### Sanger Seq. Samples # 4-6

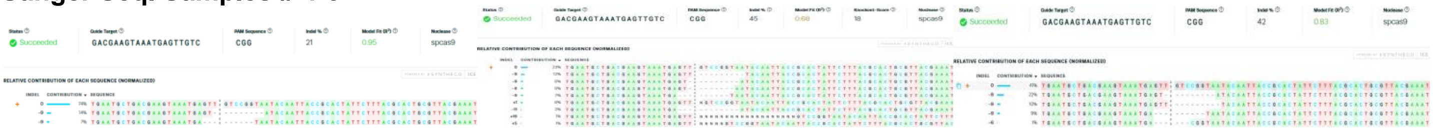

### Sanger Seq. Samples # 7-9

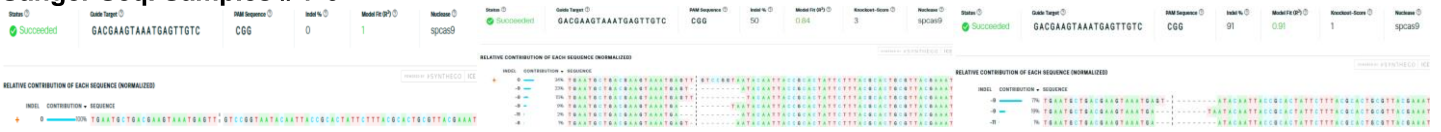

### Sanger Seq. Samples # 10-12

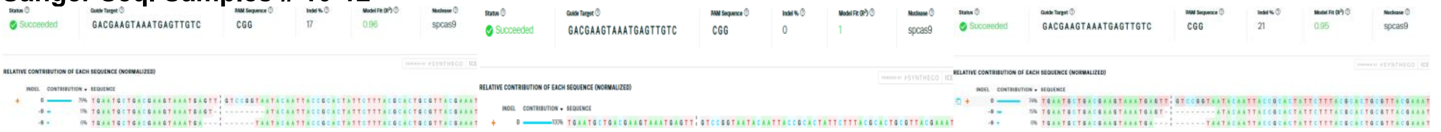

### Sanger Seq. Samples # 13-15

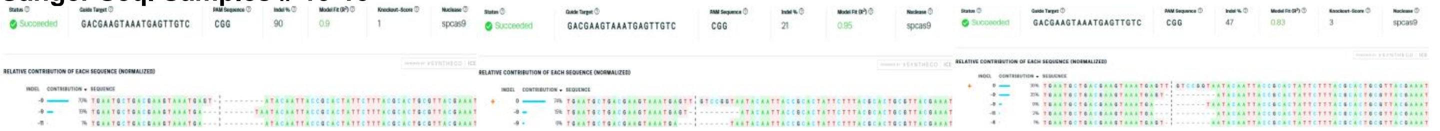

### Sanger Seq. Samples # 16-18

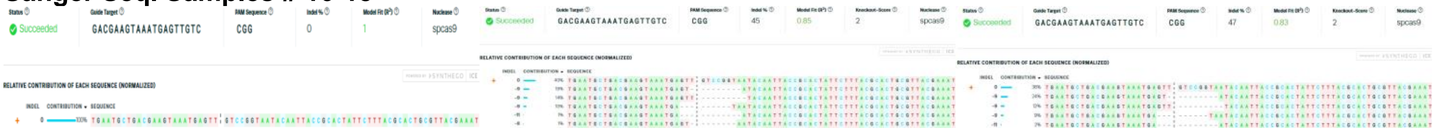

### Sanger Seq. Samples # 19-21

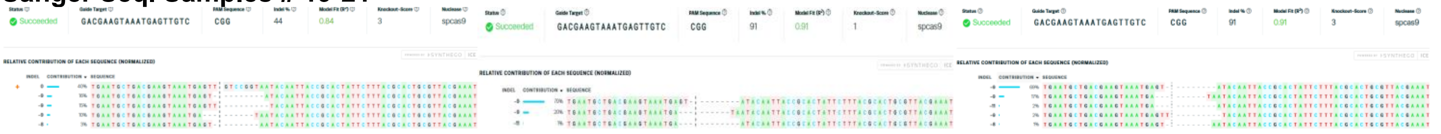

### Sanger Seq. Samples # 22-24

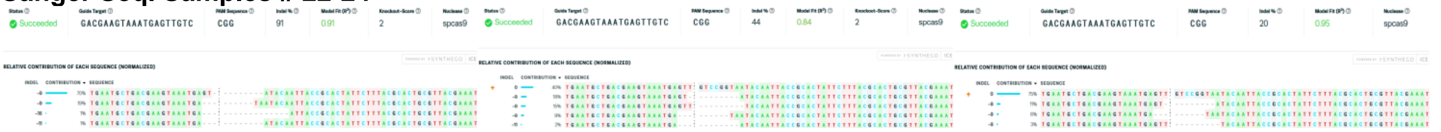

### Sanger Seq. Samples # 25-27

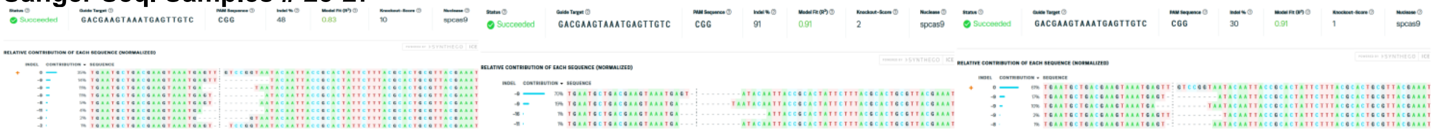

### Sanger Seq. Samples # 28-30

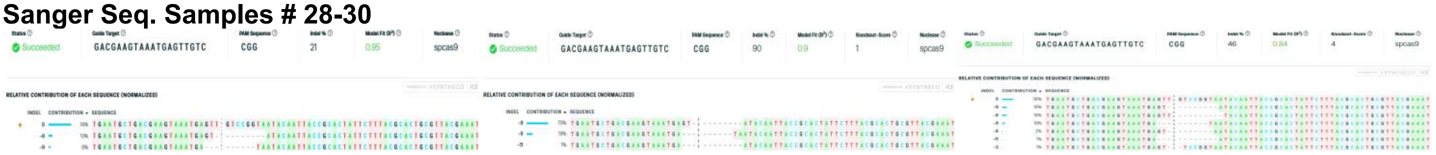

### Sanger Seq. Samples # 31-33

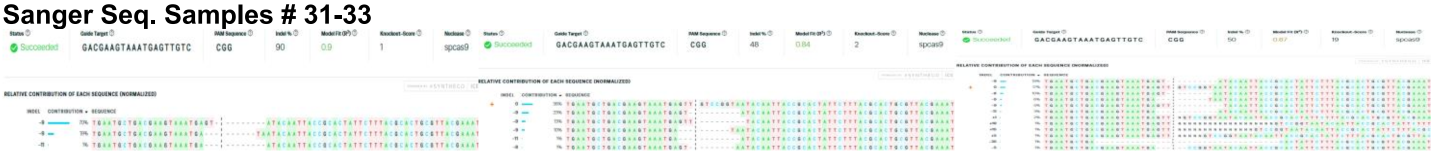

### Sanger Seq. Samples # 34-36

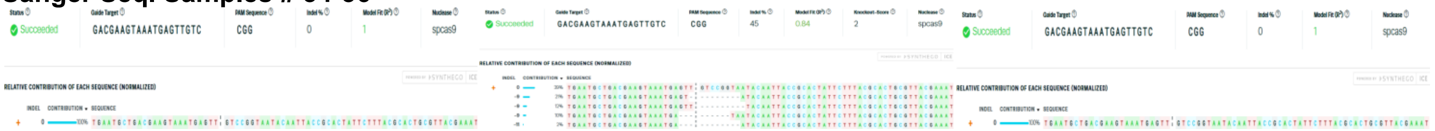

### Sanger Seq. Samples # 37-39

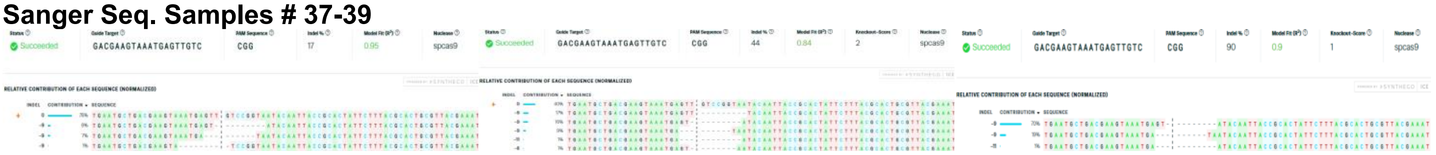

### Sanger Seq. Samples # 40-42

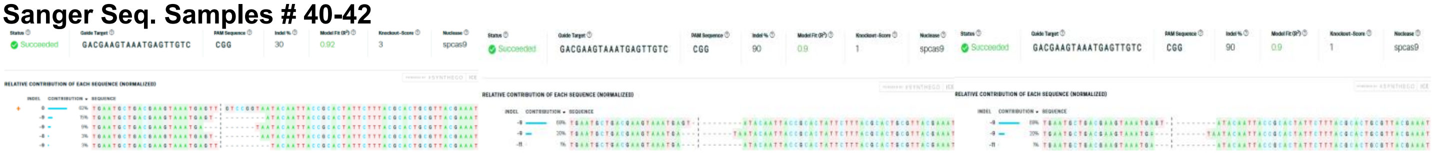

### Sanger Seq. Samples # 43-45

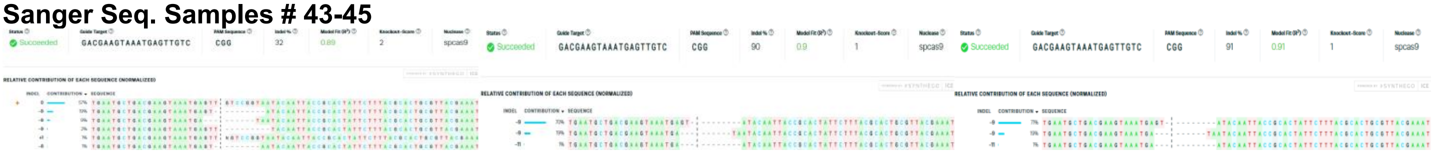

### Sanger Seq. Samples # 46-48

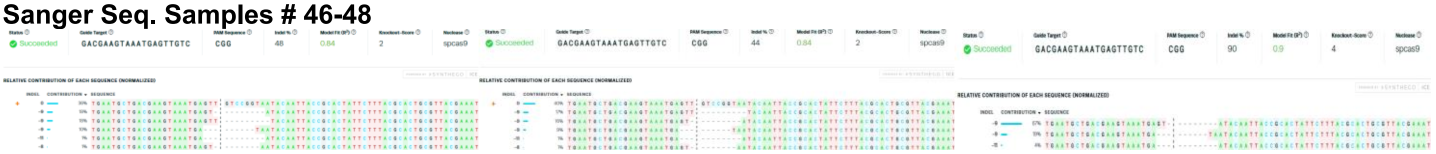

# B

### Sanger Seq. Samples # 1-3

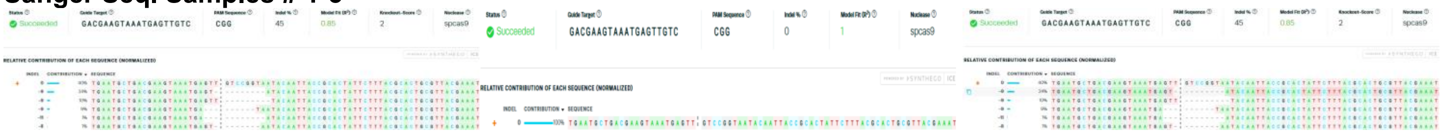

### Sanger Seq. Samples # 4-6

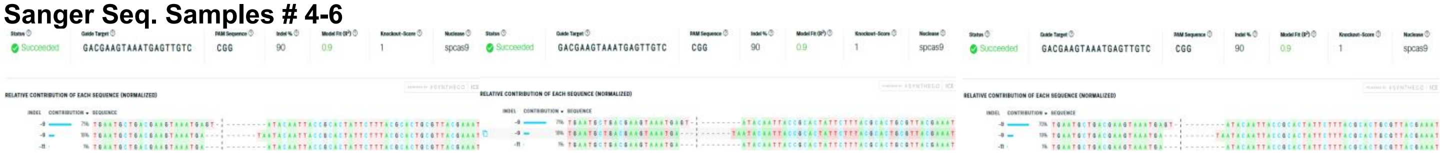

### Sanger Seq. Samples # 7-9

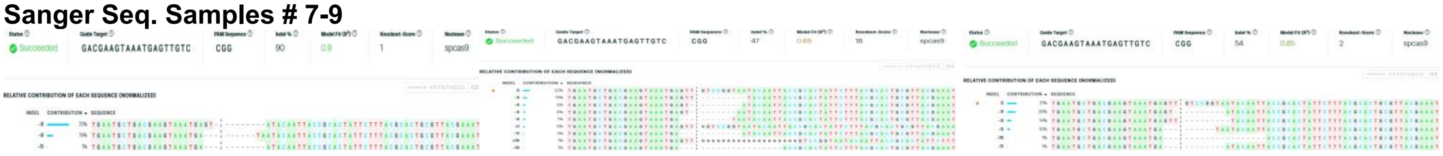

### Sanger Seq. Samples # 10-12

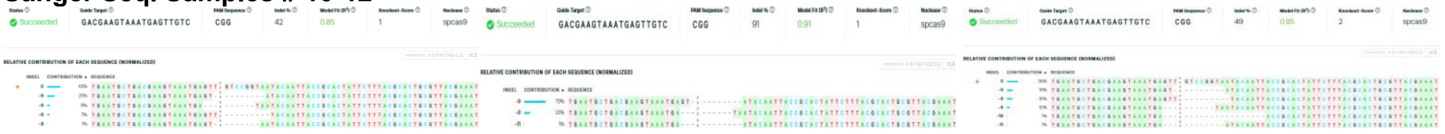

### Sanger Seq. Samples # 13-15

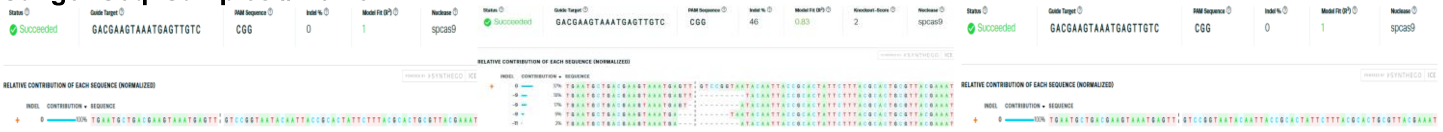

### Sanger Seq. Samples # 16-18

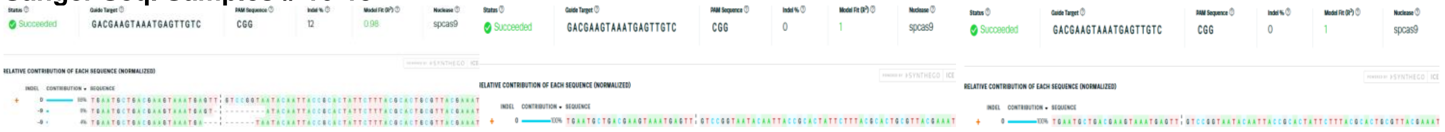

### Sanger Seq. Samples # 19-21

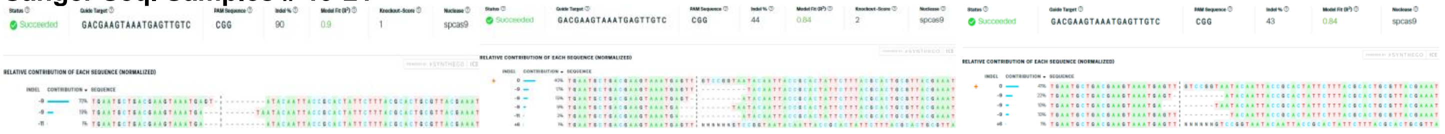

### Sanger Seq. Samples # 22-24

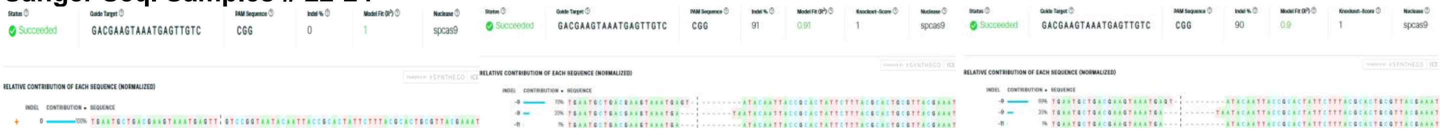

### Sanger Seq. Samples # 25-27

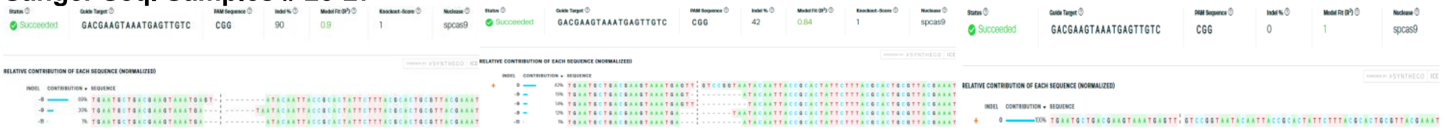

### Sanger Seq. Samples # 28-30

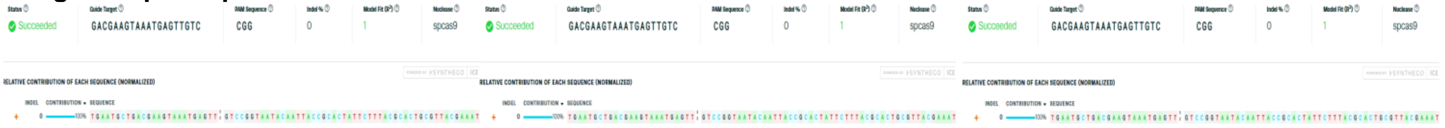

### Sanger Seq. Samples # 31-33

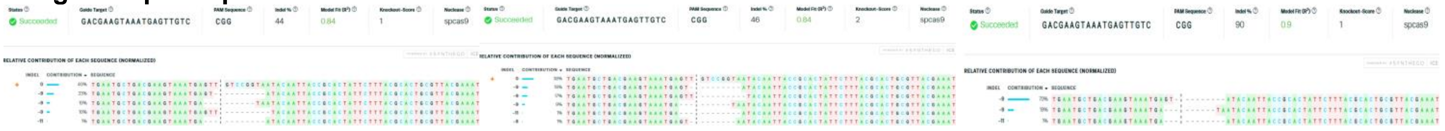

### Sanger Seq. Samples # 34-36

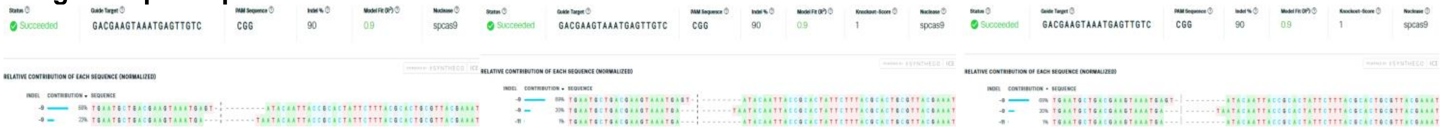

### Sanger Seq. Samples # 37-39

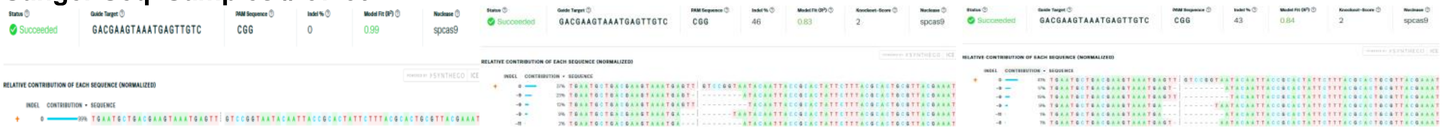

### Sanger Seq. Samples # 40-42

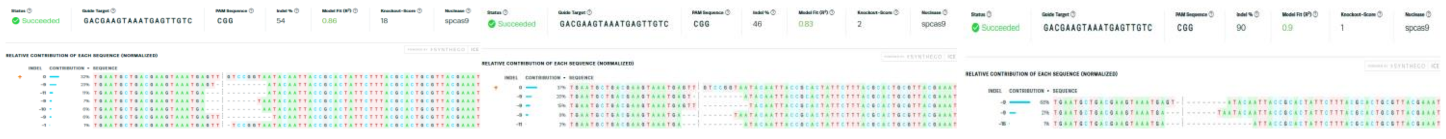

### Sanger Seq. Samples # 43-45

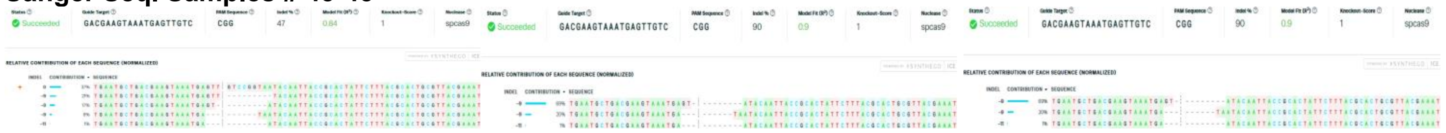

### Sanger Seq. Samples # 46-48

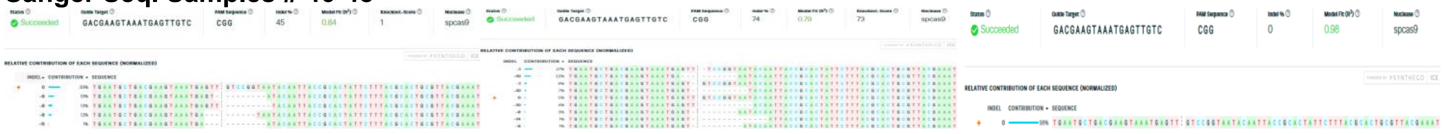

### Sanger Seq. Samples # 49-51

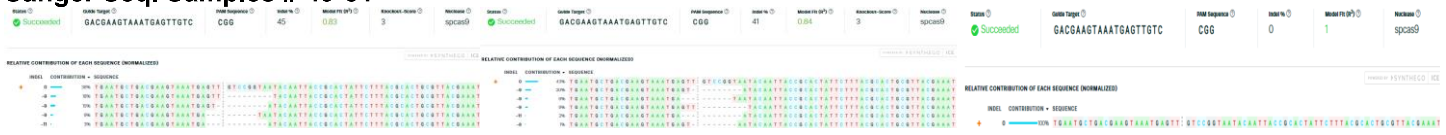

### Sanger Seq. Samples # 52-54

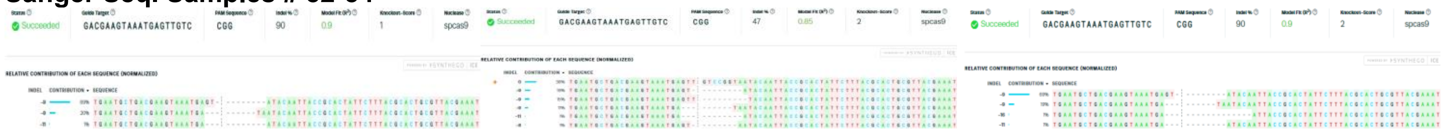

### Sanger Seq. Samples # 55-57

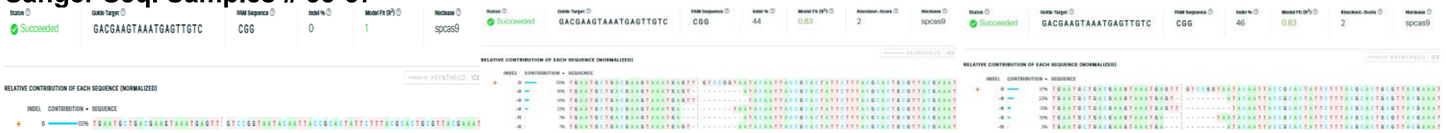

### Sanger Seq. Samples # 58-60

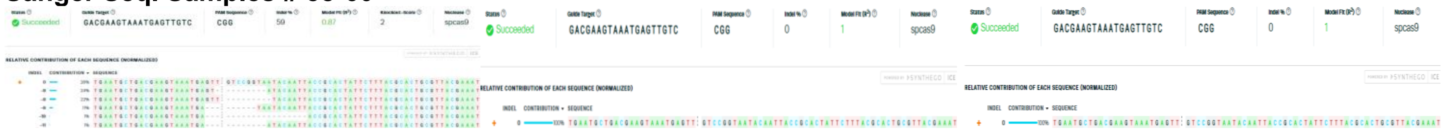

### Sanger Seq. Samples # 61-63

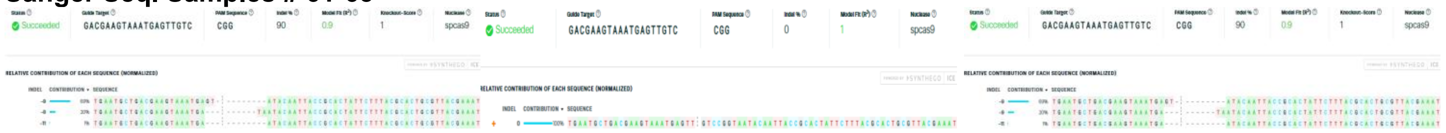

### Sanger Seq. Samples # 64-66

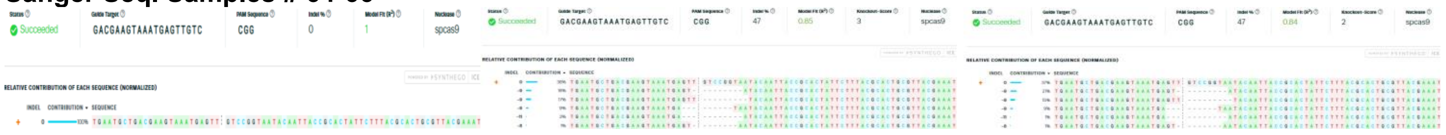

### Sanger Seq. Samples # 67-69

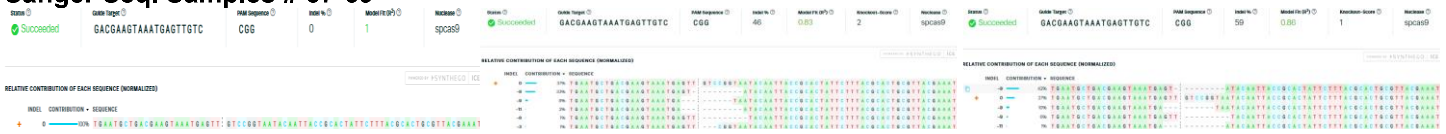

### Sanger Seq. Samples # 70-72

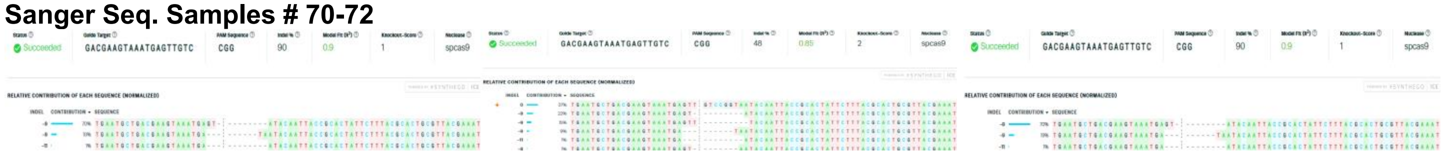

**Supplementary Table 2: compiled data of samples associated with *LcupOrco* behavioral assays.** (**A**) larval diet preference test; (**B**) adult female olfaction assay. The numbers one through eight after the letters in table B, refer to the main chambers of the olfactometer and numbers after dots, to single females from the same chamber. Abbreviations: F = fresh beef; FC = fresh cold beef; FH = fresh hot beef; HET = heterozygous; HOM = homozygous; NC = non-choice; NG = non-genotyped; R = rotten beef; RC = rotten cold beef (25±1 °C); RH = rotten hot beef (33±1°C); WT = wild-type. Hyphen symbol means non-tested by Sanger Sequencing.

# A

| Larva # | Diet chosen | Larval weight | Sanger Seq. Sample # | Confirmed genotype by Sanger Seq. |
|---|---|---|---|---|
| **1** | RC | 11.3 | 1 | HET |
| **2** | RC | 22 | 2 | HET |
| **3** | RC | 16.8 | - | NG |
| **4** | RC | 12.4 | - | NG |
| **5** | RC | 14.8 | 3 | HET |
| **6** | RH | 12.3 | 4 | HET |
| **7** | FC | 8 | 5 | HET |
| **8** | FC | 19.2 | 6 | HET |
| **9** | RC | 39.6 | - | NG |
| **10** | RC | 35.2 | - | NG |
| **11** | FC | 33.1 | - | NG |
| **12** | RC | 22.8 | - | NG |
| **13** | RC | 17.9 | 7 | WT |
| **14** | RC | 24 | - | NG |
| **15** | RC | 15.2 | - | NG |
| **16** | RH | 24.1 | 8 | HET |
| **17** | RC | 18.3 | - | NG |
| **18** | RC | 3.9 | 9 | HOM |
| **19** | RC | 16 | - | NG |
| **20** | RC | 11.9 | 10 | HET |
| **21** | RH | 6.8 | - | NG |
| **22** | RC | 32.6 | - | NG |
| **23** | RC | 17.1 | - | NG |
| **24** | RC | 14.1 | 11 | WT |
| **25** | RC | 29.7 | - | NG |
| **26** | RC | 20.8 | - | NG |
| **27** | RC | 14.7 | 12 | HET |
| **28** | RC | 15.2 | - | NG |
| **29** | RC | 3.8 | - | NG |
| **30** | RC | 19 | - | NG |
| **31** | RC | 9.8 | 13 | HOM |
| **32** | RC | 24.8 | - | NG |
| **33** | RC | 33.9 | 14 | HET |
| **34** | RC | 29.1 | - | NG |
| **35** | RC | 23.2 | - | NG |
| **36** | FC | 13.7 | 15 | HET |
| **37** | RH | 31 | 16 | WT |
| **38** | RC | 11.3 | - | NG |
| **39** | RC | 16.3 | 17 | HET |
| **40** | RC | 23.9 | - | NG |
| **41** | RC | 18.2 | - | NG |
| **42** | RC | 20 | - | NG |
| **43** | RC | 20.7 | 18 | HET |
| **44** | RC | 13.3 | - | NG |
| **45** | RC | 5 | - | NG |
| **46** | RH | 4.1 | - | NG |
| **47** | RH | 3 | 19 | HET |
| **48** | RH | 5.6 | 20 | HOM |
| **49** | RC | 19.5 | - | NG |
| **50** | RC | 18.8 | 21 | HOM |
| **51** | RC | 5.4 | - | NG |
| **52** | RC | 5.8 | 22 | HOM |
| **53** | RC | 21.2 | - | NG |
| **54** | RC | 14.7 | 23 | HET |
| **55** | RC | 20.6 | 24 | HET |
| **56** | RH | 31.4 | - | NG |
| **57** | RC | 10.8 | - | NG |
| **58** | RC | 10.2 | 25 | HET |
| **59** | RC | 21.7 | - | NG |
| **60** | RC | 13.6 | 26 | HOM |

| Larva # | Diet chosen | Larval weight | Sanger Seq. Sample # | Confirmed genotype by Sanger Seq. |
|---|---|---|---|---|
| **61** | FC | 9.3 | - | NG |
| **62** | RC | 14.8 | 27 | HET |
| **63** | RC | 21.5 | 28 | HET |
| **64** | RC | 26.5 | - | NG |
| **65** | RC | 23 | - | NG |
| **66** | RH | 20 | - | NG |
| **67** | RH | 26.6 | 29 | HOM |
| **68** | RH | 15.4 | - | NG |
| **69** | RH | 16.5 | 30 | HET |
| **70** | RC | 16.7 | 31 | HOM |
| **71** | RC | 19.2 | - | NG |
| **72** | RC | 30.9 | 32 | HET |
| **73** | RC | 31.6 | - | NG |
| **74** | RC | 24.3 | - | NG |
| **75** | RC | 16.2 | 33 | HET |
| **76** | FC | 19.2 | - | NG |
| **77** | FC | 16.1 | - | NG |
| **78** | RH | 19 | - | NG |
| **79** | RC | 33.4 | - | NG |
| **80** | RC | 24.6 | 34 | WT |
| **81** | FC | 34.7 | 35 | HET |
| **82** | RH | 2.9 | - | NG |
| **83** | RH | 4.1 | - | NG |
| **84** | RH | 16.6 | - | NG |
| **85** | RC | 16.9 | - | NG |
| **86** | RC | 3.2 | 36 | WT |
| **87** | FC | 32.9 | - | NG |
| **88** | RH | 29.4 | 37 | WT |
| **89** | RH | 30.8 | - | NG |
| **90** | RC | 3.9 | - | NG |
| **91** | FH | 4.2 | 38 | HET |
| **92** | RH | 11.4 | - | NG |
| **93** | RH | 23.1 | - | NG |
| **94** | RC | 27.6 | - | NG |
| **95** | RC | 6.8 | 39 | HOM |
| **96** | RC | 10.6 | - | NG |
| **97** | RC | 22.5 | 40 | HET |
| **98** | RC | 32.2 | - | NG |
| **99** | RC | 3.8 | 41 | HOM |
| **100** | RC | 13.2 | - | NG |
| **101** | NC | 13.1 | 42 | HOM |
| **102** | FC | 25.7 | 43 | HET |
| **103** | RH | 20.9 | 44 | HOM |
| **104** | RC | 22.5 | - | NG |
| **105** | RC | 26.6 | 45 | HOM |
| **106** | RH | 29.5 | - | NG |
| **107** | RH | 6.7 | - | NG |
| **108** | RH | 18.3 | 46 | HET |
| **109** | RC | 9.8 | - | NG |
| **110** | RC | 8 | - | NG |
| **111** | NC | 4.6 | 47 | HET |
| **112** | RH | 9.9 | - | NG |
| **113** | RH | 4.5 | - | NG |
| **114** | RC | 12.2 | 48 | HOM |
| **115** | RC | 35.3 | - | NG |
| **116** | RC | 18.2 | - | NG |
| **117** | RC | 13.4 | - | NG |
| **118** | RC | 8.7 | - | NG |
| **119** | RC | 13.6 | - | NG |
| **120** | RC | 9.3 | - | NG |

**B**

| Female # | Preference | Sanger Seq. Sample # | Confirmed genotype by Sanger Seq. |
|---|---|---|---|
| **1** | R1.1 | 1 | HET |
| **2** | R1.2 | 2 | WT |
| **3** | R1.3 | 3 | HET |
| **4** | R1.4 | NG | - |
| **5** | R1.5 | NG | - |
| **6** | R1.6 | NG | - |
| **7** | NC1.1 | NG | - |
| **8** | NC1.2 | 4 | HOM |
| **9** | NC1.3 | NG | - |
| **10** | NC1.4 | NG | - |
| **11** | NC1.5 | NG | - |
| **12** | NC1.6 | 5 | HOM |
| **13** | NC1.7 | 6 | HOM |
| **14** | NC1.8 | 7 | HOM |
| **15** | NC1.9 | NG | - |
| **16** | NC1.10 | NG | - |
| **17** | NC1.11 | NG | - |
| **18** | NC1.12 | NG | - |
| **19** | NC1.13 | NG | - |
| **20** | NC1.14 | NG | - |
| **21** | NC1.15 | NG | - |
| **22** | F2.1 | 8 | HET |
| **23** | F2.2 | 9 | HET |
| **24** | F2.3 | 10 | HET |
| **25** | F2.4 | 11 | HOM |
| **26** | R2.1 | 12 | HET |
| **27** | R2.2 | 13 | WT |
| **28** | R2.3 | 14 | HET |
| **29** | NC2.1 | 15 | WT |
| **30** | NC2.2 | 16 | HET |
| **31** | NC2.3 | 17 | WT |
| **32** | NC2.4 | NG | - |
| **33** | NC2.5 | NG | - |
| **34** | NC2.6 | NG | - |
| **35** | NC2.7 | NG | - |
| **36** | NC2.8 | NG | - |
| **37** | NC2.9 | NG | - |
| **38** | NC2.10 | NG | - |
| **39** | NC2.11 | NG | - |
| **40** | NC2.12 | NG | - |
| **41** | NC2.13 | NG | - |
| **42** | NC2.14 | NG | - |
| **43** | NC2.15 | NG | - |
| **44** | NC2.16 | NG | - |
| **45** | F3.1 | 18 | WT |
| **46** | F3.2 | 19 | HOM |
| **47** | R3.1 | 20 | HET |
| **48** | R3.2 | 21 | HET |
| **49** | R3.3 | 22 | WT |
| **50** | R3.4 | NG | - |
| **51** | R3.5 | 23 | HOM |
| **52** | R3.6 | NG | - |
| **53** | NC3.1 | 24 | HOM |
| **54** | NC3.2 | 25 | HOM |
| **55** | NC3.3 | 26 | HET |
| **56** | NC3.4 | NG | - |
| **57** | NC3.5 | NG | - |
| **58** | NC3.6 | NG | - |
| **59** | NC3.7 | NG | - |
| **60** | NC3.8 | NG | - |
| **61** | NC3.9 | NG | - |
| **62** | NC3.10 | NG | - |
| **63** | NC3.11 | NG | - |
| **64** | F4.1 | 27 | WT |
| **65** | F4.2 | 28 | WT |
| **66** | F4.3 | 29 | WT |
| **67** | F4.4 | NG | - |
| **68** | R4.1 | 30 | WT |
| **69** | R4.2 | 31 | HET |
| **70** | R4.3 | 32 | HET |
| **71** | R4.4 | NG | - |
| **72** | NC4.1 | 33 | HOM |
| **73** | NC4.2 | 34 | HOM |
| **74** | NC4.3 | 35 | HOM |
| **75** | NC4.4 | NG | - |
| **76** | NC4.5 | NG | - |
| **77** | NC4.6 | 36 | HOM |
| **78** | NC4.7 | NG | - |
| **79** | NC4.8 | NG | - |
| **80** | NC4.9 | NG | - |
| **81** | NC4.10 | NG | - |
| **82** | NC4.11 | NG | - |
| **83** | NC4.12 | NG | - |
| **84** | NC4.13 | NG | - |
| **85** | NC4.14 | NG | - |
| **86** | NC4.15 | NG | - |
| **87** | F5.1 | 37 | WT |
| **88** | F5.2 | 38 | HET |
| **89** | F5.3 | 39 | HET |
| **90** | F5.4 | 40 | HET |
| **91** | F5.5 | NG | - |
| **92** | R5.1 | 41 | HET |
| **93** | NC5.1 | 42 | HOM |
| **94** | NC5.2 | NG | - |
| **95** | NC5.3 | NG | - |
| **96** | NC5.4 | NG | - |
| **97** | NC5.5 | NG | - |
| **98** | NC5.6 | NG | - |
| **99** | NC5.7 | 43 | HET |
| **100** | NC5.8 | 44 | HOM |
| **101** | NC5.9 | 45 | HOM |
| **102** | NC5.10 | NG | - |
| **103** | NC5.11 | NG | - |
| **104** | NC5.12 | NG | - |
| **105** | NC5.13 | NG | - |
| **106** | NC5.14 | NG | - |
| **107** | NC5.15 | NG | - |
| **108** | NC5.16 | NG | - |
| **109** | F6.1 | 46 | HET |
| **110** | F6.2 | 47 | HET |
| **111** | F6.3 | 48 | WT |
| **112** | F6.4 | NG | - |
| **113** | R6.1 | 49 | HET |
| **114** | R6.2 | 50 | HET |
| **115** | R6.3 | 51 | WT |
| **116** | R6.4 | NG | - |
| **117** | R6.5 | NG | - |
| **118** | NC6.1 | 52 | HOM |
| **119** | NC6.2 | 53 | HET |
| **120** | NC6.3 | NG | - |
| **121** | NC6.4 | NG | - |
| **122** | NC6.5 | 54 | HOM |
| **123** | NC6.6 | NG | - |
| **124** | NC6.7 | NG | - |
| **125** | NC6.8 | NG | - |
| **126** | NC6.9 | NG | - |
| **127** | NC6.10 | NG | - |
| **128** | NC6.11 | NG | - |
| **129** | NC6.12 | NG | - |
| **130** | F7.1 | 55 | WT |
| **131** | F7.2 | 56 | HET |
| **132** | F7.3 | 57 | HET |
| **133** | R7.1 | 58 | HET |
| **134** | R7.2 | NG | - |
| **135** | R7.3 | 59 | WT |
| **136** | R7.4 | 60 | WT |
| **137** | R7.5 | NG | - |
| **138** | R7.6 | NG | - |
| **139** | R7.7 | NG | - |
| **140** | R7.8 | NG | - |
| **141** | R7.9 | NG | - |
| **142** | R7.10 | NG | - |
| **143** | NC7.1 | 61 | HOM |
| **144** | NC7.2 | 62 | WT |
| **145** | NC7.3 | 63 | HOM |
| **146** | NC7.4 | NG | - |
| **147** | NC7.5 | NG | - |
| **148** | NC7.6 | NG | - |
| **149** | NC7.7 | NG | - |
| **150** | NC7.8 | NG | - |
| **151** | NC7.9 | NG | - |
| **152** | NC7.10 | NG | - |
| **153** | F8.1 | 64 | WT |
| **154** | F8.2 | 65 | HET |
| **155** | F8.3 | 66 | HET |
| **156** | F8.4 | NG | - |
| **157** | F8.5 | NG | - |
| **158** | F8.6 | NG | - |
| **159** | F8.7 | NG | - |
| **160** | R8.1 | 67 | WT |
| **161** | R8.2 | 68 | HET |
| **162** | R8.3 | 69 | HET |
| **163** | R8.4 | NG | - |
| **164** | NC8.1 | 70 | HOM |
| **165** | NC8.2 | 71 | HET |
| **166** | NC8.3 | 72 | HOM |
| **167** | NC8.4 | NG | - |
| **168** | NC8.5 | NG | - |
| **169** | NC8.6 | NG | - |
| **170** | NC8.7 | NG | - |
| **171** | NC8.8 | NG | - |
| **172** | NC8.9 | NG | - |
| **173** | NC8.10 | NG | - |
| **174** | NC8.11 | NG | - |

**Supplementary Figure 8: eggs obtained from *LcupOrco* mutated females.** (**A**) eggs obtained from 8-day-old *LcupOrco*$^{+/-}$ females 3 days after crossing them to *NPF*$^{-/-}$ males expressing the ZsGreen marker. The same marker was removed from *LcupOrco* females after confirming that the *LcupOrco* insert landing site was not located withing the *LcupOrco* gene locus in the *L. cuprina* genome. The insert was removed by selecting non-fluorescent larvae and crossing males obtained from them for two generations vs. *wt* females. In addition, males from each generation were genotyped using protocols described in **Supplementary Note 1** (see below) to confirm the presence of indels and point mutations within the *LcupOrco* coding region. The fluorescence in eggs confirmed the mating between the *LcupOrco*$^{+/-}$ females and *LcupNPF*$^{-/-}$ males and egg fertilization. *LcupOrco*$^{-/-}$females did not lay eggs at this time; (**B**) part of *LcupOrco*$^{-/-}$ females laid eggs 10 days after mixing them with *LcupNPF*$^{-/-}$ males but eggs were not fertilized as showed under green filter (**C**), and no larvae emerged from them; part of the *LcupOrco*$^{-/-}$ females laid fertilized eggs 17 days after mixing with *LcupNPF*$^{-/-}$ males, as showed under bright field (**D** and **F**) and green filter (**E** and **G**); these eggs produced healthy progeny.

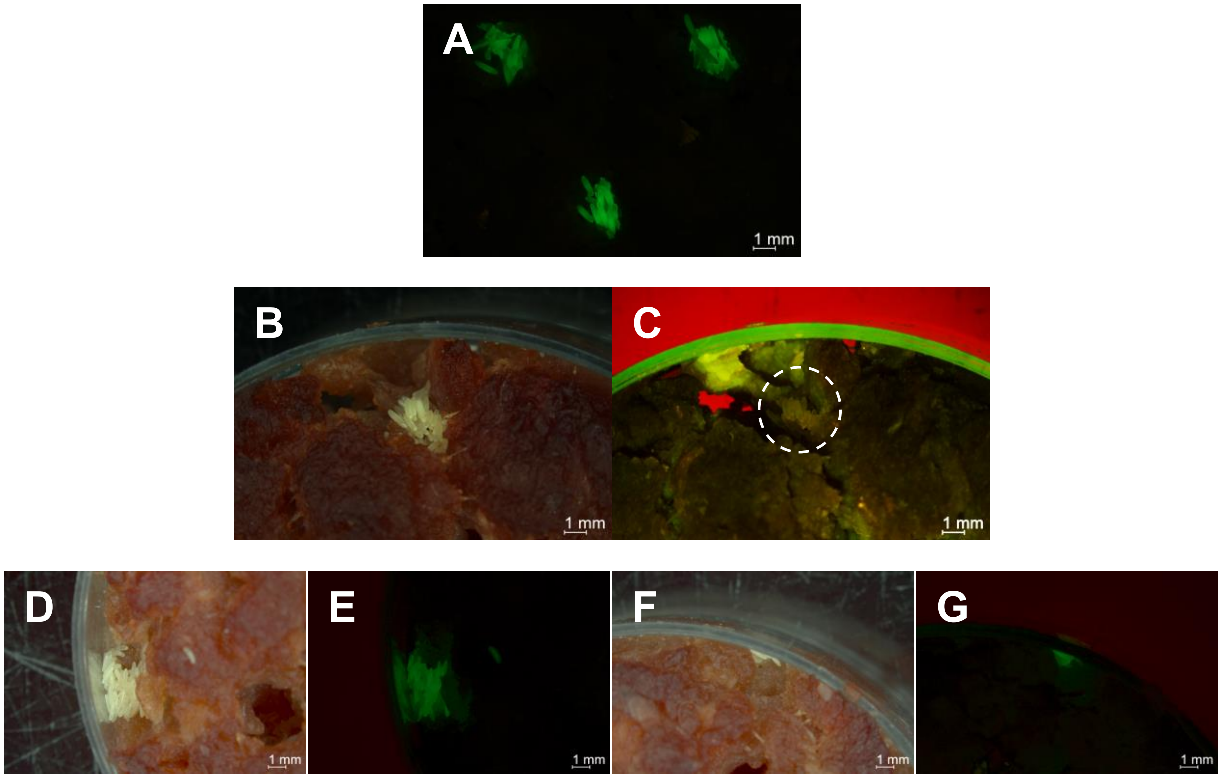

**Supplementary Figure 9: ovaries from 10-day-old *L. cuprina wt* and *LcupOrco* mutated females.** Three females per condition were dissected (columns). All females were provided with tap water and sugar, but the presence of protein in the diet and mating condition changed between groups; *wt* virgin females not fed (**A**-**C**) and fed with protein (**D**-**F**); *wt* mated females not fed (**G**-**I**) and fed with protein (**J**-**L**); *LcupOrco*$^{-/-}$ mutated females fed with protein (**M**-**O**).

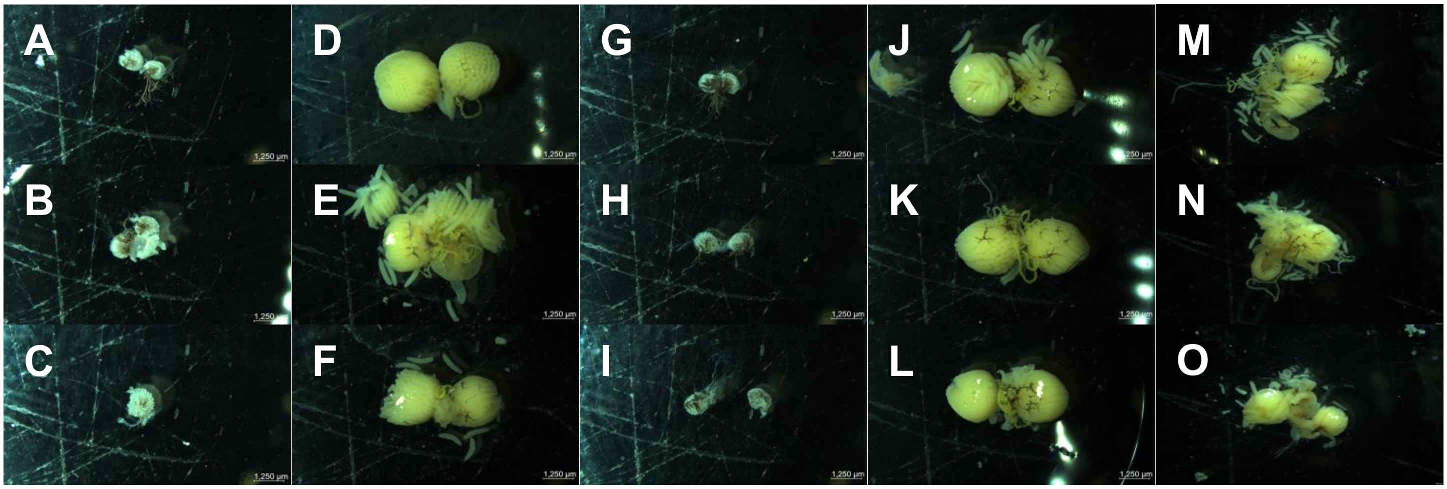

### Supplementary Materials and Methods

**Supplementary Figure 1**: **larval stages and tissues used for the RNA-Seq experiment.** (**A**) whole larva stage 1 (L1); (**B**) whole larva stage 2 (L2); (**C**) lateral view of a whole late larva stage 3 (L3) and (**D**) ventral view; (**E**) whole early larva stage 3 (WL); (**F**) first segment designated as "head" (H) from an early L3, detailed within a dashed white circle; and (**G**) gut from an early L3, where crop have been separated from the cardia originally connected by the foregut (not present in the picture).

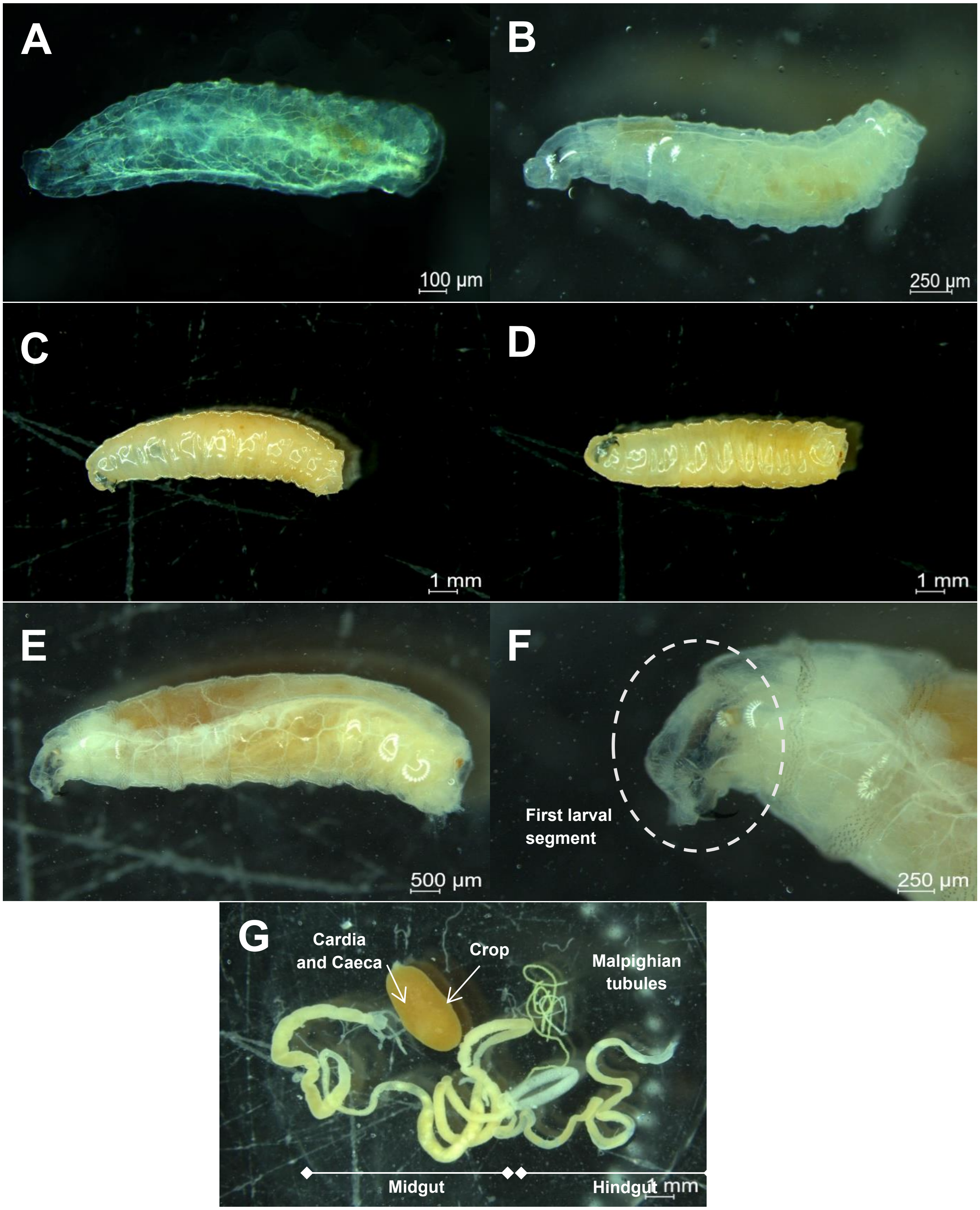

**Supplementary Note 1: *LcupOrco* gene, guide RNA (gRNA) sequence, and primers used to determine potential indels and point mutations at the gRNA cutting site (genotyping).**

**(A) Orco gRNA (Lcup-Orco-crRNA1) without the PAM sequence at the 5'-end.** The ortholog sequence of this gRNA was previously used to edit the first exon of the *LcupOrco* gene in a species closely related to the fly under study, the new world screwworm *C. hominivorax* [1]. Its location, including the PAM sequence underlined, was highlighted in bold in the *LcupOrco* gene sequence provided below.

Lcup-Orco-crRNA1 sequence: GACGAAGTAAATGAGTTGTC (+)

**(B) Primers used for genotyping and to test the gRNA cutting efficiency *in vitro* using the EnGen Spy Cas9.** Primers positions were highlighted in bold in the *LcupOrco* gene sequence provided in **D** (see below).

Lcup-Orco-sgRNA_1_Fw sequence: *ATGCAGTCGAATCTACAACC*
Lcup-Orco-sgRNA_1_Rv: *gccaacaataatcatggtttca*

**(C) PCR cycling parameters**

Initial denaturalization = 3 min at 98°C.

Denaturalization = 30 sec at 98°C.
Annealing = 30 sec at 60°C. Repeat x 40 cycles
Extension = 30 sec at 72°C.

Final extension = 5 min at 72°C.
Hold = ∞ at 10°C.

The Q5 High-Fidelity Taq Polymerase 2X Mix [New England Biolabs (NEB), Ipswich, MA, USA, Cat. # M0491S] was used for DNA amplification. Reaction was performed in 20 µl of final volume and 300 nM for final primer concentration.

**(D) *L. cuprina Orco* gene sequence including 5'- and 3'-UTR regions, detailing exons and introns as UPPERCASE and lowercase letters, respectively, primers positions and gRNA highlighted in bold and PAM sequence underlined.**

```
cttaatatttaaagtttatttttaattttttacaaaagtcgaaaatttaaaaggtatgcaaagtagggagaaaactgaaaaagttaatacttaaagtatgtacaaattataa
gtacattatacatggaaaattatgttaaagtgtatttaaaccttaatcattacaggcgaaaccgaagtgtattttccaaaaaggattatattgattaaagcaatatttaaaa
aatatttgtttgaattcagaaatatctatttttatactctacaccactataattaggagtgtattatgtgttggttctgacatttgtaaatattggtccttatatatatcag
tcagctcagcatcactttctgagttgattgtggtatgtctgtctatgaatttcgacacatactagtttctttatccaaggacatttgacttatgtaaaatctggctttattt
ggatcatttggaccgattgtggtagggtcaattagggaccggttcacagtggtatattaaaaagagggaaataattcggtaacttctaaacggtaaatccgattttaatgaa
atttggtatgcacaaagagaaggtgttgtcgagcttaggttttgaatttggaccttaaaggccaaccagggacccgatggggggtcctcaaattaggacacctcggctatgt
taaattttaaaatgatcctatttcttcttttgagttccgatttaaaaaaaaaattcgtttatagaatctcctcatcaaacactataaaaaatgtcgtagcagtaaattatc
ttttatagttgaggatatattctcatttgaaaatcaaaattttaaaattttaccgtccttactttagtttttgataatagcgggtcagaaaaattcccgattttcgccat
ttttattcgcttaacaacaagtttatatatcatgcagtgaaaaaaattatgtaaaaatcatgactgagtccaaagttataggcattttaatttaataaattaaaaaaggcga
tttttgccattttttgggaaaaagtatctttttcttttttaagttatctgaaaagtttctaataagatgtatatagaatttatacttttttgaaagctgactttacataaat
acgttaaatgaaacaaaacctaaaactttttgttaccgagtggaccaggtccatccaaaaaaaaaccctatttttttaatgaaaattcaattttgagcaaaaattctcaaatcg
catagtcgatatctaattatagtgacctttttttatatgacccaatatgttcttaatcatttgtaacgggtttcgataaccccgccactggtatggatgtaataggcaaaaa
accaaaaaatccaatttttgggatttttttaaactttttttgggaattccgggatttctaataagaaaattcgaaatttttatttaattttatattttgagtaaaaaaatctac
caaactgtaaaatttttattaaaaaaatattcataaataaaattttttattgctatttgaaaaatttgtatcttcgcaataactgaataaaaaacattttttaatttttttttaaa
aagtattccgcgatttttttaatttaaaaattatatacagttttgaaaagtagacaaaattatataacatacctacctaatgtttttaagtgacaaattaattagggaaaa
ctcaacatcttttagaaaatttacctagtgctcgatgaggagattctatatacgaaatttttttaaatcggaactcaaataaagaaataggatcgttttaaaaatttaacat
agccgaggtgtcctaatttgagaacctgtactcgttcctgtactcgtatttctccacagtgataaaagtttaagtcacactctgaagtagatcttatagaagggattgtgcc
aattatagaccgatcctaataaaatttctttaatagatttaagttcatataagacttgtttatggagaatttagctatgactgttgtaattgttttcccaagttccactta
tattatccatatgcgtaaaattttaactctatatatctttaatcattcgaaccctatcgtgctttaaacagacagacggacatggctagatcatcttacaatttaatgagga
cccagaatctttttttgatggtagttataaaaaatctgctgtaggtccgattgcgtcctcaagacaagtctgaggaagatataatttcaccaaaacgaagtctattgctacaa
atttttctgtccgtccgtccgtccgtccgtccgtccgtctggctgtccatgtaaaccttgtgcgcaaggtaccgcaattttcaagataattggatgaaatttggcacaaaca
cctttttaggcccaaggacgaagcctcttgaaaatggttaatatcggtccattatttcgactatccccatacaaccgtacccccaaatagggcctttgggcttataatta
atttaaatgatctattatgtcaacaaaagtcgagaaaactaagttttatagaactttaaatgacactaccgattttttgtaatgattgggctacatttgaccctagcccccat
acaaactccccttcagaaaatgacttaaaggtcaaaattcacttataaacactaatgacacttttaaattctacataaataatattgaagtagacttaactcccccctaccaa
catttataaggatagggccatatttgccctactcccctttgagccctcttgtaaaaaatttatttttggcaaataaaaataaataaaaatattccgaaataaagttaaaaa
caaatcaaatgctttatttttttaaataatccccattttaacatacatactgactggtgtagggtatcatatggttggctacatccgactatacattcatacttgttttaaa
ttataaacatagttaagacttattgaagacttcgcttaatataagtctaaactatgttttaagtatttttgtagaaataagtactagaacaattaacataaatttagcctga
aaagccatatttgttaggtatggttgtatggaggccatatattttcttttttttttcattacaaacataagcgcaaacccaatataacttcccttagtaaagtggtgtagg
ctaaacttaggtcaacttgaatcttacagattcaagtttggaagatttcctaaaaaaagaaatacttttgctaaatttttgaaaatataatttgacttaattctttaatgtt
tatattaaacctttttctaaatatatcatatttcagataataagagagatatgtctgaagtttaaataaaatcaaataatgttaacccataaactgtgaagctgaaaatcaag
ttatgcctattttttatgaaccttcactattatttaatttaaaccgatttaaaaaaaaaaaacttgagtaaagtatataataccgcaggaataatttcagaaattgttcta
aaaacaactagttttagaatgaatgcaaatagaatcactttagattctttaacaacaacctttttcactaatgtgtcgtcattatttgtaaaaccctttcttcggaacattgt
tatggttcttaaagagttttaaggaactagttctttgatactaattatcagggctaggatttctatgcaaatgcatattttttagtcagtaactcaaaataacttttaactag
ttttcttttcaaatcaaagaattttctacaaaatggcatcgatttgttgttgcatattttgttataaatacataaattgcatattttgctttaaattgcatatattttgcat
atttttaacattttttattgcatattttgcatgtttttttaaaattttctaaaacaaattgttttgtttttttttctgtttttatattttaaaacaaattaggtatttat
gtacatatttagatttttttaaataactttcaacaaatttcaatttaaaaaaaaaatactaggttctatgaaatacaatttacaatcacgtccaaagtattggtccttaaag
```

ttcagttcaatttctgagtctgtccatatgagtgtatatataaaaaggtaactagatgaaattttgttcttagatctctctcacgttcataattcacttaaaaacattacca
aacgataaaattcggaagaaatgtaatttatacaaaaatatgaaaattcaccaaattttattagaattgccatcaagctttccgcttacagaaattagatatatgtatacta
ttttaagttgtttcttagtctttaggttagtggtcagtgttctagtctggtagaccggagttggtgggttcgattcccacccgtggcactggttaaggagcgcacaacaggc
ctaggtgtatttcttcgaatttgatgtgtatacttacatcctcctttcaaacttactaactaactaacttagttaactacctaactaacttactagtctttatatatgtcat
tctcgggcttgtggttccgttaaagtggtattatagtacataaaacagtccttaataacctgtaaaaacttaagacctgattatatattaatatgtactttcgcagagaaat
atttttggtacatttttttcacattttagcgcatatttaccattttatactgtataaaacataaattttaaatttttagtgcatgaaaatcctttgcttgctaaagagt
tttaagaaactagttctttgatactaattgtacattgattctatactagtttaatggaatgcgtttctcaaaattgattagcgaatacattctccaaagatttcctagaact
agttccggacttgacaaattcgaactaaaatgattgatttaagaactagttccagaaccgttccaaagaatgagtgctgaatccatagcgcttcctcagaatcagttacgat
gctgacaatttagaacaacaataattcatctcaatgtcaaagtagaaagagtacaggctttaatcacaaaaactgacaaaatgggcggaattactatagtgggcgcggcatc
ttccatattaagaaaatattaaatctgtttatctcagaaaatgtaacagagccatcacaagtttcaatgtgattagttaggataaaaaatgggcagactagcagcagtgggc
gtggcaactcccagaaactaatagaactagattactaaaattttgcacaattgactttcatattcaactcaacattttgggaaaataaccggacgttgatcaggtggtggct
tggaacccccatatgaattacatgggttgattggcccctcccatatgaattgtatacaatacaattttatttaatttttttcatagcttacattaaaatcctaattagggttc
gataaaattagcgaacttgcaataatttgagtggtatctcccatgagtaagtgggttttttttgcctttaggatagcaaagcatactgggtatatctagtattataattatat
aaaatttccgtaatatatgtaccaaatattttattctatattagttcttaaagtgcttaccaaaatttatttcgttgacctgtgttattccgataccgaatcatttgctcgt
gaaataatataaaattatatataaaaatgacagcttgttgctttgatcgaaattattggatcaatcgtccattaagataaatattttgtattgaaaaaaatattaaatgtat
ttttcttaattgatccctttgaaattttaacaattgcattatctcactttaccttatctccatgcag***ATGCAGTCGAATCTACAACC***AACGAAATATGTTGGCCTTGTGGCC
GACTTAATGCCCAACATTAAGCTGATGAAATATTCTGGTTTATTTATGCATGCGTTTACGGGTGGATCACCGCTGTTGAAAAAAGTCTATTCGTCCATACATTTAGTGTTAA
TATTGGCACAATTCATGTTTATACTTGTCAATATGGCTCTGAATGCT**GACGAAGTAAATGAGTTGTCCGG**TAATACAATTACCGCACTATTCTTTACGCACTGCGTTACGAA
ATTTATTTATCTGGCTGTGAATCAGAAGAATTTTTACAGgtaagtgatgggggtcaccactaaacggcggagtatggatgtgaggattgtttgtatgtgtgcgtgtgtatgt
gagtgtttgtttaatgggcgactatgggattgaaaa***tgaaaccatgattattgttggc***ttttaattgctacttgtgcaaaatgcgaacaaaatacaaggaataaattaaatc
tggcctctgacatggttgacaatattaaatgggtctttttttattgtttgcaaactgtgtctgtgctagtgactaggctgagctgtgtgccacagtaaaaatgtcacaagaa
tcaattaatcgagtaatttattttgcattaaaagttattctttgttctttctttttcaaccattaactttgtggtgtatctgcagAACTTTGAACATTTGGAATCAGGTTAA
TACTCATCCATTGTTTGCTGAATCCGATGCTCGCTATCATTCAATCGCTTTAGCAAAAATGAGAAAATTATTCTTCTTGGTAATGCTGACGACAGTAGCATCTGCAGTGggt
aagttaaatgcttgggaggagtttatgcctgaacgtgtaatactgtgattaaagtggtaattaattaaatttatctttgtactttaGCTTGGATAACCATCACTTTCTTTGG
TGAAAGTGTAAAGTTTGCCTTCGACAAGGAGACCAACTCCTCTATAACAGTGGAAATACCCAGATTGCCTATTAAATCATTCTACCCCTGGGATGCTAGCCAGGGTATATTC
TATACGATATCATTTGCCTTTCAAGGCTACTACTTACTATTCTCCATGGTACACTCGAACTTGTGTGATGTGCTCTTCTGTTCGTGGTTAATATTCGCCTGCGAACAACTGC
AACATCTAAAGGGCATTATGAAACCTTTAATGGAATTATCTGCCTCCTTAGATACATATCGTCCCAATTCAGCAGCACTCTTTCGTTCACTTTCTGCCAACTCCAAGTCAGA
GTTAATCATTAATGAAggtaagccaaaagacagctgccagccagagtgttttcagttgatgatgtatgtatgtgcataaatcaattttattttaatttcgttttataGAAAA
AGAACCTCCCAGTGACCTGGATATGACCGGCATCTATAGTACCAAGGCCGATTGGGGTGCACAGTTTCGTGCGCCCACAACACTACAAACATTCAATGGTGTCAACGGTGGC
AATCCGAATGGTTTGACAAAGAAACAGGAAATGATGGTACGCAGCGCTATCAAATATTGGGTGGAGAGGCATAAACATGTTGTCAGgtaaataagttagcgacccaactaac
tccatccattctaactaagtagcataaaaattcgtttattgaacccaagtttgtgtcagagttctttatatttgttcaatcgtaacgaaaaggaaaacataaacaaaacgtc
agtgatggaaggaaggaaacagttgttgtttaaggaaatttaaattttaattacaaacgaactgtgtaatgtcctttaagcatagaaaacactcagagaaatgtgttagtta
atgaacagaaactagtatatatatcggcttcgggttcttaatattgatctcctcctgagacatccgtgtaaagaatattctgcttccggtaaaggatcgaacacgtctgtcc
aatatgttaaatatatattatcaacgtgtccttaaaccggatttgatcaaaataatgttttgataaagataagaacatgcagactgtggtttcgacattatatcttacttaa
ttttcggtcagaggaatctttcaaatagtaccacatacagccacgtgcccagtttcttaaattggaattggtccaactataattttaagtcaatatttcataattgaatta
agactctattttatctgttatattaaacagatcggataacttgttagtgttatattttgatatttccatttagtttctgattaaaccagtggtgctcaaacacatactatc
atagatagttttatgtgtgttggtaaggcagcagagaacagaattgcattcaaaataaaagtatactaataaaatgactccaaactataaagttaatgctaactgttaaata
tttattatggattttgacagcgagcgtgtatcaaagtctatatataagaaactatcaactgttctctcatgttctactacattaattcggctttcgttgacaaatgttgatc
agcaactgatttgattttgtttgagagcaaaccaactgatattataaatagaggaagaggggccaggtgaaataatagaccgatcttaaccatttttaataggctttgtccc
aggatattgaatagaatattgtgtaccaaatttcattgaaatatctccaaaatttcgacctgtagtttgattaaaaggtttgcatggacggacaggcggacggacatagcta
aatcgactcagaaaatgattctggtatactttaaggtggatatatacgaccaatattattgtgcgttataaattgttatagctagaatcttcaaaaactttgggtagcaaa
gcacacagggtatagctagtatacatatatattcagggaatattttataaatatattgtagtacaattccttgaattagagccccttgcagatcatcttaattaaataata
atgtttcacacataacaaaattcaatttctaatttttttgaaacaatttttactgtcgattttctgtcagtgtaactaatgctgaagtattaaagacaataaacaaggccct
ttcgtttttcagtctattgcaatcgaataaagtttttttataagtaacttagtttgtttgttaaactcatcactttttatttactatttatagacaaatggtttgtttagg
ttttcaacagttatagatccttctattgacgcgtccgtccagctgtccgttttctatttgtctattccttcttttaattttgttgcatgtttatttgtggcaaaaaaaaaag
taaaaccaatggaaaattcacagataaaaaataaagccaataagcattttctatatctaacatattcatagtcctgactaaatccatcatcgtccatcatgtttgcatcgat
gatttggcttttattatcctatgtttgctggagtgtgtctgtacatctgactgtttggtttccgtttccatgtttgtttaatgtctgtttatgtatatgtatagttgggtat
taattgaaataaaagtaaatattttaattccttaaagtaacattgctgaatatggtatttagtcttattgattttctctgatttctccattaatttatttatatatttttta
tttctttgagtttttattaacacattattttatctaaactgttgaactaagaggattatttttcatatttattagacaaattaaatgaaattaatatttaatgaactagact
agtggtaaataatgatttttttaagaaatctattaatgtgcttggtttatagtacagacaaaagcaaatgttaatccagggactattgtacacttgactaaaggacttacca
gcgctgaatctggcctctaagccatcgtgtcaggaagtttcaaaaaacaccttttttaatcaattttaagacgctataacagtggtcagttatgaatttttatgccataatta
cgttaatatccctctacctttccaaaactttgcatttttttaatgataagaaatgtcttaaacgtccaaaattagtatcgtaatgaaattctgcataagaatttgtaataca
aaatacatgtatatcctttccctaaatttaacaagtaagagtgatatattcgaatataccgaatcaaaccgtttgcagtaaatgtaaaacgttaggttgatattattatgag
attttggcttgtacaaaacatacttctttgtatcgaacttcatagctatactagcatttttttagccagtaatgttcgtttaagccaatttctaaggggggccttatatgggg
gcagggtcaaccatggaccgattctcattaaattcggtagacagatttgtaattataaggaacttacttgtgtcgaacttcaccgctatactcgtatttgaagcaagaagt
gagcgtttaagctattctctgaggagaaacttatacgatgagttggataattatgaacggatcctcataaatttcggtagattgattaagctatgtccgtctgtgtgtttgt
tgaaatcagtttttagagtaccccagatatcggcagatacgaatcttcaataattctgttagacatgctttcgacaagatcgcgatttaaaatcagcaaaatcggtccataa
ataccggatatatgagcaaaaagccgagacaacctctaaaaatttcatcaaaaattgagattttttttcaggctcagacagaaattgcaaaaaacaaaatacataatcatac
ggaaaattttattattgtactcttgtttataaaaacaaggcagagcagcactagtgtgttgttattggctagaaacatgaaaatatgtatgtgaagcctagcttaagttaga
agccataaaagggaactttttagtactttttcgttttcaaaaggtactttttgaatttccgatactattgtacctacaaatatgaaattaagtatgtagatgaaaacccata
aaagggatcttttgggtactttttcgcttttcaaaaggtactcttttgaattttctataatattgaatctagaaacatgaatttgagcatgttaagcccaatgaaagtgatg
gaaaagattactttatggtactgacagtttttgtaacacatcaaggtgaaggttatataagattcggtacagccgaatatagaaccctcacttgtttagtctctagcatttt
ttttttcgaaatatttacttcaacaaaatttgtgttatctatttatcttctccacaaaatgcttcattgatttttttttaaaaacaaggacttgactttttttcaactaaag
ttcacaaataaattctcaaacacatttcttaaatttaaagaatacaaagctcgatgaaatttcatcaccatttcaaataacattatgattataatttgtaaaatttataaat
taaaacaatccccccctagtgtacaagaaagggtattaaagatacgttaaggaaataaagcactttactttcatatttttaggttcattattatagaaaattcatcaaatgaa
aacttgaggaacttgatattgaaaattgtgtataatacattcttaggaactgttgcgaaatataaatgtgtttgtttatttacttcgttttttttttttttttttttttgtc
aagtctaaaacacaattactatgtagtagaaaaatgtcaaactaaaagcaaatattgatttatgtaaaaggaatcgaaatgaaatatttcaatatttgtcaaatttatttat
agttttacacacatacctatgtatatttaaattaataaacaaacatttgcacatacacaattgtaaaccttaataaaattgcttttaacaaggactaaacaaactttctata
aatattgtgtaaatgtagatgcttttaacagctacaactacaagtacatgtatttatatttagtacaacaaaaaagaaataatttttaggggaaaaatttaacatgattttt
gtttttcacaaacctacatgccaccctcttccgaccactaataacattgcgttgttgtttttttctcgtgcatttacagATTAGTTGCTGCCATCGGTGACACGTATGGCGCC
GCATTACTTTTACACATGTTGACATCAACCATTAAGTTGACATTGTTGGCATATCAGGCCACCAAAATAACGGGTGTCAATGTTTACGCTTTTACCGTTATTGGTTATTTGG
GTTATGCATTGGCTCAAGTATTTCACTTTTGTATTTTTGGTAATCGTTTAATTGAAGAGgtaagaggtcgattttaaatactattaaatacacacacatctaaatgtactta
aatttgaatacagtatcttgtcggttggttgaagtacattaaatactagttgtattactgctaagcaatatttgtcatagataactacaaagagatgtacattgggtagatt
atgtgtactttgtgctaaacggaagaaattaatttaaacaaacgactaataataaccaataattatgaaataattttaaaatatgtggcaaagctgtagtgtaaattccaga
caaaatcagattatattttagactatatttattccactagtctatagtctagtctatagtctagtctatagtctagtctatagtctagtctagtctatagtcaagtctatag
tctagtctgtagtctagtctatagtctagtctatagtctagtctatagtctagtctatagtctagtctatagtctagtctatagtctagtctatagtctagtctatagtcta
gtctatagtctagtctactgtctagtctatagtctagtctatagtataatatatagtatagtctaaagtctagtctataaactagtctatagtcttgtctatagtgttgtct
aagtttgtctataaaccagcctgttatctaatctgtagtctagtctatacgtaatctacaatcagggctacgaacttgtccataatctagtctatagtctgtattatagtg
cactctgtggtttagaatatagtccatgtactttatggattgtactctataacttaggatctcatttctactatattcgggcctaaaatctggattatatttagtctagtct

```
gttctatgacaaagttccatgtttagcttacgtctttgacctttacttaacattcaaaatttatgaaaataatataaatttccaaaacatcatattgtgcatgactttaaag
tgtaaatatttattatattatttagttttgcgttttgtagagcaataaggaaatgaaaaagtagaaaaaaactagactcatgcaagcacaaaaacatttttgaagaacttccg
tttgaaatataacaaaaatagattttcgccttttcgccacttttcacttttatgaaaaagtttttgtataaagaaatacacacaggcaacaatgaccatgaaaataaaa
gcacagttaaagaagcagcaaaatatcagcagcacattcactcatcggaaattcgaaagccggaaatgcactaaagtgcttttgttgttttgcgatggtttgaaaatgtata
agtgtgtaagtcttgcacgtatttgtgaatgtgtagaaatttatctctatgtgttagatatttatagaagtgtttatgtgcggctttgtagtctatttatgaacatatatgg
tgtaagtgtatatttgttagcaaatgttttattgttaaagtaaaagttttaaaactttatttttttacattgctttccaatttaatatttcactatttcttatttttcttta
ttttttgtttttcttgccagAGTTCATCCGTTATGGAGGCTGCATACTCGTGTCACTGGTACGATGGTTCTGAGGAGGCAAAAACCTTTGTACAAATTGTTTGTCAACAATG
CCAGAAAGCAATGTCCATTTCGGGTGCAAAATTCTTTACAGTCTCATTGGATTTATTTGCTTCGgtatgttgtttattgtattttaaattaatttttctgtgttatttta
caaagagacattacaggatatgtgaacaaatatgtacatagagaataacatacaacaaataaacctacacatatcctgacgtttacacaaaggcccacatatcacagacaat
tgattcatacctgtgtgttgtaggtcacacaggaacatttaaaaaaaaggaaacagtgacttgcaaagcaaatatttgccattaactaatattgttatcacaacgtatatct
tacatatacatatgtatatacatatgtatatagatacttcgatttatatttttttgaataatattttcaatttttttcaactcagcatggatgatactgtctggtgtgtttgtg
acagttacatttattgtcgcacacaaatgctaaacgtacaaaaggccagacactctagggatcgtatttcaaagggcttttattcgaattttttaacaaaaagtacaaacagg
ctgtaaagtgaatgtcctcaaattgtttttataacattaaaagtgctgttaaaataaaaatggtcatcgttttgttttattcctatagtgaattcaaatttgattgtatagg
ctaagagacctttatacacccagttatgtaacccaagaagcaaatatagctgcattcgtgatcgaaaagatttcttagcacgaatatgtatacgttctttgaacagacaaaa
cgcgacggagggacaatcatacctaacagcagctcgtcacccgcaggttttcaccaaatgcacagcacctccacaccattaaccgccattgccactcagctctcacaagatt
aaattagcaagggccctgtaaaagaacggtcttagtgactcttgacctatcgaaggccttcgactgtctcgccacactcttcgaagatatcctgcaatccacaatgcataac
aagtcgaagaggtggatcgtgaattatataagtggttgtcagtcgtttgtagagttgagagacagacgcttcaaactccgtagagttataccagtcgtactaactaagcact
aacggcacttttgtgcaatttttacctctcgaagctcctcgcacctccacaaggcgtggacgtgatttcatatgcggtcgactgtacgattatggctacggaccacaatatt
aatgaattatgcattgtggtgaatggctatctccatgaaatacatcagttcttcactgcgtcactgggagtgacatttaatagtctttttacttcctcagctcacgtcactg
aaatcacgcacaaattgcgaaaccatgttagctatatataaaacgatcgttcggtctgtggttaattatgctgttccagagtggtcaccttcgctcagtgatacccagtg
gagaaatattcaaagctgtaaaaatgccaccctaaggactgcatacaaattaccttggaacatcatctgtatgaggaaacggaggccctcccagtcaaggaacacaatgtca
tgctgattaaacagttccttctaggatgccatcggaggagtcatcccaacattaacatcatagatggagggacggagggatgctcctccaaggcatgtcgagggaaatatat
gaagatatgcgcaggatcgattggattagactcgaaaacagcagctttgaacgacattcatggagacgccataaatttcgctctcctaggatgccgtatgaatgttgtactt
ggaggtcgcccaccgcctatagcagacacggaaggaaacctgccgcggaaaacaaaagtagttctggcacaactacgatcgggatggagctacaggctcagtgcctactggt
cacgtatagaccgggccgttcaaaacgaatgtactgcagttagagtccccatgacacccacgacatattaaactgcccatccaatcctacttcgctccttccaattgtcaat
catgcagctccaatgtggttcctaacgttgatacccaatgaaccattcgcatagctgccaaaatgcatccctcaagaccgttacggattgattacaggtgtctttagaacac
caactttatgagaaaacgaaaatcctaccagtcaaggaacacaatattacaattttctttagggttgacttcagtgaaatcaccccaaccccaacagttgtagtttcttcg
agttatatacatgatccacatcgttcattgcagtactgaacgatattcatagaggtacggtaaattgcagattatcgtattattatcgtacttggggaagaacacacacaca
cctattggtctcgtatagacagcacgactctcaaacacatgcccagaatgtgttgaagttccacaggctgaaagcatacaatttacggacaacgtaaagaggcagagatagc
ctcgccacggatcacataatcatgtaccgcactttgatctggcaaaaccatatagctcctgccgtatcggtatatatcggtatatatatatatttatatatattatcttccc
ctattccgaaatactcagtacggattccaacaaaataaagccaccttcatttacgcggaaatgatactgatattctttaaggaactaagtgaatttgtttaccttatttaaa
agtgatgtttattgacttccaaagaagcgaaaagaatcaatccatttattttttaattgaaggtatatttttttttgtactagaattttttgaaattaaacaaattttaatatta
tttgagtcaaattagttatattctataatggtacaatttcacttcctaataactgcaaaaaaagcaacataaaaattacttcctgagaatgacttcagatgtagtgaatttg
gaatcaaatataaaggacatcccttctatgtcaagcctgtgttaaaatcatcactttttttaggtctttttcagtacttccatggcgtcaattctacttctttttttgcttct
ttggaagttcttttttttgctgggtattgtcaattagaaagaccttcactttgtgcaacggcatacaaaagcttgtgcgttttcttggtatcggaataagcttctgtttttcc
agaaatctagaaattttataccctacaccactatagtggggagggtattatacgtttgtgctgatgtttgtaacatacaaaaatattggtccaatacccaccttaaagtat
accgatcgtttcagaatcatttttttgagtcgattaagacatgtccgtccgtccgtctgtccggctggctgtccatgtaaaccttgtgcgcaaggtacaggccgcaattttc
aagataatttgacgaaatttggaccaagcatgtttttttgacacagggacgaagcctattgaaaatggttgaaatcggtccattatttcacctagcccccatacaaccgtacc
tcccgatttgaacttttttatgccataattacgtcaaaaattcagctatcactttaaaaattggcacaaataagttttatataagtataaatgacactgcagattttcgtaag
gatcggcctatatttgaccctagcccccatacgaacccccccttcaaaaaatgacttaaacgtctaaaattgacttgtaaccatttgtatcgcaatgaaactcaacaaaacta
actgttatttagaaatatatcctttcccaaattaaccaaggatcggtccatatttgacctatataaagcctcatttagaaattttagtttttttatcaataaattgcttaaa
tattttgaatttattggtaatattcaacataaaactttctttataaaaagtaaaacttttaaaaatatactcatggtgtagggtattatatggtcggccatgcccgactata
cttttctacttgttttctcttgttatgagggcaatttttagagccctcttagaaatatagccaaggcctgaagaatcataaaacttaaaaatttggtagtacttttaataaa
tattgactaaatttcatcttcttacaacccaatgtgcttccactaaatgttaaaaaaatcagtttttccaaaacaactttttcttagaaatatttgcgaaaacaaaaactttta
aaatatttcattcttgtaacaacattaacttggtaaatactatttgttatcaaaacatatacacacatgtacataactagtacatctacatacacaaatacacatcgacaaa
tacataagataaaatactaagtacttggaaatttgttatttacgagtatttcttataatattttcagGTTCTTGGTGCAGTTGTCACATACTTCATGGTGTTGGTGCAGC
TCAAGTAAaattatcaaatatacttacttaatgcaaaaacggcagcaactgcagcggcagctgcaaaaacatttgctgttgtatattttgcattggcaacattttaagataa
gaaacagcaacaccagctacagcaacaataacactctgctaacaatgaaatataaaacaaggacattagcaacaacaacaacaaaaatatatttttaaacttattttataaga
caacaaataattcacctacatttagaaactttatacacacagatacatagtacttcaatacctatacttatatgtctgtgtgcatgtaaaagtattagtgtatctatgagc
tttctaataccaccacttatcaacttaatactaccattttatacagggcaaaccacattttatacatttaagagttgtatggaaagttggttgtttgtttggttagttgcc
cttaaaggcaacaatagatgaagtaaatcagagaatttgatatattgctctcactttaaatttcatgtactggaatggctcaacaactgcatcaacagacgtttattttttgt
aaaaacatttattgtctaagggcgttggttgactgctgccatcatccacaggaatcatttagttgtcagttcttttctgctatttttttgtttctcaaaaatatcttatattc
ttttagattttatcatttgctcgaacattattatttgttgttgtttttgttagaaaatatttctttagtttttaagtaatgttagagattttacattgaaaatgtaaacgaa
tatgtttcgtttaagtttttgaaaattactaaaaaattttgagaaaaatcgaatatttattattattttaaggtaaattcgcactcaatagaatttacgaaatttagaaaaa
ttacaaacattatttacaacttaataaagttcttattatttatgtttaagtatttaatacaaaattttatttaaaaaaattataaaaatttgttggaaataaataat
```

**Supplementary Figure 10: *LcupOrco* insert construction.** (**A**) *LcupOrco* gene fragment including two sequences corresponding to the left (LHA) and right (RHA) homology arms of 1000 bp in length each one, necessary for the CRISPR homology directed repair (HDR) protocol cloned using plasmid pUCIDT-AMP GoldenGate (ordered from IDT). In addition, a DNA section including the restriction cutting sites (RCS) for enzymes XhoI (NEB) and NotI (NEB) was added in between of both sequences in the same plasmid; (**B**) donor plasmid including the ZsGreen Marker followed by the *Hsp83* promoter located between positions 1891 to 7441 [2]; (**C**) ZsGreen Marker obtained from the donor plasmid. The plasmid was linearized using restriction enzymes BpmI (NEB), XhoI and NotI at 37°C for 60 min and the reaction was finished at 65°C for 20 min. After restriction digestion the mix was electrophoresed in a 1.5% agarose gel at 60 volts for 120 min. The band with the size corresponding to the marker (~5 Kb) was recovered from the gel and purified using the Zymoclean Gel DNA Recovery Kit (Zymo) following the manufacturer's specifications; (**D**) to obtain the final construct, 50 ng of linearized pUCIDT-AMP GoldenGate plasmid + 175.5 ng of the marker fragment (1:3 ratio) were ligated overnight (O/N) at 16°C using the T4 DNA Ligase (NEB). After ligation, 1 µl of the reaction mix was used to transform 10-beta competent *Escherichia coli* cells (NEB) following the manufacturer's specifications. Seventy-five µl of undiluted transformed 10-beta cells were poured into an Ampicillin 20 mg/ml (+Amp) Lysogeny Broth (LB) plate and incubated O/N at 37°C; (**E**) Ten clones were recovered from the LB plate and grow in 3 mL of liquid LB +Amp O/N at 37°C and 250 rpm. To obtain the plasmids from clones the ZR Plasmid Miniprep Kit (Zymo) was used following the manufacturer's specifications and evaluated with the BpmI, XhoI and NotI restriction enzymes following the same protocols described above for restriction digestion and electrophoresis; (**F**) Three positive plasmids for the selected restriction cutting sites were sequenced using

Oxford Nanopore and aligned to the *in-silico* construct to confirm the sequence identity. Abbreviations: Kb = kilobase; MW = molecular weight.

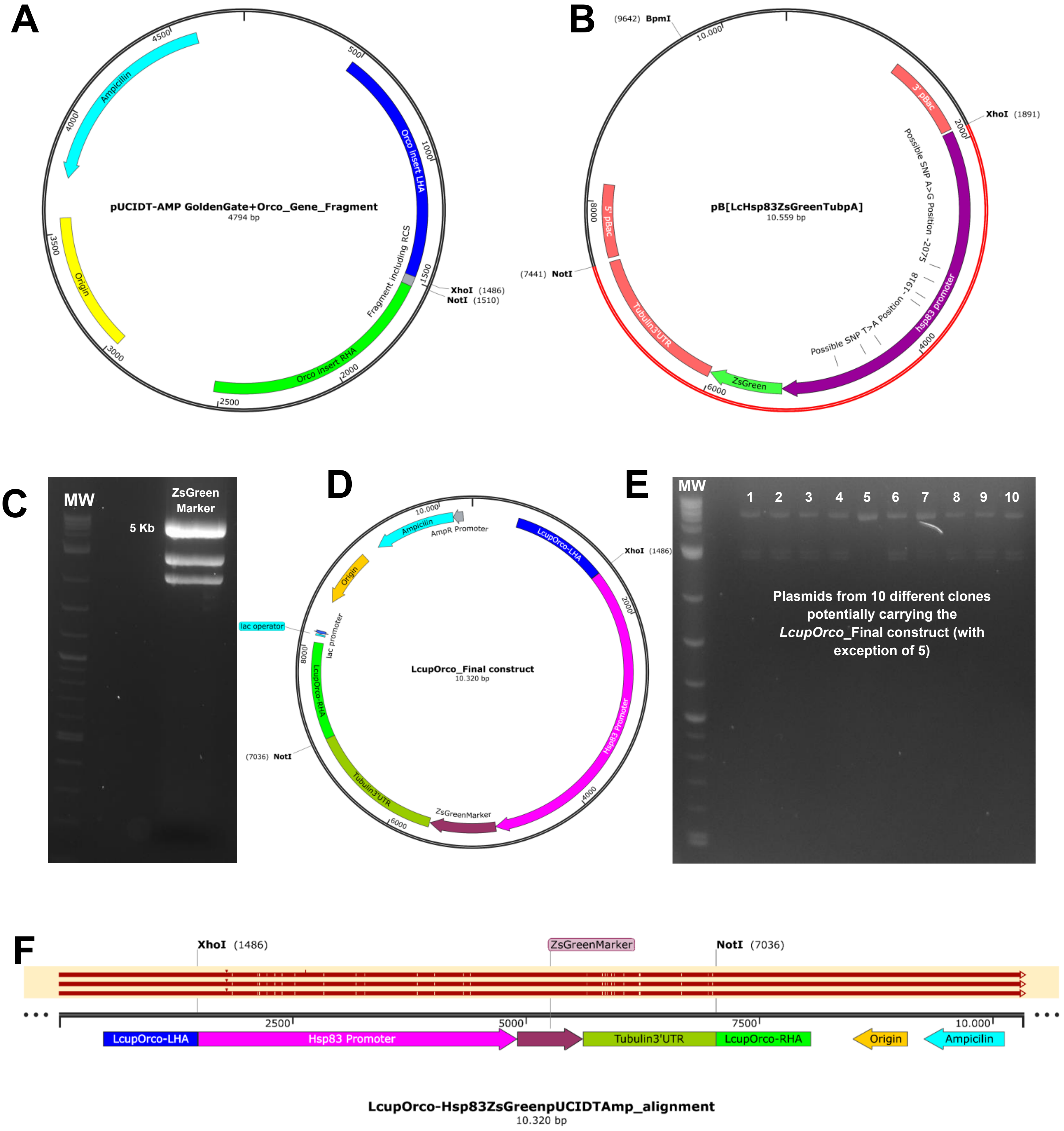

**Supplementary Figure 11: containers used for fly rearing and crosses.** (**A**) plastic bottle used for fly rearing; (**B**) sandglass shaped arrangement made using two five ounces clear-plastic cups joined from removed bottoms used for fly crosses after embryo injections. These containers also included: a top holed cap and a bottom cap with an opening of ~ 1 inch in diameter; a rounded piece of white paper towel; a small glass vial filled with tap water and a small protein cookie made with yeast, milk and egg powder and cane sugar.

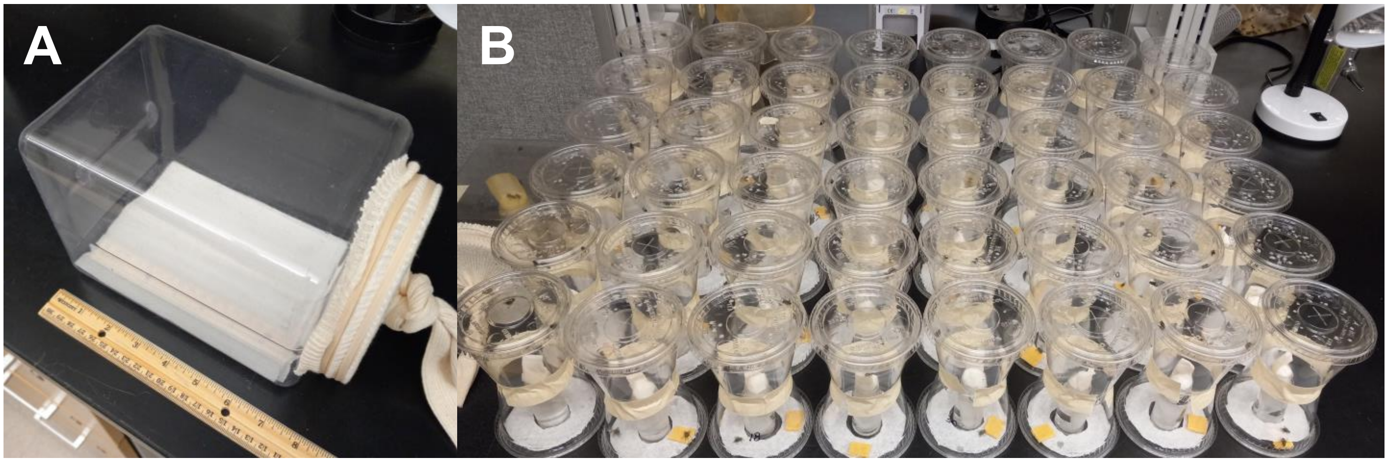

**Supplementary Figure 12: fly crosses completed after embryos microinjections.** Abbreviations: G = generation; *wt* = wild-type.

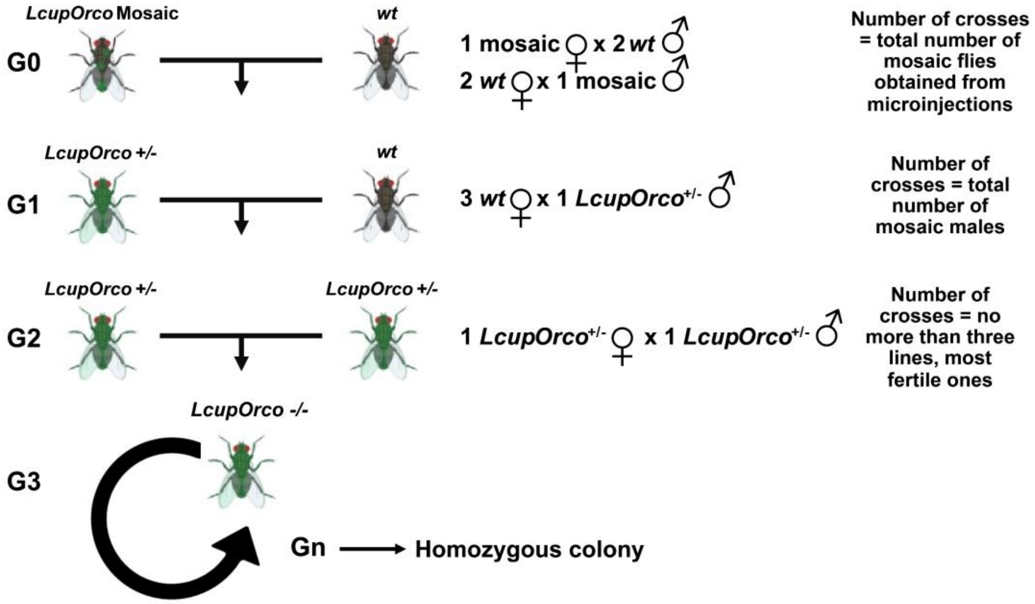

**Supplementary Note 2: primers used for *LcupOrco* insert detection of landing site.** Three types of pair of primer associated with three different strategies were used to determine if the landing site of the *LcupOrco* insert was withing the *LcupOrco* locus in the *L. cuprina* genome. First one was associated with the left side (LS) of the insert including the upstream genome region, the *Hsp83* promoter and part of the green marker; the second one was designed to amplify the right side (RS) of the insert including the part of Tub3'-UTR and the downstream genome region; and third one was to amplify the whole *LcupOrco* insert (All) including from upstream to downstream *L. cuprina* genome regions flanking the *LcupOrco* locus.

Lcup-Orco-Genot-LS_Fw1: GCGAATACATTCTCCAAAGATTTCC
Lcup-Orco-Genot-LS_Rv1: GTTCAGTTCTAGTTCGGTTCTAGTT
Lcup-Orco-Genot-LS_Fw2: TGATTAGCGAATACATTCTCCAAAGATTTC
Lcup-Orco-Genot-LS_Rv2: GTTCGGTTCTAGTTCAGTTCTAGTTTAGT
Lcup-Orco-Genot-RS_Fw1: TGTATCATGGGACGCTAAGATCAA
Lcup-Orco-Genot-RS_Rv1: CTCTGACACAAACTTGGGTTCAAT
Lcup-Orco-Genot-RS_Fw2: TCTACCACGATGATCTCATTCACA
Lcup-Orco-Genot-RS_Rv2: TGCGTACCATCATTTCCTGTTTC
Lcup-Orco-Genot-RS_Fw3: TATTGTATCTACCACGATGATCTCATTCAC
Lcup-Orco-Genot-RS_Rv3: GCTACTTAGTTAGAATGGATGGAGTTAGTT
Lcup-Orco-Genot-All_FW1: ACTAGTTTAATGGAATGCGTTTCTC
Lcup-Orco-Genot-All_RV1: CTGACACAAACTTGGGTTCAATAA
Lcup-Orco-Genot-All_FW2: GCGAATACATTCTCCAAAGATTTCC

Lcup-Orco-Genot-All_RV2: AGTTAGTTGGGTCGCTAACTTATTT

**PCR cycling parameters for LS and RS primers, using Q5 Taq (NEB)**

Initial denaturalization = 3 min at 98°C.

Denaturalization = 30 sec at 98°C.
Annealing = 30 sec at 64°C. Repeat x 40 cycles
Extension = 2 min at 72°C.

Final extension = 5 min at 72°C.
Hold = ∞ at 10°C.

**PCR cycling parameters for primers to amplify the whole insert, using LongAmp Taq (NEB)**

Initial denaturalization = 3 min at 94°C.

Denaturalization = 30 sec at 94°C.
Annealing = 30 sec at 64°C. Repeat x 30 cycles
Extension = 7 min at 65°C.

Final extension = 10 min at 65°C.
Hold = ∞ at 10°C.

Reaction was performed in 20 µl of final volume and 300 nM for final primer concentration. For LS and RS amplification, the Q5 High-Fidelity Taq Pol was used, and for the whole insert amplification the LongAmp® Taq DNA Polymerase (NEB, Cat. # M0323S).
